# Tom70-mediated mitochondria–nuclear envelope contacts regulate nuclear pore complex inheritance during gametogenesis

**DOI:** 10.1101/2025.11.20.689567

**Authors:** Cyrus T. Ruediger, Benjamin S. Styler, Eric M. Sawyer, Silvan Spiri, Grant King, Madison E. Walsh, Gloria A. Brar, Danielle M. Jorgens, Elçin Ünal

## Abstract

Gametogenesis rejuvenates the cellular lineage and excludes senescence-associated factors from gametes. In *Saccharomyces cerevisiae*, this involves sequestration of nuclear constituents into the Gametogenesis-Uninherited Nuclear Compartment (GUNC), which is excluded from gametes. Here we identify the conserved mitochondrial import receptor Tom70 as a key regulator of GUNC-mediated exclusion. Loss of *TOM70* disrupts the sequestration of nuclear pore complexes, but not senescence-associated aggregates and nucleolar components, into the GUNC. Tom70’s role appears independent of its canonical function in mitochondrial import and instead reflects a meiosis-specific requirement for mitochondria-nuclear envelope tethering. During meiosis II, Tom70 concentrates around the GUNC, where it recruits the nuclear envelope tethering protein Cnm1. Loss of *CNM1* partially phenocopies *tom70Δ*, consistent with parallel tethering interactions. These findings uncover a previously unrecognized organelle contact-dependent pathway that remodels the nuclear envelope to support selective nuclear inheritance. More broadly, they highlight how organelle contacts integrate with nuclear quality control to safeguard gamete integrity.

## Main

Gametogenesis is a specialized developmental program that generates reproductive cells through the conserved process of meiosis. In addition to ensuring faithful chromosome segregation, gametogenesis deploys multiple endogenous mechanisms that prevent the inheritance of cellular damage, including senescence-associated factors^1–4^. Elucidating the molecular logic of these pathways provides fundamental insight into how organisms preserve lineage fidelity across generations and informs our understanding of aging and germline health.

The budding yeast *Saccharomyces cerevisiae* is a powerful system for dissecting conserved molecular mechanisms during gametogenesis. Diploid yeast can synchronously undergo gametogenesis (sporulation), enabling real-time observation of nuclear and organelle dynamics^5,6^. Moreover, yeast has long served as a valuable model for studying mechanisms of aging: like multicellular organisms, it has a finite replicative lifespan and displays conserved age-associated hallmarks^7–11^. Remarkably, aged cells generate gametes with fully restored lifespan, highlighting the rejuvenating potential of gametogenesis^12^.

Meiosis is accompanied by striking, developmentally programmed remodeling of organelles, most prominently during meiosis II. At this stage, *de novo* gamete plasma membranes (also known as prospore membranes; PSMs) are synthesized^6,13^, while the endoplasmic reticulum (ER), mitochondria, and nucleus undergo large-scale reorganization^2,3,14,15^. The cortical ER retracts and reorganizes around the nucleus^15,16^, while mitochondria simultaneously detach from the cell cortex and relocalize to the nuclear periphery (Fig. 1a)^15,17–20^. Although temporally coordinated by a common transcriptional program, these events are mediated by distinct molecular mechanisms.

**Figure 1:**
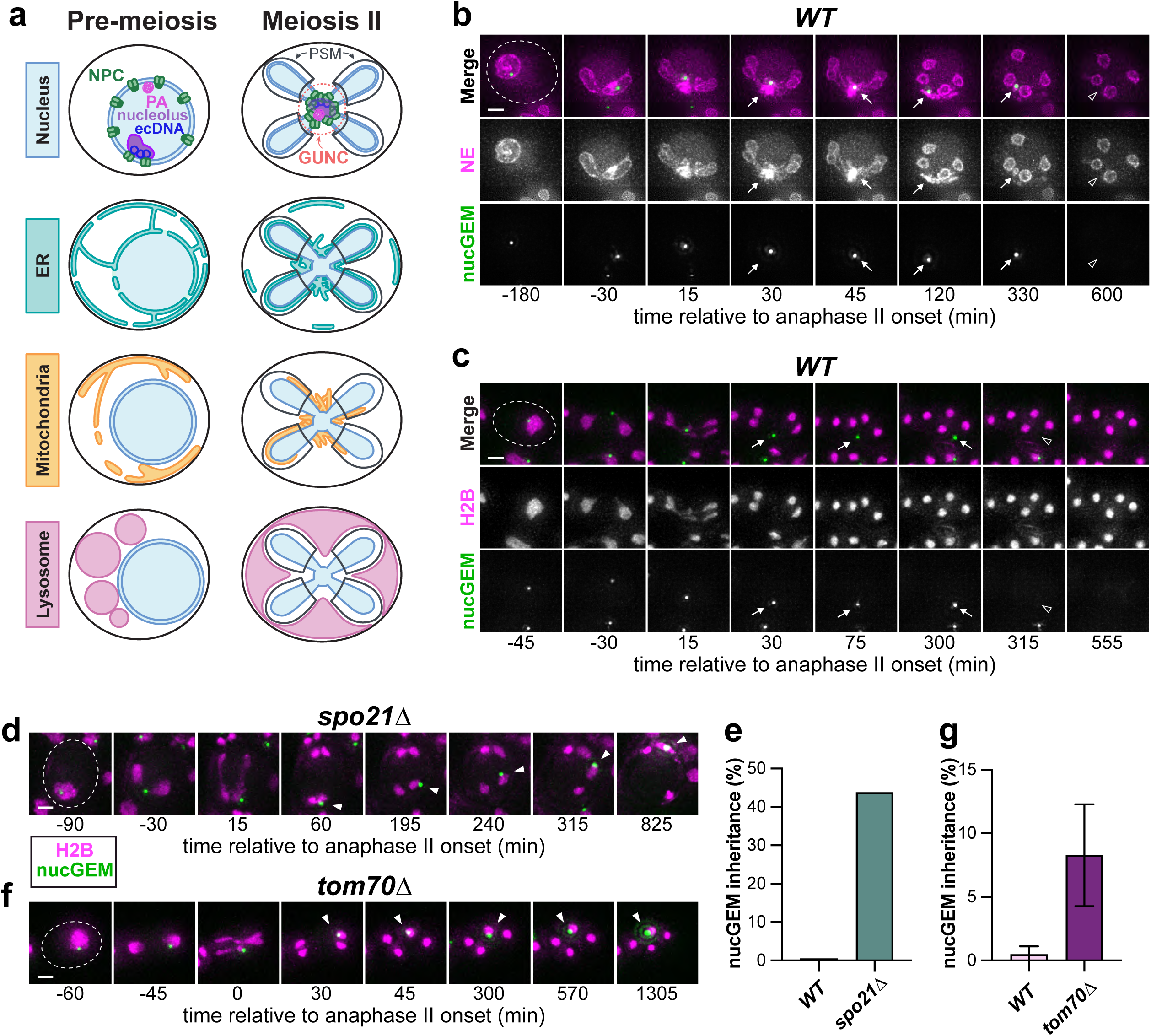
Genome-wide screen identifies Tom70 as a key regulator of nuclear protein clearance during gametogenesis. **a,** Schematic of organelle remodeling during budding yeast meiosis II, modified from ^3^. Nuclear pore complexes (NPC) and senescence-associated factors (protein aggregate, PA; extrachromosomal rDNA circles, ecDNA) are confined to a sequestered membrane-bound nuclear compartment (GUNC). The endoplasmic reticulum (ER) and mitochondrial network detach from the cell periphery and associate with the GUNC region. The vacuole/lysosome-equivalent fuses and expands during meiosis II. Vacuolar lysis subsequently releases hydrolases which degrade excluded senescence factors and organelles. **b,** Montage of a wild-type cell expressing a fluorescently-tagged nuclear particle (nucGEM; NLS-PfV-mTSapphire) and nuclear envelope marker (NE; 3xmScarlet-Heh2^93–378^) progressing through meiosis and gamete maturation. Arrows denote GUNC region, open arrowhead emphasizes nucGEM degradation upon vacuolar lysis. Time is shown in minutes relative to onset of anaphase II (0 min). **c,** Montage of nucGEM sequestration during meiosis relative to chromatin (H2B; histone Htb1-mCherry). Arrows indicate GUNC region, open arrowhead emphasizes nucGEM degradation. **d, f** Montage of nucGEM relative to chromatin during meiosis in a (d) *spo21Δ* or (f) a *tom70Δ* mutant. Arrowheads emphasize segregation alongside chromatin and inheritance (or persistence) in gametes. *spo21Δ* mutants, which do not form viable gametes with plasma membranes, do not undergo vacuolar lysis and often exhibit nuclear rupture or recoalescence following the meiotic division. **e, g** Quantification of nucGEM inheritance for (e) *spo21Δ* and (g) *tom70Δ* mutants. (e) Shown is a single experiment, n = 185-208 cells. (g) Shown is the average ± standard deviation for two experiments, with n = 205-230 cells each per genotype; p = 0.10, two-tailed unpaired t-test. Scale bars, 2 µm.

Concurrently in meiosis II, the nucleus undergoes an unconventional five-way division that generates four gamete nuclei and an additional membrane-bound compartment, termed the Gametogenesis Uninherited Nuclear Compartment (GUNC)^21,22^. The GUNC sequesters long-lived nuclear constituents, such as nuclear pore complexes (NPCs) and nucleolar proteins enriched in intrinsically disordered, aggregation-prone domains^23,24^. The GUNC also sequesters senescence-associated factors, including misfolded, chaperone-bound proteins (e.g., Hsp104 substrates) and extrachromosomal circular DNA (ecDNA) from the rDNA locus together with associated nucleolar proteins^25–29^. Remarkably, regardless of cellular age, collapsed ER and mitochondria accumulate around the GUNC to form a transient super-organellar structure that is excluded from gametes^21^. Following PSM closure and gamete cellularization, vacuolar/lysosomal membrane rupture executes a large-scale autophagy-like process known as mega-autophagy, which drives bulk degradation of the GUNC and its constituents, as well as other organelles left outside spore wall-protected gametes (Fig. 1a) ^21,30,31^.

To date, only a single factor, Spo21, has been implicated in GUNC-mediated exclusion^21^. Spo21 promotes PSM expansion^32^, a process essential for the encapsulation of gamete nuclei and other organelles. In *spo21Δ* mutants, as well as in other contexts where PSM formation is disrupted, the GUNC fails to form^21,33^. As a result, nuclear constituents normally destined for exclusion remain diffusely distributed throughout the meiosis II nucleus^21^. In the absence of PSM encapsulation, all nuclear and cytoplasmic material is ultimately degraded during the wave of mega-autophagy that follows meiosis^31^, thereby precluding downstream analysis of gametes. These observations raise several key questions: Beyond PSM assembly, do additional pathways regulate GUNC-mediated exclusion? Are diverse GUNC constituents excluded from gametes through a shared mechanism or through distinct processes? Do coincident organelle remodeling events, such as ER collapse and mitochondrial redistribution, contribute to this exclusion? And what are the consequences of disrupting GUNC-mediated exclusion for gamete quality and fitness?

Here we begin to address these questions by uncovering a novel mechanism that regulates nuclear compartmentalization during gametogenesis. Through an unbiased genome-wide screen, we identify the conserved mitochondrial import receptor Tom70 as a key regulator of GUNC-mediated exclusion and a crucial determinant of gamete health. Remarkably, Tom70’s role in this context is independent of its canonical function in mitochondrial protein import and appears unrelated to PSM formation. Instead, we find that Tom70 forms prominent mitochondria-nuclear envelope contact sites near the GUNC, stabilized in part by the meiosis-specific nuclear envelope protein Cnm1. These organelle contacts contribute to the formation of a lateral diffusion barrier that restricts nuclear pore complex inheritance into nascent gamete nuclei. This role is specific, as Tom70 is dispensable for the exclusion of senescence-associated aggregates and nucleolar components, indicating that individual GUNC constituents are sorted through distinct pathways during gametogenesis. Together, our findings uncover a developmentally regulated organelle contact-dependent mechanism that remodels the nuclear envelope to enforce selective nuclear inheritance.

### Tom70 regulates nucGEM inheritance

To identify new factors that regulate GUNC-mediated exclusion during gametogenesis, we sought a tractable nuclear marker amenable to genome-wide screening. During this effort, we serendipitously found that meiotic overexpression of fluorescently-tagged nuclear-localized virus-like particles (nucGEM, nuclear Genetically Encoded Multimeric nanoparticle^34,35^) generated a single, bright nuclear focus that was reliably partitioned into the GUNC (Fig. 1b, arrows; and Supplementary Video 1) and degraded later in meiosis (Fig. 1b, open arrowheads). Importantly, nucGEM behavior closely resembled that of Hsp104-marked protein aggregates in aged cells–both were robustly partitioned into the GUNC in wild-type meiosis (Fig. 1c, arrowheads; Supplementary Video 2), but mis-localized to chromosomes in *spo21Δ* mutants (Fig. 1d,e; arrowheads)^21^. These properties made nucGEMs a robust proxy for nuclear aggregates and a powerful, age-independent tool for probing GUNC-mediated exclusion. Using a high-content imaging platform, we screened the yeast non-essential gene deletion collection in the highly sporulation-proficient SK1 strain background and identified mutants defective in clearing nucGEM foci during meiosis.

The complete dataset, screen design, validation, and mechanistic analyses will be reported in a future publication. Here, we focus on one particularly striking candidate identified among the top hits. This gene, *TOM70*, encodes a well-studied outer mitochondrial import receptor, yet its loss impacts meiotic nuclear compartmentalization and gamete health, raising implications of a novel function for organelle remodeling and inter-organelle crosstalk during gametogenesis.

To better understand how Tom70 contributes to nuclear protein clearance during gametogenesis, we performed time-lapse microscopy and monitored nucGEM foci relative to chromosomes throughout meiosis. In wild-type cells, nucGEM foci were efficiently excluded and rarely inherited by gametes (0.5% inheritance, Fig. 1c,g; Supplementary Video 3). Instead, the foci segregated away from dividing chromosomes during meiosis II, consistent with their sequestration into the GUNC. In contrast to wild type, 8.3% of *tom70Δ* progenitor cells failed to exclude the nucGEM, resulting in its inheritance by one of the gametes (Fig. 1f,g). This phenotype is indicative of a defect in GUNC-mediated exclusion, rather than a failure in mega-autophagy in *tom70Δ* mutants.

### Tom70 restricts the inheritance of nuclear pore complexes into gametes

Given Tom70’s apparent role in excluding nucGEM from gamete nuclei, we next asked whether Tom70 similarly impacts the inheritance of endogenous nuclear proteins that are normally excluded via the GUNC. To address this question, we examined NPCs, which are sequestered into the GUNC during meiosis, irrespective of progenitor cell age. We focused on two core nucleoporins: Nup170-GFP, a component of the NPC inner ring, and Pom34-GFP, a transmembrane nucleoporin, alongside Htb1-mCherry (histone H2B) to mark chromatin^21^. Time-lapse microscopy in wild-type cells revealed robust exclusion of both Nup170 and Pom34 from gamete nuclei (Fig. 2a and 2b, arrows; Supplementary Videos 4 & 5), with only minimal signal surrounding mature nuclei (Fig. 2a and 2b, arrowheads). In contrast, *tom70Δ* mutants exhibited defective NPC sequestration (Fig. 2a and 2b, arrows; Supplementary Videos 6 & 7), resulting in increased inheritance of both Nup170 and Pom34 by gamete nuclei (Fig. 2a and 2b, arrowheads). This abnormal NPC sequestration was apparent in approximately 70% of *tom70Δ* mutant cells (Fig. 2c and 2d; for representative images of each category, see Extended Data Fig. 2a and Supplementary Video 8). These defects were meiosis-specific, since auxin-induced depletion of Tom70 exclusively just prior to meiosis using the AID2 system^36,37^ recapitulated the Nup170 mis-sequestration phenotype (Extended Data Fig. 2b,c).

**Figure 2:**
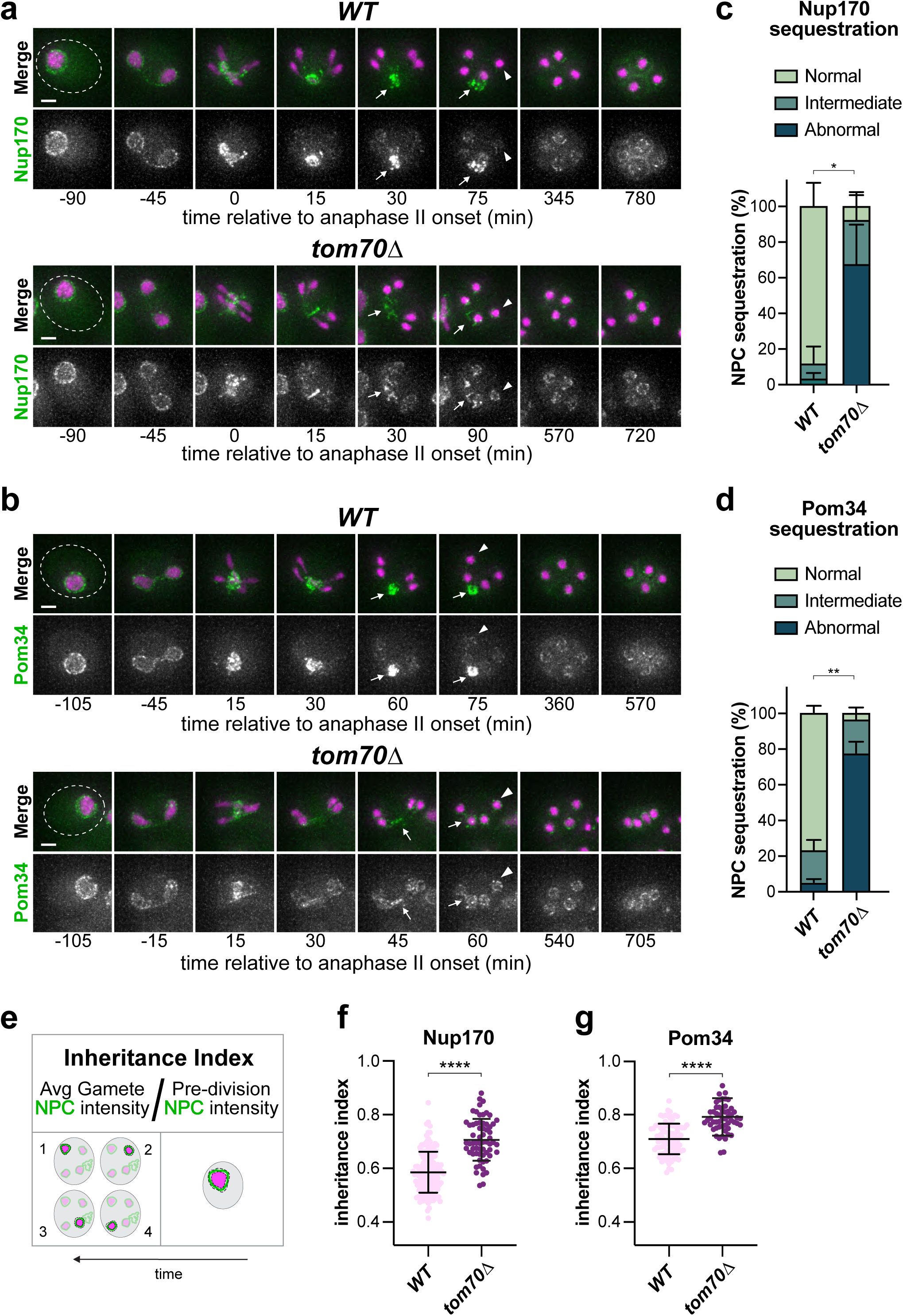
Tom70 restricts the inheritance of nuclear pore complexes into gametes. **a, b** Montages of (a) inner ring nucleoporin Nup170-GFP or (b) transmembrane nucleoporin Pom34-GFP sequestration relative to chromatin (Htb1-mCherry, magenta) in sporulating wild-type (*WT)* cells and *tom70Δ* mutants. Arrows indicate nucleoporin signal accumulation at the GUNC, arrowheads highlight the nucleoporin signal at spore nuclei upon PSM closure. **c, d** Quantified penetrance of abnormal nucleoporin (NPC) sequestration in (a) and (b), respectively (see Extended Data Fig. 2a for representative images of category). Shown is an average ± standard deviation from two experiments, (c) n = 89-153 and (d) n = 58-113 cells per genotype in each. ** (p<0.01), * (p<0.05), two-tailed unpaired t-test of percent abnormal sequestration. **e,** Schematic for NPC inheritance index calculation using single-cell measurement of NPC signal at progenitor and gamete nuclear peripheries (average of four gametes/progenitor). Magenta is chromatin signal, green is nucleoporin signal, faded structures represent distinct focal planes, and dashed black lines indicate ROI boundaries. See Methods for additional detail. **f, g** Quantified inheritance index for (f) Nup170-GFP and (g) Pom34-GFP in individual wild-type and *tom70Δ* cells. Shown the average ± standard deviation from a single experiment, (f) n = 63-150 and (g) n = 95-54 progenitor cells per genotype each. Only cells forming four closed spores that remained intact through the entire 18h movie were quantified. **** (p<0.0001), Kolmogorov-Smirnov test. Scale bars, 2 µm.

To assess the degree of NPC inheritance, we developed a nucleoporin inheritance index to quantify NPC inheritance at the single-cell level (Fig. 2e, see Methods for further detail). This analysis revealed a significant increase in NPC inheritance in *tom70Δ* compared to wild-type cells (Fig. 2f and 2g). Similar defects were observed when tracking the channel nucleoporin Nup49 (Extended Data Fig. 2d), indicating that missegregation is not specific to Nup170 or Pom34, but likely extends to other core NPC components. By contrast, Nup2, a nuclear basket nucleoporin that is not normally excluded via the GUNC^21,38^, behaved similarly in wild-type and *tom70Δ* cells (Extended Data Fig. 2e), supporting the specificity of Tom70’s role in regulating the inheritance of core nucleoporins. Together, these findings demonstrate that Tom70 is required for the GUNC-mediated exclusion of endogenous nuclear proteins, including multiple subunits of the nuclear pore complex.

### Tom70 is dispensable for the gamete exclusion of senescence-associated nuclear factors, revealing selectivity of sequestration pathways for distinct substrates

In addition to core nucleoporins, several nuclear senescence-associated factors are normally confined to the GUNC, thereby preventing their inheritance by gametes. These include nuclear protein aggregates, ecDNA, and excess nucleolar material (Fig. 3a)^3,21^. To determine whether the nuclear sequestration defects observed in *tom70Δ* mutants extend to these age-associated accumulations, we performed time-lapse microscopy on replicatively aged cells, enriched as previously described^21,39,40^.

**Figure 3:**
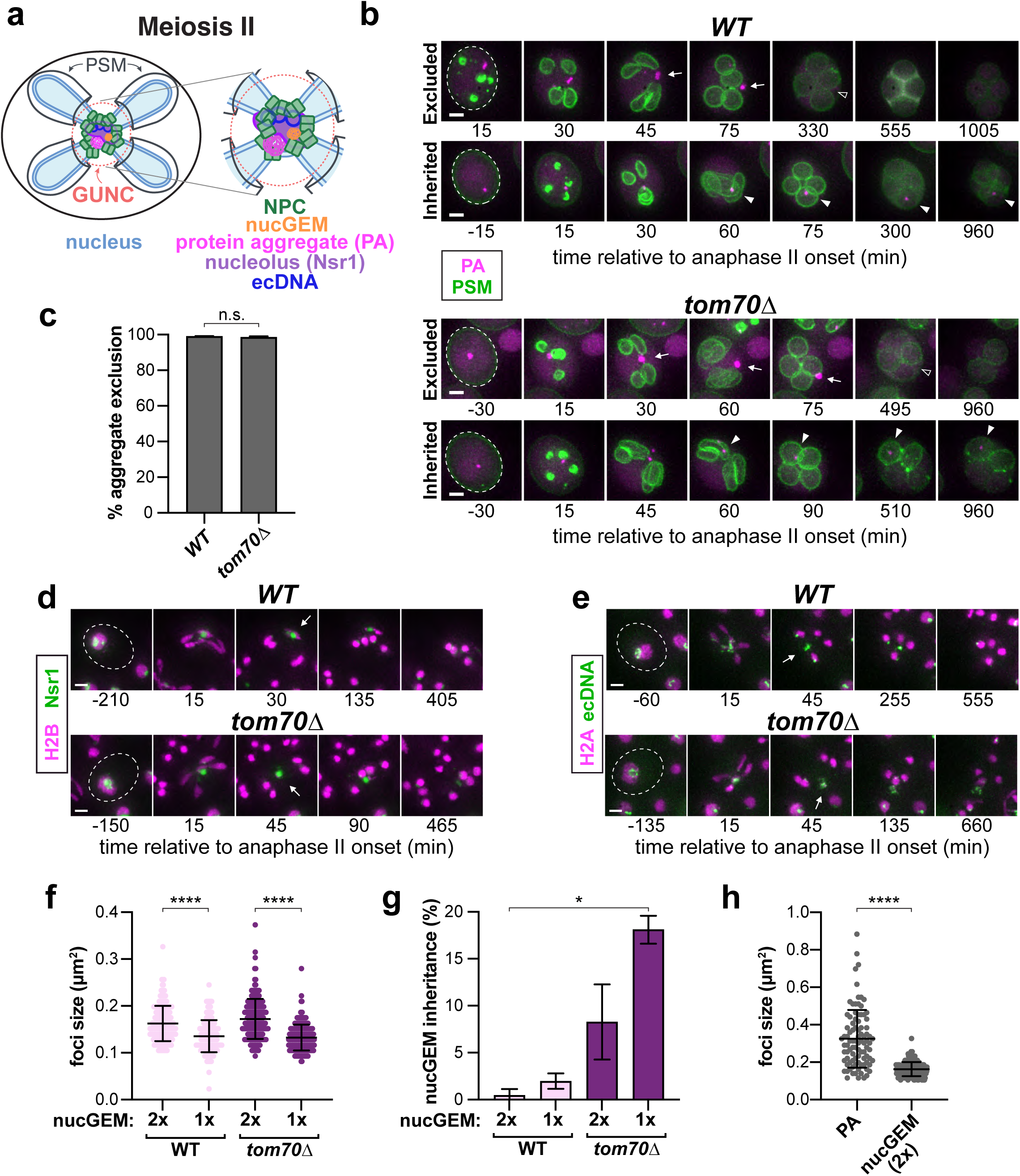
Tom70 is dispensable for the gamete exclusion of senescence-associated nuclear factors, revealing size-dependent inheritance limits. **a,** Schematic of GUNC-sequestered factors, including previously examined nucGEM and NPCs, as well as age-associated protein aggregates (PA), excess nucleolar material, and extrachromosomal rDNA circles (modified from ^3^). **b,** Montages of replicatively-aged wild-type and *tom70Δ* mutant cells either excluding or inheriting protein aggregates, marked with Hsp104-mCherry. Gamete plasma membranes (PSM), shown in green, are marked by GFP-Spo21^51–91^ overexpressed from a *P_ATG8_* promoter. Cells were aged by pulse biotin labeling followed by sorting after outgrowth. Time is shown in minutes relative to onset of anaphase II (0 min), inferred by the coincident nucleation of PSMs. **c,** Quantification of age-induced protein aggregate inheritance, as seen in (a). Aged cells were mixed 1:1 with young cells to promote efficient sporulation (see Methods), and inheritance of any non-transient protein aggregate inheritance was scored for cells of mixed age. Shown is average ± standard deviation from two biological replicates, n = 187-420 per genotype each. p = 0.76, two-tailed Mann-Whitney U test. **d, e** Montage of replicatively-aged wild type and *tom70Δ* mutants tracking chromatin (Htb1-mCherry, H2B; or Hta1-mApple, H2A) and either (d) Nsr1-GFP-marked enlarged nucleoli, or (e) TetR-GFP-marked extrachromosomal rDNA circles (recognized via a heterozygous rDNA-5xTetO repeat). Arrows indicate GUNC region. Note: two gametes always inherit a TetR-GFP dot, which represents the copy of the rDNA-5xTetO array at the chromosomal locus. **(f)** Quantification of GEM particle foci size (mid-anaphase II) upon decreased expression via reduction in transgene dosage, from homozygous (2x) to heterozygous (1x), in both wild type and *tom70Δ* mutant single cells. Bars indicate the average ± standard deviation from a single experiment, n = 168-231 cells per genotype. **** (p<0.0001), Kolmogorov-Smirnov test. nucGEM particle foci size does not differ between wild type and *tom70Δ*: 2x (p = 0.23), 1x (p = 0.52). **g,** Quantification of nucGEM inheritance (as in Fig. 1), with either homozygous or heterozygous overexpression from a *P_ATG8_* promoter, in *WT* and *tom70Δ* mutants. Shown is an average ± standard deviation from two independent experiments, n = 185-220 per genotype each. Homozygous data is replotted from Fig. 1g. **h,** Quantification of protein aggregate (PA; Hsp104-GFP) foci size mid-anaphase II, relative to homozygous (2x) nucGEM particles; replotted from (f); 2x, *WT*. Similar to (c), protein aggregate sizes were quantified from a mix of aged and young cells. Bars show the average ± standard deviation from a single experiment, n = 92 cells. **** (p<0.0001), Kolmogorov-Smirnov test. Scale bars, 2 µm.

In wild-type and *tom70Δ* aged cells, Hsp104-mCherry-marked protein aggregates (PA) that form near the nuclear periphery were efficiently sequestered from gametes (Fig. 3b,c). Similarly, the enlarged nucleoli and ecDNA characteristic of old cells, marked by Nsr1-GFP or TetR-GFP, respectively, were excluded from gamete nuclei in *tom70Δ* mutants to a comparable extent as in wild type (Fig. 3d,e). These results suggest that the sequestration defect in *tom70Δ* cells may be selective, primarily affecting NPCs while leaving larger or more static nuclear structures unaffected. Alternatively, distinct exclusion pathways may operate on different nuclear substrates. Importantly, these findings reveal that individual GUNC constituents can exhibit divergent fates during gametogenesis.

### Differential nuclear protein inheritance in *tom70Δ* gametes may reflect size-dependent exclusion limits

To directly assess whether size influences GUNC-mediated sequestration and exclusion, we took advantage of the nucGEM reporter. To modulate nucGEM size, we altered the dosage of the nucGEM transgene. Diploid cells carrying either one or two copies of the reporter, driven by the promoter *P_ATG8_* which exhibits strong expression in meiosis, were analyzed by time-lapse microscopy. We observed a clear correlation between approximate particle size and inheritance. Cells carrying two copies contained larger nucGEM foci (Fig. 3f, compare 2x to 1x), which were more efficiently excluded from gamete nuclei (Fig. 3g, compare 2x to 1x). In *tom70Δ* mutants, this relationship persisted, and the sequestration defect was most pronounced for the smallest nucGEM particles (Fig. 3g). These findings suggest that the differential inheritance of nuclear substrates, such as nucGEM and NPC components versus senescence-associated factors, may in part stem from inherent size differences. Consistently, age-associated protein aggregates were significantly larger than nucGEM foci (Fig. 3h).

### Tom70 is necessary for gamete health but not rejuvenation

Having established that Tom70 regulates nuclear inheritance, we next investigated how its loss affects overall gamete health and fitness. To this end, we evaluated multiple metrics–meiotic progression, efficiency of spore production, gamete viability, mitotic cell cycle re-entry kinetics, spore colony size, and replicative lifespan resetting–in two distinct *tom70* mutant backgrounds: (1) *tom70Δ*, in which the entire *TOM70* coding sequence is deleted and (2) *P_tetO_-TOM70*, in which *TOM70* mRNA expression is constitutively repressed unless induced by addition of anhydrotetracycline (aTc)^41^.

Although *tom70Δ* mutants completed meiotic divisions with kinetics comparable to wild type and produced viable gametes, these gametes exhibited severe fitness defects, as evidenced by delayed cell cycle re-entry and significantly smaller spore colony size (Fig. 4a-c, Extended Data Fig. 4a). The number of gametes produced per cell was also modestly reduced (Extended Data Fig. 4b). These phenotypes were specific to meiosis, since *tom70Δ* mutants displayed no significant growth defects relative to wild type during vegetative proliferation (Extended Data Fig. 4c). Consistent with this, meiosis-specific expression of *TOM70* in the *P_tetO_-TOM70* strain fully rescued gamete quality (Fig. 4d, Extended Data Fig. 4d).

**Figure 4:**
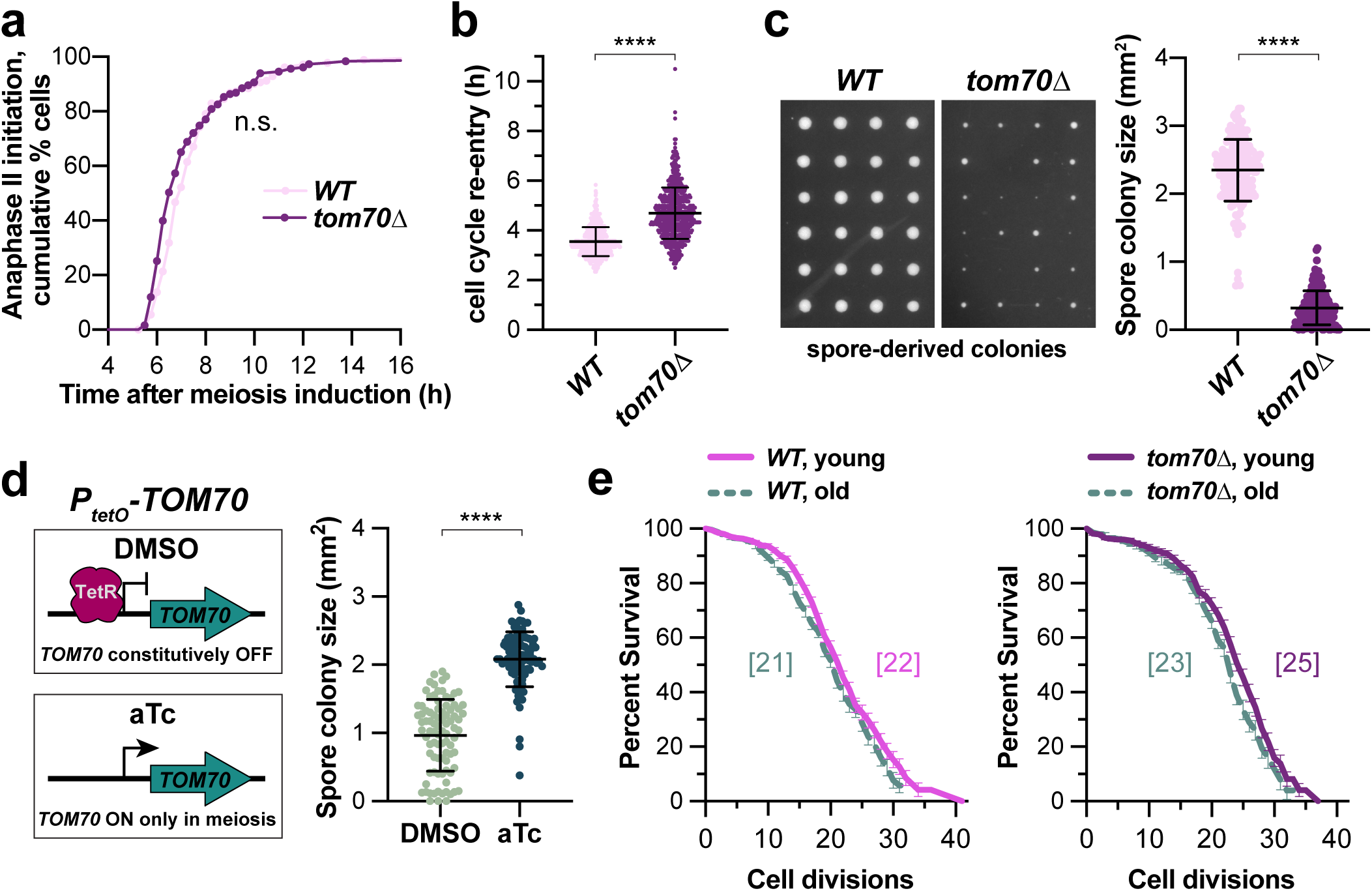
Tom70 is necessary for gamete health but not rejuvenation. **a,** Quantification of cumulative percent cells initiating anaphase II over time, assessed by timelapse microscopy of chromatin (Htb1-mCherry). Shown is representative of two experiments, n = 181-182 cells per genotype. p = 0.16, logrank Mantel-Cox test. **b,** Quantification of gamete cell cycle re-entry upon addition of rich media (YPD) at 30°C, assessed by initial emergence of first daughter bud. Times are shown for individual cells from 3 pooled biological replicates, n = 169-207 per genotype per replicate. Bars indicate average ± standard deviation. **** (p<0.0001), Kolmogorov-Smirnov test. c, (left) Representative colony growth of individual gamete cells (prepared after 24h in SPO), after 48h of subsequent growth on rich media (YPD) at 30°C. (right) Individual colony sizes from two independent experiments are plotted, n = 86-95 cells per genotype per experiment. Bars indicate average ± standard deviation. **** (p<0.0001), Kolmogorov-Smirnov test. **d,** (left) Schematic for conditional expression via the *P_tetO_* promoter; *TOM70* is only expressed upon addition of anhydrotetracycline (aTc). (right) Quantification of gamete colony sizes following addition of either 5 µg/mL aTc or vehicle control upon induction of meiosis (0h in SPO media). See Extended Data Fig. 4d for protein level comparison to wildtype prior to initiation of meiotic divisions (6h in SPO media). Shown is a single experiment, bars indicate average ± standard deviation, n = 92 cells per condition. **** (p<0.0001), Kolmogorov-Smirnov test. **e,** Replicative lifespan was determined for *WT* or *tom70Δ* gametes derived from either young or replicatively-aged progenitor cells, as previously described^42^. Shown is the Kaplan-Meier plot of two or more pooled replicates ± standard error of the mean, n = 190-315 cells total per genotype. *WT* (p = 0.016), tom70Δ (p = 0.005); logrank Mantel-Cox test.

Despite these impairments in gamete quality, the ability to reset replicative lifespan following gametogenesis remained intact in *tom70Δ* gametes, as measured using our recently described microfluidics-based pedigree assay (Fig. 4e, Extended Data Fig. 4f)^42^. This aligns with our earlier findings that *TOM70* is dispensable for GUNC-mediated clearance of senescence-associated nuclear factors, including ecDNA and protein aggregates. Together, these findings uncover a surprising uncoupling between gamete health, defined by growth potential and cell cycle re-entry efficiency, and gametogenic rejuvenation, defined by replicative lifespan resetting. This suggests that the mechanisms that restore gamete replicative potential can operate independently of those that ensure gamete fitness.

### Mitochondrial import is dispensable for nuclear sequestration and GUNC-mediated exclusion

Tom70 is a single-pass outer mitochondrial transmembrane protein, which primarily facilitates the recognition and import of a specific subset of mitochondrial proteins from the cytosol. It acts as a docking site for mitochondrial targeting signals (MTSs) and cytosolic chaperones alongside their hydrophobic cargo, helping maintain solubility and guiding them to the TOM translocon for insertion^43–45^. To determine whether disruption of mitochondrial import impacts nuclear remodeling, we employed multiple strategies to perturb distinct components of the import machinery. However, these interventions failed to phenocopy the NPC sequestration defects observed in *tom70Δ* mutants. Specifically, we found deletion of *SAM37* (a β-barrel insertase subunit of the SAM/TOM complex^46^) and auxin-induced depletion of Tom20 (an outer membrane import receptor required for import of mitochondrial substrates^47–49^) each had negligible effects on NPC sequestration and GUNC-mediated exclusion during meiosis (Fig. 5a and 5b). Similarly, chemical uncoupling of mitochondrial membrane potential (thus impeding import into the inner membrane and matrix) with FCCP^50^ had no discernible effect on NPC sequestration (Fig. 5c, Extended Data Fig. 5). These findings indicate that the role of Tom70 in nuclear remodeling is unlikely to stem from a general defect in mitochondrial protein import, but instead reflects a more specialized, import-independent function.

**Figure 5:**
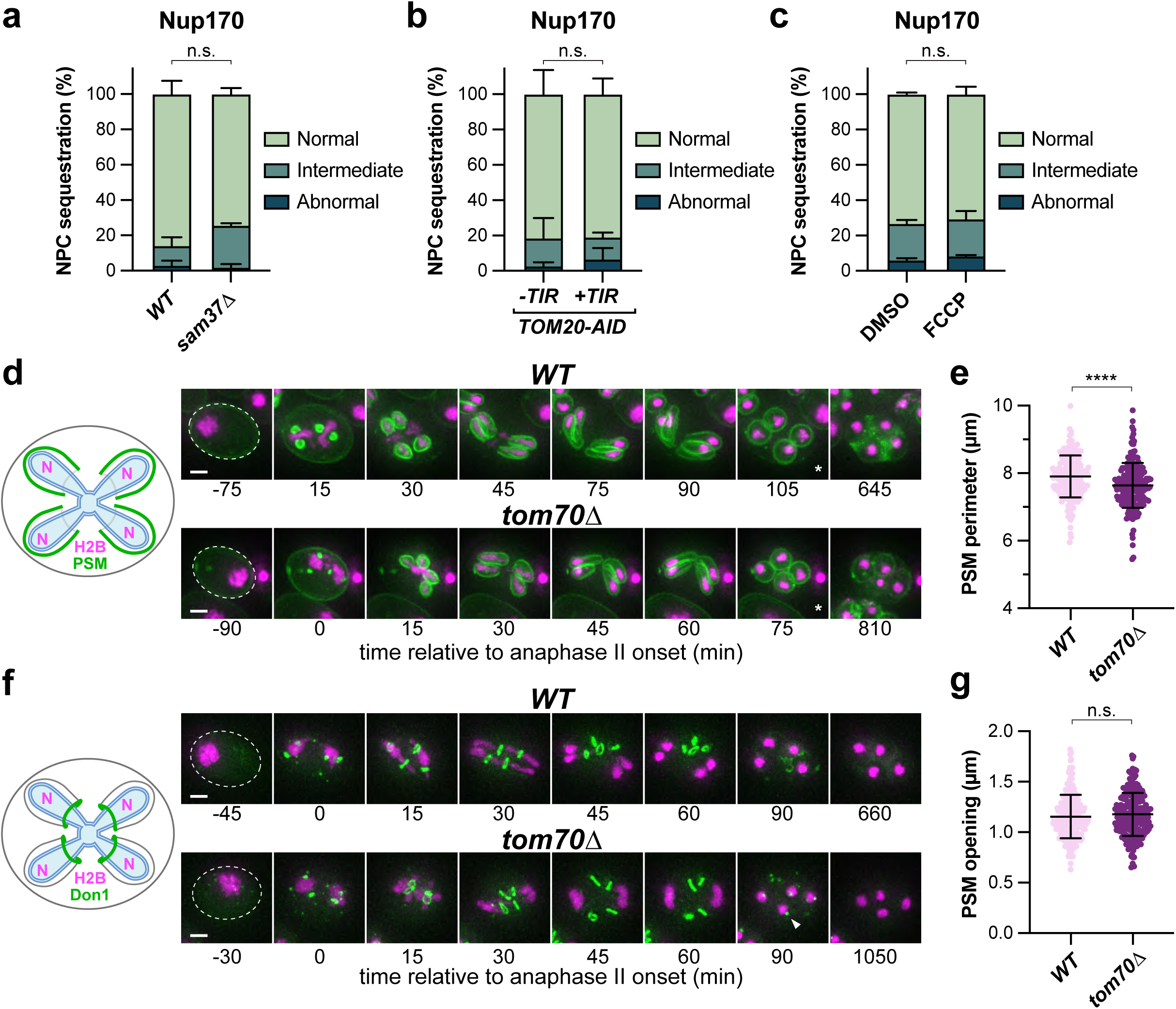
Tom70 promotes NPC sequestration and GUNC-mediated exclusion independently of mitochondrial import and gamete plasma membrane formation. **a-c,** Penetrance of abnormal meiotic nucleoporin sequestration (Nup170-GFP) was quantified for: (a) *sam37Δ* mutants of the SAM/TOB complex, (b) cells depleted of Tom20-AID by addition of 50 µM CuSO_4_ at 2.5h SPO and 1 mM NAA auxin at 4h SPO (see Extended Data Fig. 5a for assessment of depletion), (c) cells treated with mitochondrial uncoupler FCCP (50 µM) at 4h in SPO (see Extended Data Fig. 5b for assessment of mitochondrial depolarization). Shown is an average ± standard deviation from two experiments: (a) n = 43-82 cells, (b) n = 63-102 cells, and (c) n = 190-234 cells per genotype per experiment. (a) p = 0.64, (b) p = 0.49, and (c) p = 0.10; two-tailed unpaired t-test. **d,** Montages of gamete plasma membranes (PSM; see left schematic) marked by *P_ATG8_*-expressed GFP-Spo20^51–91^ and chromatin (H2B; marked by Htb1-mCherry) in sporulating wild-type and *tom70Δ* cells. **e,** Perimeters of individual PSMs were quantified immediately upon closure, assessed by rounding of membranes (see asterisks in d). Only cells forming four closed PSMs which remained intact were considered. Shown is a representative of two experiments, n = 199-201 PSMs per genotype. Bars indicate average ± standard deviation. **** (p<0.0001), Kolmogorov-Smirnov test. **f,** Montages tracking the leading edge complex (marked by Don1-GFP; see left schematic) and chromatin (H2B; marked by Htb1-mCherry) in sporulating wild-type and *tom70Δ* cells. Arrowhead indicates persisting Don1-GFP foci in *tom70Δ* mutants. **g,** Diameters of individual Don1-marked rings were quantified along their longest aspect at near-maximal size, approximately 30 minutes before PSM closure. Only cells forming four closed spores which remained intact were considered. Shown is a representative of two experiments, n = 228-232 Don1-GFP rings per genotype. p = 0.35, Kolmogorov-Smirnov test. Scale bars, 2 µm.

### Tom70 is dispensable for gamete plasma membrane formation

We next examined PSM formation in *tom70Δ* mutants, since PSM formation is crucial for GUNC-mediated exclusion^21^. We assessed two key features: (1) the total size of PSMs and (2) the diameter of their open rims. For the former, we quantified the perimeter of newly completed PSMs using GFP-Spo20^51–91^, a probe that binds phosphatidic acid-rich membranes^51^. For the latter, we measured the diameter of leading-edge complexes (which form ring-like structures at the PSM opening, marked by Don1-GFP^32,52^), at the approximate onset of nucleoporin sequestration defects, 30 minutes prior to PSM closure. *tom70Δ* mutants exhibited a subtle reduction in PSM size, with completed PSMs having roughly 95% the perimeter of wild type, corresponding to 90% of the inferred surface area (Fig. 5d,e). In contrast, PSM opening diameters were indistinguishable between wild type and *tom70Δ* (Fig. 5f,g). Although *tom70Δ* mutants displayed a defect in Don1 clearance (Fig. 5f, arrowhead), its timing –occurring after GUNC-based sequestration defects and PSM closure– is unlikely to account for the nuclear inheritance phenotype. Taken together, these data indicate that PSM formation and morphology are largely intact in *tom70Δ* mutants. Since meiotic nuclear sequestration and GUNC-mediated exclusion have thus far been attributed solely to PSM formation, our findings point to an alternative mechanism by which Tom70 contributes to nuclear remodeling.

### Tom70 is enriched around the GUNC and promotes nuclear envelope remodeling

Our data suggest that Tom70 contributes to nucleoporin sequestration and GUNC-mediated exclusion through mechanisms independent of its established role in mitochondrial protein import and unrelated to any obvious function in gamete plasma membrane formation. This prompted us to explore whether Tom70 might influence exclusion through other means. Tom70 is known to participate in organelle contact sites in both yeast and mammalian cells, clustering at the mitochondrial surface where it can interface with the ER and potentially other membranes^53–56^. This is particularly relevant in meiosis II, when the cortical ER retracts and reorganizes around the nucleus, and mitochondria undergo a dramatic collapse toward the nuclear periphery ^15,16,19^. Time-lapse microscopy of Tom70-GFP revealed that it initially marks mitochondria uniformly at the cell cortex during early meiosis. However, by anaphase II, Tom70-GFP becomes concentrated at GUNC-adjacent mitochondria (Fig. 6a). Co-visualization of mitochondria and the nuclear envelope (NE) confirmed extensive GUNC-proximal mitochondrial enrichment (Fig. 6b).

**Figure 6:**
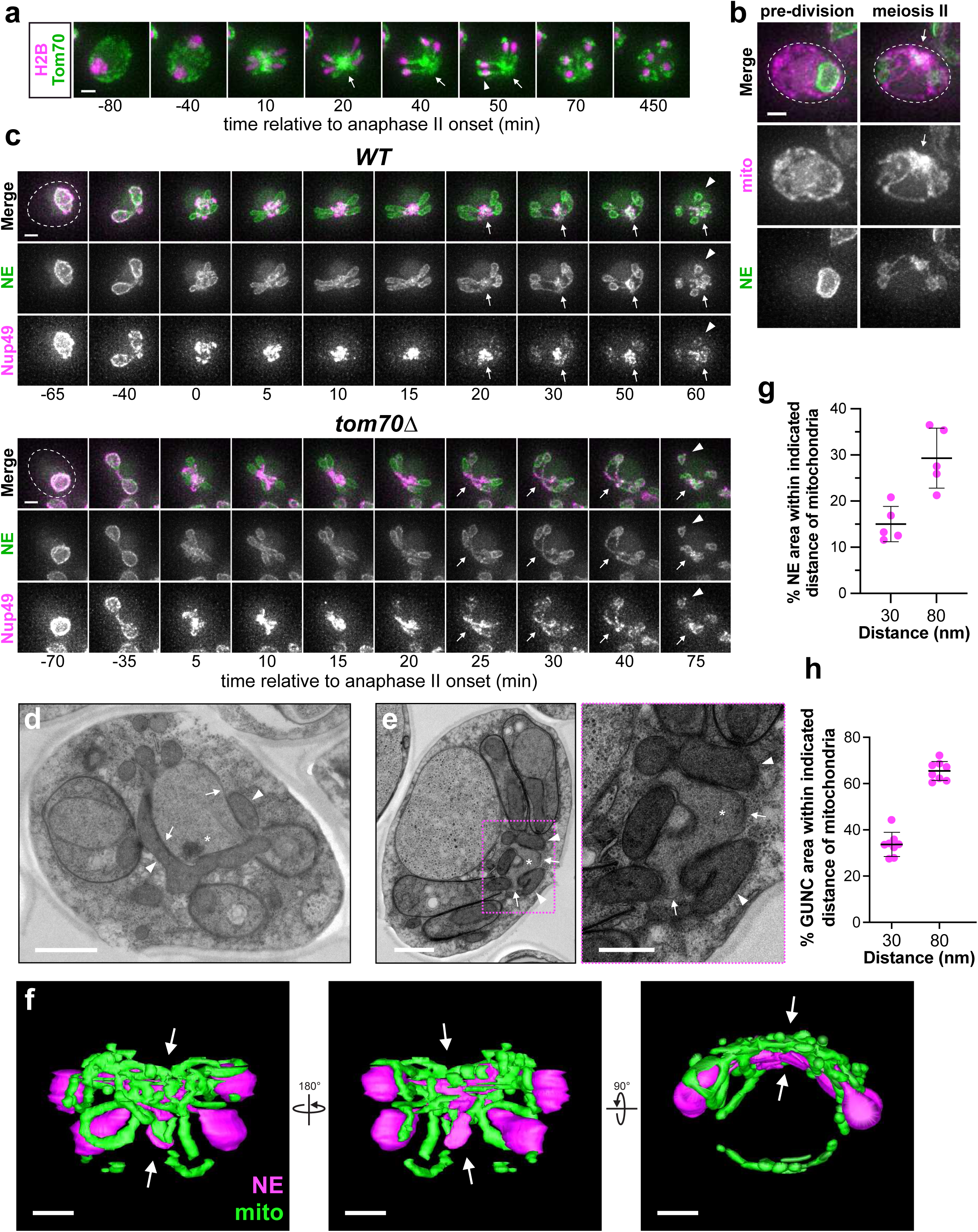
Extensive mitochondria–nuclear envelope contacts form around the GUNC during meiosis II, where Tom70 enrichment promotes nuclear envelope remodeling. **a,** Montages of Tom70-GFP alongside H2B (Htb1-mCherry) in sporulating cells. Arrows emphasize accumulations of Tom70-GFP at GUNC-apposed mitochondria in mid-meiosis, while arrowheads emphasize enrichment of Tom70-GFP in inherited nucleus-apposed mitochondrial tubules. **b,** Pre-division and meiosis II timepoints from timelapse microscopy tracking mitochondrial outer membrane (mito; Tom20^1–42^-3xmScarlet) and nuclear envelope (NE; 3xGFP-Heh2^93–378^). Arrows emphasize mitochondrial enrichment at the GUNC region. **c,** Montage of nuclear envelope morphology (NE; 3xGFP-Heh2^93–378^) and nucleoporins (Nup49-2xmScarlet) during meiosis in wild type and *tom70Δ* mutants. Arrows emphasize the GUNC region, while arrowheads indicate gamete nuclei following PSM closure. **d, e,** Transmission electron microscopy of wild-type cells in (d) early-mid anaphase II or (e) late anaphase II, with magnified inset. Asterisks mark GUNC, arrows point to nuclear envelope, and arrowheads indicate mitochondrial tubules. Scale bars; 1 µm (d and e), 500 nm (e inset). **f,** Whole-cell reconstruction of nuclear envelope (magenta) and mitochondrial network (green) from serial transmission electron microscopy of a *P_GAL_-NDT80* cell undergoing anaphase II. Arrows indicate GUNC region. Scale bars, 2 µm. **g, h,** Whole-cell percentages of either (g) total NE surface area or (h) specifically GUNC surface area within 30 nm or 80 nm of mitochondrial membranes were quantified from serial electron microscopy reconstructions. Points represent measurements from individual cells, bars indicate average ± standard deviation. Scale bar sizes (varying) are indicated above. Scale bars, 2 µm; unless otherwise indicated.

To determine whether Tom70 affects nuclear envelope architecture, we examined the localization of a 3xGFP-tagged Heh2 transmembrane domain (3xGFP-Heh2^93–378^), which labels the inner nuclear membrane (INM)^21,57,58^, alongside Nup49-2xmScarlet. In wild-type cells, the INM reporter delineates the GUNC as a distinct central nuclear compartment, coincident with NPC compaction (Fig 6c and Supplementary Video 9). In contrast, *tom70Δ* mutants showed a marked change in NE morphology, with reduced compaction of NE signal at the GUNC (Fig. 6c, arrows), alongside reduction in NPC clustering at the GUNC and an apparent misrouting of NPCs into gamete nuclei (Fig. 6c, arrowheads; Supplementary Video 10), consistent with our previous observations. Together, these findings point to a critical role for Tom70 in orchestrating NE remodeling during meiosis, potentially through influence on mitochondria-nuclear envelope contact sites rather than through conventional mitochondrial import functions.

### Extensive contact sites between mitochondria and nuclear envelope are established around the GUNC during meiosis II

To visualize membrane contact sites between mitochondria and the NE/ER at higher resolution, we performed serial-section transmission electron microscopy (TEM) on high-pressure frozen samples. We examined wild-type cells and a synchronized meiosis strain (*P_GAL_-NDT80*, *GAL4-ER*), which permits robust enrichment at specific meiotic stages^59^. As expected, mitochondria-plasma membrane contacts, prominent during vegetative growth and earlier meiotic stages, were absent in meiosis II, consistent with previous reports^19^. In contrast, we observed strikingly extensive contacts between mitochondria and the NE (mito-NE contacts), as well as its contiguous ER, specifically during meiosis II (Fig. 6d,e).

These mito-NE contacts were not detected in vegetative or early meiotic cells and did not emerge even in Num1-depleted mutants, where mitochondria detach from the cell cortex (Extended Data Fig. 6)^60,61^. This indicates that mitochondrial collapse alone is insufficient to drive these contacts, suggesting active remodeling or tethering mechanisms unique to meiosis II.

Quantitative analysis of the serial TEM data revealed that contact regions between mitochondria and NE could exceed 1000 nm in length. Whole-cell reconstructions (Fig. 6f and Supplementary Videos 11-15) showed that 15% of the NE surface was within 30 nm of mitochondria, consistent with the canonical distance defining *bona fide* membrane contact sites, while approximately 30% was within 80 nm, a relaxed threshold suggested for some contact sites (Fig. 6g)^62,63^. These contacts were particularly enriched around the GUNC: 35% of the GUNC NE was within 30 nm and 65% within 80 nm of mitochondria (Fig. 6h). These findings establish the GUNC as a focal site of extensive mito-NE contact during meiosis II, highlighting a potential role for these interfaces in nuclear remodeling and inheritance.

### Cnm1 is a meiosis-specific protein that localizes to the GUNC in a Tom70-dependent manner

Given the striking morphology and extent of mito-NE contacts observed during meiosis II, we sought to identify if Tom70-dependent molecular machinery was responsible for their formation. Tom70 has been previously implicated in membrane contact site formation through direct interactions with two distinct ER- and NE-localized proteins: Lam6/Ltc1^53,56^ and Cnm1^64^, respectively.

Although levels of Lam6, an integral ER-localized sterol transporter, increased during the meiotic divisions (Extended Data Fig. 7a), Lam6-GFP puncta were only occasionally observed near the GUNC (Extended Data Fig. 7b, arrow), where mito-NE contacts are most pronounced. Instead, Lam6 predominantly localized to puncta near gamete NE (Extended Data Fig. 7b, arrowheads) and nuclear-vacuole junctions (NVJs; Extended Data Fig. 7b, open arrowhead), a localization consistent with prior reports^53,56^. This localization was not consistent with a major role in mediating the extensive mito-NE contact we observed at the GUNC. Furthermore, deletion of *LAM6* had no discernible effect on mitochondrial behavior (Extended Data Fig. 7c), arguing against a primary role in GUNC-associated contact formation.

Cnm1, in contrast, is a NE-specific transmembrane protein that mediates mitochondrial-nuclear tethering through direct interaction with Tom70^64^. Although originally identified in an overexpression screen in mitotically dividing cells, the physiological context of Cnm1 tethering function has remained unclear^64^. Excitingly, we found that *CNM1* is a meiosis-specific gene: it is robustly upregulated during meiosis II, at both the transcript and protein levels, with minimal expression at earlier meiotic stages and in vegetative cells (Fig. 7a)^65,66^. To monitor its dynamics, we generated a Cnm1-GFP fusion at the endogenous locus. Immunoblotting confirmed a sharp increase in Cnm1-GFP signal at ∼7.5 hours in sporulation (SPO), corresponding to meiosis II, followed by a decline thereafter (Fig. 7b). This induction was dependent on the meiosis-specific transcription factor Ndt80 (Fig. 7b)^67^, which activates mid-meiotic gene expression by binding to mid-sporulation elements (MSEs)^68^. Furthermore, repression of Cnm1 expression outside of meiosis was dependent on Sum1, a transcriptional repressor of MSE-regulated genes (Extended Data Fig. 7d)^69,70^. Consistently, the *CNM1* promoter contains several MSE-like motifs, indicating that *CNM1* is likely a *bona fide* Ndt80 and Sum1 target. These findings demonstrate that *CNM1* upregulation is a programmed feature of the meiotic transcriptional cascade, rather than a response to nutrient deprivation.

**Figure 7:**
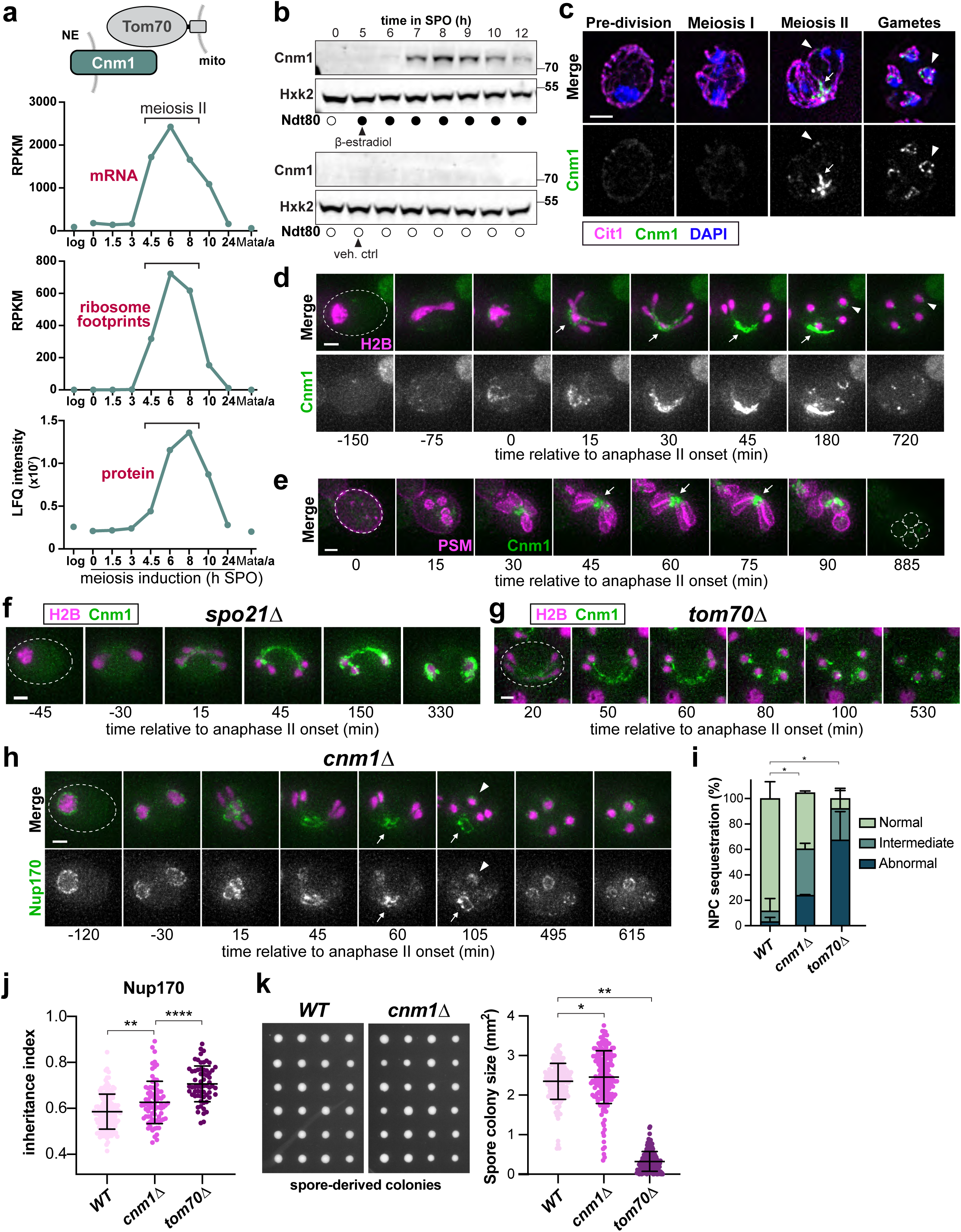
The meiosis-specific protein Cnm1 localizes to the GUNC in a Tom70-dependent manner and contributes to GUNC-mediated exclusion. **a,** Schematic (top) of bipartite tethering complex between mitochondrial (mito) Tom70 and nuclear (NE) resident Cnm1. Expression (below) of *CNM1* either during log-phase vegetative growth in rich media (veg), the indicated time in sporulation media (h SPO), or in control non-sporulating diploids (Mat**a**/**a**) plotted from previously generated meiotic datasets ^66^. Top plot indicates transcript levels (RNA-seq), middle plot indicates translation levels (ribosome-protected footprints), and bottom plot indicates protein levels (mass spectrometry). Time window corresponding to execution of meiosis II by cells within the semi-synchronized population is indicated by brackets. **b,** Western blotting of endogenously-tagged Cnm1-GFP at indicated time points in SPO upon synchronized entry into the meiotic divisions by induction of *P_GAL_-NDT80* (1 µM β-estradiol, top), or repression of *P_GAL_-NDT80* and prophase arrest (vehicle control, bottom). Hxk2 serves as a loading control. **c,** Images of Cnm1-GFP-expressing *P_GAL_-NDT80* cells, fixed 2.5h after *NDT80* induction as in (b), alongside Cit1-mCardinal-marked mitochondria and DAPI-stained chromatin. Arrows indicate Cnm1-GFP colocalization along mitochondrial tubules, arrowheads indicate Cnm1-GFP at spore nuclei. **d, e,** Montage of sporulating cell expressing Cnm1-GFP alongside either (d) Htb1-mCherry marked chromatin (H2B), or mKate-Spo20^51–91^ marked gamete plasma membranes (PSM). Arrows mark Cnm1-GFP localization to the GUNC, while arrowheads mark localization at spore nuclei. Montages of **f,** *spo21Δ* or **g,** *tom70Δ* mutants expressing Cnm1-GFP and Htb1-mCherry (H2B) during meiosis. **h,** Montages of nucleoporin Nup170-GFP relative to chromatin (Htb1-mCherry; H2B) in sporulating *cnm1Δ* mutants (compare to wild type in Fig. 2a). **l,** Quantification of abnormal nucleoporin (NPC) sequestration in *cnm1Δ* mutants, with wild type and *tom70Δ* data replotted from Fig. 2c. Shown is the average ± standard deviation from two experiments, n = 144-153 cells each. * (p < 0.05), two-tailed unpaired t-test. **j,** Inheritance index quantification for Nup170-GFP within *cnm1Δ* mutants is plotted against wildtype and *tom70Δ* (data from Fig. 2f). The average ± standard deviation from a single experiment is shown, n = 79. Only cells forming four closed spores that remained intact through the entire 18h movie were quantified. **** (p<0.0001), Kolmogorov-Smirnov test. **k,** Representative image (left) of colonies grown for 48h on YPD at 30°C from individual gametes, with quantification (right). Individual colony sizes from two independent experiments are plotted, n = 93-94 cells each. Wild type and *tom70Δ* data are reproduced from Fig. 4. Scale bars, 2 µm.

To determine whether the timing of Cnm1 expression coincides with GUNC-associated tethering, we next examined its subcellular localization. Cnm1-GFP became concentrated around the GUNC during meiosis II, localizing between DAPI-stained nuclei and colocalizing with mitochondrial tubules, marked by Cit1-mCardinal (Fig. 7c, arrows; Extended Data Fig. 7e). Time-lapse imaging in cells co-expressing Htb1-mCherry revealed that Cnm1-GFP first appeared at the onset of anaphase II and was predominantly confined to the GUNC region (Fig. 7d, arrows; Supplementary Video 16; Extended Data Fig. 7f), with smaller foci or patches persisting at gamete nuclei (Fig. 7d, arrowheads; Extended Data Fig. 7f).

Cnm1 enrichment at the GUNC was further supported by co-visualization with a PSM marker. In cells expressing mKate-Spo20^51–91^, which labels PSMs, Cnm1-GFP was largely restricted within the PSM-defined boundaries of the GUNC (Fig. 7e and Supplementary Video 17). This spatial confinement was completely disrupted in *spo21Δ* mutants, which lack proper PSM and GUNC formation; in these cells, Cnm1-GFP was uniformly distributed along the NE during meiosis II (Fig. 7f), suggesting that PSM assembly is required to concentrate Cnm1 at the GUNC.

Importantly, deletion of *TOM70* (*tom70Δ*), the mitochondrial receptor for Cnm1^64^, also resulted in loss of Cnm1-GFP enrichment at the GUNC, instead producing a uniform NE signal reminiscent of *spo21Δ* mutants (Fig. 7g). These findings indicate that Tom70 is required for localization of Cnm1 to meiosis-specific mito-NE contact sites. Taken together, our data suggest that Cnm1-Tom70 mediated tethering spatially positions mitochondria at the nuclear envelope around the GUNC during meiosis II, forming extensive contact sites that influence nuclear organization and remodeling.

### Cnm1 contributes to GUNC-mediated exclusion

Given the specific expression and localization patterns of Cnm1, we hypothesized the endogenous context of function for this tethering protein may occur at the GUNC. To probe Cnm1 function, we examined nuclear envelope morphology and nucleoporin inheritance using time-lapse imaging of Nup170-GFP in *cnm1Δ* null mutants. *cnm1Δ* mutants exhibited significant defects in nucleoporin sequestration and inheritance relative to wild type, though partially phenocopying those observed in *tom70Δ* cells (Fig. 7h-j and Supplementary Videos 18 & 19). However, unlike *tom70Δ, cnm1Δ* mutants displayed no obvious reduction in gamete fitness, as assessed by spore colony size (Fig. 7k). Meiotic progression kinetics, sporulation efficiency and gamete viability were also indistinguishable between wild type and *cnm1Δ* (Extended Data Fig. 7g-i). Since loss of a single tether is often compensated for by the presence of additional tethers^53,56,71^, we wondered whether this discrepancy between *tom70Δ* and *cnm1Δ* tether mutants might result from remaining Tom70-dependent tethering in *cnm1Δ* mutants, through Lam6. However, combined deletion of *LAM6* and *CNM1* did not exacerbate phenotypes related to NE structure, NPC inheritance, or spore viability/colony size (Extended Data Fig. 7j-l) relative to *cnm1Δ* single mutants, suggesting limited functional redundancy between these two tethers. These findings highlight the connection between mito-NE contacts and nuclear remodeling, as Cnm1 is associated with the tethering function of Tom70, but not its non-contact functions. Furthermore, they raise the intriguing possibility that other yet unidentified nuclear envelope- or ER-associated proteins may act alongside Cnm1 to mediate Tom70-dependent effects on nuclear remodeling during meiosis.

### Distinct domains of Tom70 mediate nuclear envelope remodeling and mitochondrial-NE contact site formation through both Cnm1-dependent and - independent pathways

The cytosolic-facing region of Tom70 is comprised of three major domains, each consisting of tetratricopeptide (TPR) repeats: the N-terminal “Clamp” domain (C1), which binds cytosolic chaperones via EEVD motifs; the central “Core” domain (C2), which recognizes mitochondrial targeting signals (MTSs) on precursor proteins; and the “C-tail” domain (C3), which also binds client MTSs (Fig. 8a)^43–45,72–74^. To determine which domains contribute to NPC sequestration, we expressed full-length Tom70, various domain truncation mutants, or an empty vector under the endogenous *TOM70* promoter in a *tom70Δ* background. Notably, Cnm1 is thought to interact with Tom70 through an internal MTS-like sequences^64,75^, presumed to associate with the C2 or C3 domains.

**Figure 8:**
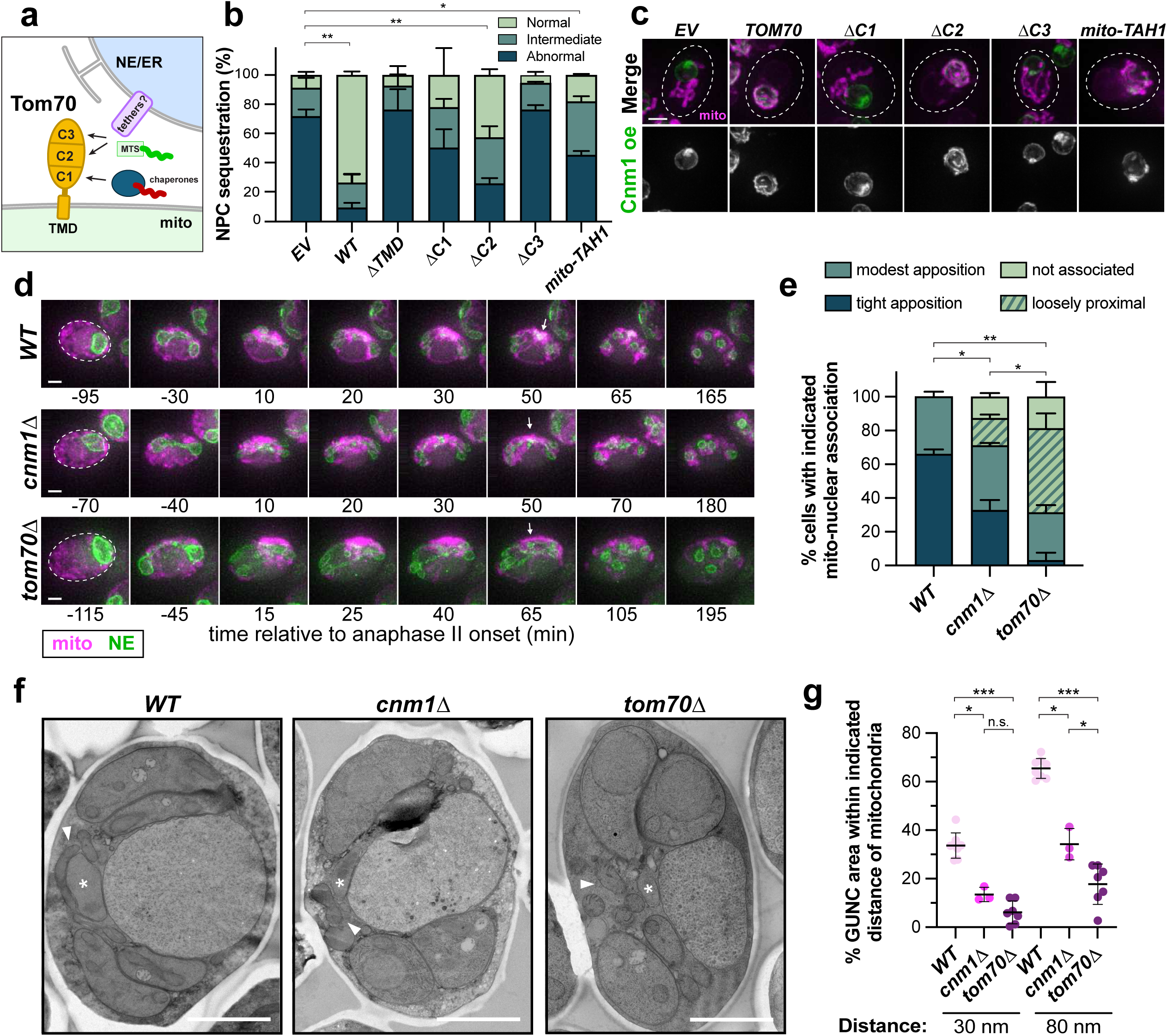
Tom70 promotes nuclear envelope remodeling and mito–NE contact formation through Cnm1-dependent and -independent mechanisms. **a,** Schematic of Tom70 domain organization and functions in both mitochondrial import and partner binding. Misfolded proteins (red) associate with Tom70 via chaperones at the C1 domain, while MTS-containing proteins (green) can directly bind the C2 and C3 domains. **b,** Penetrance of abnormal nucleoporin sequestration (Nup170-GFP) was quantified in *tom70Δ* mutants rescued with empty vector (*EV*), full-length Tom70^WT^, deletion alleles for each domain (with indicated residues removed: ΔTMD=2-41, ΔC1=99-214, ΔC2=148-460, ΔC3=461-617), or the C1-like mito-Tah1. Shown is the average ± standard deviation from two experiments, n = 78-131 cells each. * (p<0.05), ** (p<0.01), *EV* vs *ΔTMD* (p = 0.71), *EV* vs *ΔC1* (p = 0.15), *EV* vs *ΔC3* (p = 0.39); two-tailed unpaired t-test. **c,** Representative still images of mitotically dividing *tom70Δ tom71Δ* mutant cells overexpressing Cnm1-GFP (expressed under the strong constitutive *P_GPD_/P_TDH3_*promoter) with TMRM-stained mitochondria (mito), rescued with *EV* or *TOM70* constructs as in (b). Cells were grown to log-phase in YPD. Contrast scaling of TMRM-staining was altered independently to make mitochondrial structure equally apparent in each shown cell. Scale bars, 2 µm. **d,** Montage of sporulating wild type, *cnm1Δ,* and *tom70Δ* cells expressing both 3xGFP-Heh2^93–378^ (NE) and Tom20^1–42^-3smScarlet (mito) markers from the *P_RDL1_* promoter. Arrowheads indicate mitochondrial clustering around the GUNC to varying degrees in individual *WT*, *cnm1Δ* and *tom70Δ* mutants. **e,** Quantification of wild type, *cnm1Δ* and *tom70Δ* mutants in (D) by binning strength of mitochondrial enwrapping of the GUNC. Shown is an average ± standard deviation from two experiments, n = 42-55 cells per genotype total. * (p<0.05), ** (p<0.01), two-tailed unpaired t-test of the “tight apposition” bin. **f,** Select transmission electron microscopy of *WT*, *cnm1Δ* and *tom70Δ* mutants. Asterisks denote the forming GUNC compartment, while arrowheads emphasize mitochondrial tubules. **g,** The degree of mitochondrial-nuclear contact was quantified by 3D reconstruction of GUNC volumes by serial transmission electron microscopy. The percentage of GUNC area with 30nm or 80nm of mitochondria are shown for individual cells of the indicated genotype in late meiosis II. * (p<0.05), *** (p<0.001), n.s. (p = 0.17); two-tailed Mann-Whitney U test. Scale bars, 2 µm.

If Tom70 promotes NPC sequestration primarily via its roles in proteostasis or mitochondrial import, the chaperone-binding C1 domain might be critical. In contrast, if it acts primarily through tethering to the nuclear envelope via Cnm1, then the C2 and/or C3 domains presumed to mediate MTS-like interactions should be required. Surprisingly, we found that all three domains contribute to NPC sequestration to varying degrees (Fig. 8b). The C3 domain proved most essential: cells expressing Tom70^ΔC3^ failed to rescue the NPC sequestration defect, resembling *tom70Δ* mutants harboring either empty vector or a cytosolic Tom70^ΔTMD^ construct (Fig. 8b). The C2 domain was least critical, as Tom70^ΔC2^ conferred substantial rescue (Fig. 8b). Deletion of the C1 domain (Tom70^ΔC1^) resulted in an intermediate phenotype (Fig. 8b). The C1 domain of Tom70 was previously shown to be unstable on its own, preventing sufficiency experiments using this domain; however, the EEVD-binding TPR-containing protein Tah1 is competent to substitute for the C1 of Tom70, enabling sufficiency experiments for a mitochondrial chaperone-binding domain^43^. Such targeting of Tah1 to mitochondria partially suppressed the NPC sequestration defect in *tom70Δ*, providing further support of a role for the C1 domain in GUNC-mediated sequestration (Fig. 8b). Importantly, all truncation mutants were expressed at similar levels to wild-type Tom70 (Extended Data Fig. 8a), ruling out differences in protein abundance as a confounding factor. Similar trends were also observed in the rescue of *tom70Δ* sporulation efficiency, suggesting similar Tom70-dependent functions may contribute to nuclear remodeling and robust gamete formation (Extended Data Fig. 8b).

Strikingly, the same domain dependencies were observed for mito-NE tethering. In mitotically dividing cells expressing full-length Tom70, overexpression of Cnm1-GFP recruited mitochondria to the nuclear envelope, with clear enrichment along mitochondrial tubules. In contrast, in the absence of *TOM70* or in cells expressing Tom70^ΔC3^ or Tom70^ΔC1^, Cnm1-GFP was largely diffuse throughout the NE and mitochondria remained untethered, phenocopying *tom70Δ* mutants (Fig. 8c). Tom70^ΔC2^ mutants, on the other hand, retained the ability to tether mitochondria. Mitochondrially-targeted Tah1 also partially restored mitochondrial recruitment in some cells, although Cnm1-GFP localization remained largely unaltered (Fig. 8c).

The shared domain requirements for NPC sequestration and mitochondrial tethering suggest that Tom70-dependent mito-NE contacts may underlie its role in GUNC-mediated nuclear remodeling. However, a simple model in which loss of Cnm1-mediated tethering explains all *tom70Δ* phenotypes does not align with our data: NPC sequestration defects and nuclear envelope morphology are more severe in *tom70Δ* than in *cnm1Δ* mutants (Fig. 7h and 7i). This distinction implies that Tom70 may participate in multiple tethering interactions at the NE/ER, beyond Cnm1 alone. Supporting this, both time-lapse fluorescence microscopy and serial-section transmission electron microscopy revealed a pronounced reduction in mito-NE contact at the GUNC in *tom70Δ* mutants, exceeding that observed in *cnm1Δ* cells (Fig. 8d-g and Supplementary Videos 20-22). Altogether, our data support a model in which Tom70 promotes NPC sequestration and GUNC-mediated exclusion during meiosis through multiple tethering interactions, with functional contributions from its chaperone-binding, MTS-recognition, and membrane-anchoring domains.

## Discussion

In this study, we identified Tom70, a conserved mitochondrial outer membrane protein, as a critical mediator of nuclear envelope remodeling during meiosis, specifically in the formation and function of the Gametogenesis Uninherited Nuclear Compartment (GUNC). Surprisingly, this role could not be attributed to Tom70’s canonical function in mitochondrial import. Instead, our results point to a distinct, meiosis-specific function: facilitating extensive physical contacts between mitochondria and the nuclear envelope. These contacts appear crucial for ensuring proper exclusion of nuclear pore complexes (NPCs) and other nuclear constituents (nucGEM) from inherited gamete nuclei. Thus, our findings position organelle remodeling events, particularly mitochondria-nucleus tethering, as integral to nuclear compartmentalization and gamete fitness.

Tom70’s diverse roles at the mitochondrial surface–including chaperone docking, protein import, and organelle tethering–have been studied for decades (reviewed in ^76^), but its function during meiosis remained unexplored. We show here that Tom70 is required during meiosis for proper nuclear sequestration, temporally coinciding with the appearance of meiosis-specific mitochondria-nuclear envelope contacts. Several lines of evidence argue that the observed NPC sequestration defect of *tom70Δ* mutants is driven by impaired organelle tethering rather than disrupted mitochondrial protein import: (1) general disruption of mitochondrial import did not phenocopy *tom70Δ*; (2) *cnm1Δ* mutation, which disrupts a known Tom70-dependent tether, partially recapitulated *tom70Δ* nuclear exclusion defects; (3) all Tom70 truncations that failed to rescue NPC sequestration also failed to restore mito-nuclear tethering; and (4) ultrastructural analysis suggested that additional tethering components may also be lost or misregulated in *tom70Δ* mutants. Together, these findings support a model where Tom70-dependent membrane contact sites, rather than mitochondrial import activity, enable GUNC function. We tested whether additional Tom70-dependent or independent tethers might operate in parallel to Cnm1 to support GUNC-mediated exclusion. Lam6, an ER-mitochondrial tether, and the ERMES complex^77^ were both examined. However, disruption of ERMES or deletion of *LAM6* had no discernable effect on GUNC function. (Extended Data Fig. 8c-d). This suggests that the meiosis-specific mito-nuclear tether comprised of Tom70 and Cnm1 may be uniquely specialized for ensuring robust GUNC-mediated exclusion. Importantly, our findings suggest the contribution of other, as-yet-unidentified tethers. Future genetic screens or proteomic profiling of contact site components during meiosis may reveal the full complement of players involved.

What function(s) might this specialized contact site serve? One attractive model is that mitochondria contribute to local lipid remodeling of the nuclear envelope during meiosis II, potentially generating lateral domains that restrict diffusion of NPCs and other nuclear materials. Membrane contact sites are hubs for lipid and metabolite exchange (reviewed in^78–83^), and whole-cell lipidomic data have suggested an overall elongation of acyl chain length during meiosis^84^. While this signal may reflect lipid remodeling at the gamete plasma membrane, nuclear envelope-specific alterations remain a plausible mechanism for regulating compartmentalization.

Alternatively, the mitochondria may serve a structural role, positioning or wrapping around a subregion of the nucleus to physically constrain NPC mobility or bias membrane curvature. This steric or mechanical contribution could enhance the sorting of sequestered components into the GUNC. However, such a model is challenged by the lack of NPC sequestration/exclusion defects in ERMES mutants, which exhibit dramatic mitochondrial disorganization. Thus, mitochondrial positioning alone may contribute to, but is unlikely to fully account for, the observed effects.

Importantly, our data also reveal that sequestration of different GUNC cargoes (e.g. NPCs vs. nuclear protein aggregates) may be uncoupled. In *tom70Δ* mutants, exclusion of NPCs and nucGEMs, but not of senescence-associated factors, is compromised. This suggests that distinct biophysical properties or mechanistic pathways govern the targeting of different cargoes to the GUNC. This opens the door to genetic separation-of-function approaches to dissect the molecular underpinnings of GUNC selectivity and their respective contributions to gamete rejuvenation. Further identification of the molecular players that enable cargo sorting to the GUNC will be critical for understanding how cells reset aging hallmarks and ensure the production of healthy offspring.

Does Tom70-dependent mitochondrial enrichment around the GUNC also influence mitochondrial segregation and quality control? In metazoan germlines, evidence supports preferential inheritance of functional mitochondria, but the underlying mechanisms remain poorly defined^85–91^. In budding yeast, approximately half of the mitochondrial content is inherited by gametes, with the remainder eliminated, but whether these pools differ functionally is unknown^92^. Intriguingly, Tom70 is required for the formation of prominent mito-NE contacts around the GUNC. Loss of *TOM70* results in gametes with reduced fitness, often forming small, respiration-deficient colonies (Fig. 4d and Extended Data Fig. 8e). These findings raise the possibility that mito-NE tethering via Tom70 acts as a quality-control checkpoint, favoring the inheritance of healthier organelles. Direct visualization and functional assessment of mitochondria during meiosis will be essential to determine whether this spatial enrichment indeed drives selective segregation.

Our findings may have broader implications for gamete formation and health. Multiple instances of mitochondria-nucleus proximity are observed during oogenesis and spermatogenesis in animals, from the Balbiani body in vertebrate oocytes to the nebenkern in *Drosophila* spermatids^93–95^. Moreover, NPC exclusion and compartmentalization are conserved features of metazoan gametogenesis^96–99^. Expression of *TOM70* orthologs appears upregulated during gametogenesis in several species^100–102^, hinting at conserved roles that remain to be tested. By uncovering a role for mito-nuclear contact sites in regulating nuclear envelope remodeling and selective inheritance during yeast meiosis, our study calls for further investigation into the possible roles of membrane contacts in gamete quality control across evolution.

## Methods

### Yeast strains, plasmids, and primers

All yeast strains and plasmids in this study are listed in Table S1 and Table S2, respectively. The origins of previously constructed alleles or plasmids are indicated and attributed both below and within Table S1 and Table S2, respectively. Constructs were generated by Gibson assembly (New England Biolabs), site-directed-mutagenesis (New England Biolabs), or standard restriction enzyme-based cloning (New England Biolabs). Relevant sequence sources and primers are described within Table S3. Plasmids were verified by Sanger and/or whole-plasmid Nanopore sequencing. All *Saccharomyces cerevisiae* strains in this study are derivatives of SK1 ^5^. Deletion and endogenous N- or C-terminal tagging of genes were performed using standard PCR-based methods, with the primers used listed in Table S3^105–107^. Single-copy transgenes were integrated into the genome following excision of the desired sequence via digestion of flanking PmeI sites. Yeast transformation was performed using the lithium acetate method^108^. Proper integration was verified by PCR, followed by fluorophore visualization, Western blot, or sequencing, where applicable. Transformants were backcrossed prior to subsequent crossing and use.

The following alleles were constructed in a previous study: *spo21Δ*, *P_ATG8_-GFP-SPO20*^51–91^*, CIT1-GFP*, *CIT1-mCardinal*, and *P_CUP1_-OsTIR1* ^19^; *HTB1-mCherry* and *P_GPD_-GAL4-ER* ^109^; *NUP170-GFP*, *POM34-GFP*, *NUP2-GFP*, *P_CUP1_-OsTIR1^F74G^, HSP104-mCherry*, *NSR1-GFP*, and *DON1-GFP* ^21^; *HTA1-mApple*, *rDNA::5xtetO*, *P_REC8_-TetR-GFP* ^110^; *HSP104-GFP* ^12^; *P_PMA2_-PMA2-GFP*, *P_tetO_-STE5*, *P_RNR2_-TetR-TUP1* & *P_tetO_-TetR*, *flo8*, *amn1^BY4741^* ^42,111^; and *P_GAL_-NDT80* ^59^.

The following used alleles were generated or introduced in this study: *tom70Δ*, *NUP49-GFP*, *P_tetO_-TOM70, sam37Δ, TOM20-3V5-IAA17*^71–114^, *NUP49-2xmScarlet*, *CNM1-GFP*, *cnm1Δ*, *SUM1-3V5-IAA7*, *MMM1-3V5-IAA7*, *lam6Δ*, *lam5Δ*, and various *MX* cassette marker swaps of previously generated alleles. For additional information, see Tables S1-S3.

The nucGEM construct pLH941 (pRS305_pINO4-SV40NLS-PfV-GS-mTSapphire) was a kind gift from Liam Holt^34,35^. nucGEM CDS was inserted into a pNH604 single-integration vector alongside a *P_ATG8_* promoter fragment to drive strong meiotic expression. The resulting construct was integrated at the *TRP1* locus, as briefly described above.

The *P_tetO_* promoter was introduced at the endogenous *TOM70* locus by standard PCR-based methods^105^. First, a nourseothricin-selectable promoter swapping construct (a gift from Fabian Rudolph; FRP2375) was subcloned to reintroduce standard S4 and S1 tagging primer adaptors^41^. Then *P_tetO_* was integrated immediately prior to the *TOM70* start codon position for either wild-type *TOM70*, *TOM70-3v5*, or *Tom70-GFP*, in strains expressing the TetR repressor construct^41^.

*TOM70* truncation and AID alleles were constructed as follows. First, the wild-type SK1 *TOM70* CDS, along with a flanking 1kb of the native sequence context on each side (which should include the complete endogenous promoter and terminator elements), was inserted into a modified pNH604 vector (pNH604CR, which has improved 3’ homology to ensure excision of a remnant *hisG* cassette at the *TRP1* locus during integration). Subsequently, site-directed mutagenesis was employed to remove desired sequence (e.g. C1 Clamp, residues 99-214; C2 Core, residues 248-460; C3 C-tail, residues; 461-617) by selective amplification and blunt-end recircularization. The CDS of cytosolic *TOM70* (residues 42-617) was replaced by Gibson assembly with that of *TAH1*, amplified from wild-type genomic DNA, to generate the *mito-TAH1* construct. The resulting transgenes were integrated at the *TRP1* locus and strains were constructed in the absence of TOM70 at the endogenous locus (*tom70Δ*). In order to assess protein levels of the truncations, a single V5 tag flanked by GGGS linkers was inserted following *TOM70* residue 51 (between the transmembrane and structured cytosolic domains) via site-directed mutagenesis using tailed primers with the tag sequence.

For Tom70-AID, a V5 tag was inserted following TOM70 residue 86, in the manner described above. Residue 86 also lies between the transmembrane and structured cytosolic domains. Next, a minimal AID fragment (IAA17^71–114^) was inserted after this tag by Gibson assembly^112^. AID insertion at this site resulted in slightly improved degradation kinetics relative to insertion after residue 51. We found standard C-terminal tagging of *TOM70* with full-length 3V5-IAA17 perturbed nucleoporin sequestration, even in the absence of the TIR1 transgene (46.9% of *TOM70-AID^C-term^* cells exhibited abnormal NPC sequestration, compared to a typical 5% of wild type cells). Such C-terminal tagging of Tom70 also altered retention of Cnm1 with bulky GFP tags, further suggesting larger C-terminal tags disrupt Tom70 function. Insertion sites were chosen based on AlphaFold-predicted Tom70 structure and sites tolerating insertion between *TOM70* ortholog comparisons. AID experiments were performed in *tom70Δ* mutants carrying only one copy of the TOM70-AID transgene in order to speed clearance of the transgene, enabling efficient depletion prior to the meiotic divisions. Importantly, a single copy of *TOM70* was haplosufficient for NUP sequestration.

The nuclear envelope reporter was generated by modification of a previously described GFP-h2NLS-L-TM construct derived from yeast INM protein Heh2^21,57^. Briefly, pUB1196 was subcloned by Gibson assembly to replace the *P_ARO10_* promoter with a strong *P_ATG8_* promoter from pUB949. This strongly-expressed construct drove formation of nuclear envelope protrusions and deformations, as previously observed upon *P_GAL_*induction during vegetative growth. Addition of an additional two GFP copies and swapping to the more modest *P_RDL1_* promoter by Gibson assembly resulted in bright signal with minimal perturbation nuclear envelope morphology. A red-channel NE reporter was created through restriction enzyme cloning was used to replace 3xGFP from the resulting construct with three copies of mScarlet-I, individually amplified from a gBlock (IDT).

A generic outer mitochondrial membrane marker was generated by Gibson assembly of the Tom20 signal anchor (residues 1-42) followed by a short linker and 3xmScarlet from the above NE marker into a pNH603 *HIS3* single-integration vector under the control of the *P_RDL1_* promoter.

A *P_GPD_-CNM1-GFP* construct for overexpression of Cnm1 outside of meiosis was generated by first subcloning the *CNM1* CDS under the strong mitotic promoter *P_GPD_* (*P_TDH3_)* into a modified pNH605 *LEU2* single-integration vector (with improved targeting homology, pLC605^19^), via Gibson assembly. Then, linker and *GFP* sequence was appended at the C-terminus of *CNM1* by a second Gibson assembly.

### Sporulation conditions

For *S. cerevisiae*, strains were first thawed onto YPG (1% yeast extract, 2% peptone, 3% glycerol, 2% agar) plates and grown overnight at 30°C, before transfer to YPD4% (1%yeast extract, 2% peptone, 4% glucose, 2% agar) plates to prevent premature sporulation during further outgrowth. Cells were then grown in liquid YPD (1% yeast extract, 2% peptone, 2% glucose, 22.4 mg/L uracil, and 80 mg/L tryptophan) at room temperature RT for ∼24 h, reaching saturation (OD_600_ ≥ 10). Cultures were then backdiluted in BYTA (1% yeast extract, 2% bacto tryptone, 1% potassium acetate, and 50 mM potassium phthalate) to an OD_600_ = 0.25 and incubated at 30°C for 16 h, reaching an OD_600_ ≥ 5. To induce sporulation, cells were pelleted (3 krcf, 2 min), washed with MilliQ water, and resuspended in sporulation media (SPO: 2% potassium acetate with 0.04 g/L adenine, 0.04 g/L uracil, 0.01 g/L histidine, 0.01 g/L leucine and 0.01 g/L tryptophan, adjusted to a pH of 7 with acetic acid) at an OD_600_ = 1.85. Cultures were maintained at 30°C on a shaker for the duration of the experiment. To ensure proper aeration during each outgrowth growth, liquid cultures were shaken in flasks that held ≥10x culture volume. For sporulation experiments using the *P_GAL_-NDT80* system, cells were arrested in prophase prior to release by addition of 1 µM β-estradiol (or held, by addition of EtOH vehicle control) at 5h in SPO. Peak meiosis II division in the population occurs roughly 2.5 h following *P_GAL_-NDT80* release, or after 7.5 h total culture in SPO for unsynchronized experiments.

### Sporulation efficiency, spore viability, and germination

Sporulation of cells was assessed after 24 h culture in SPO media under a light microscope. Portions of progenitors forming 0 spores (unsporulated), 1-2 spores (monads/dyads), or 3-4 spores (triads/tetrads) were scored. Spore viability was assessed by digesting sporulated cells from a 24 h SPO culture with 1 mg/mL zymolyase (MP Biomedicals) at room temperature for 3 minutes, then dissecting individual spores from tetrads onto YPD plates. After 48 h of growth at 30°C, plates were imaged and replica plated to YPG to assess portion of respiratory-incompetent spore colonies. Spore colony size was quantified in FIJI by image thresholding and particle size analysis. Germination speed was assessed by timelapse microscopy, further described below. Briefly, ascus-packaged spores from a 24 h SPO culture were separated into single spores by 3h digestion in digestion buffer (0.1 mg/mL zymolyase, 2% β-mercaptoethanol, in H_2_O) at 30°C, followed by six rounds of twenty 50% sonication pulses (Branson; 100-132-1623R & 101-135-066R). Separated spores were diluted to an approximate OD of 1.5 then adhered to Concanavalin A-treated 96-well plates (see below). 100 µL liquid YPD was added to each well and the emergence of first bud scored for each spore, relative to time of rich media addition.

### Aged cell isolation and sporulation

Aged cell meiotic timelapse microscopy experiments were performed as previously described ^21^. Briefly, cells were biotin labeled then aged progenitor were later reisolated following an outgrowth step using an anti-biotin magnetic bead-sorting assay^40^. For each strain, an overnight YPD culture was backdiluted and grown to log-phase at 30°C. Then, 8 OD of cells were washed with three times with PBS pH 8 and labeled with 8 mg/ml EZ-Link Sulfo-NHS-LC-biotin (ThermoFisher Scientific) in PBS for 30 min at 4°C. Cells were washed once in PBS pH 8 100 mM glycine to remove excess biotin. Biotinylated cells were grown for 16 hours in YPD with 100 μg/ml ampicillin at 30°C, from a starting OD_600_ of 0.08 or 0.14. Cells were subsequently harvested, washed in PBS pH 7.4 0.5% BSA buffer, and mixed with 100 μl of anti-biotin magnetic beads (Miltenyi Biotechnology) for 15 min at 4°C. Cells were washed with PBS pH 7.4 0.5% BSA buffer and sorted magnetically using LS depletion columns with a QuadroMacs sorter, washing three times with PBS pH 7.4 0.5% BSA. A fraction of the initial flow-through (biotin-negative) was kept as young cells alongside eluted aged cells (biotin-positive) old cells. Both young and old cell collections were budscar labeled for 15 min at room temperature using 1 μg/ml Wheat Germ Agglutinin, Alexa Fluor 350 Conjugate (ThermoFisher Scientific). A mixture of aged and young cells was subsequently washed once in H_2_O and twice with SPO media. The cell mixture was resuspended with SPO at a cell density of OD_600_ = 1.85 with 100 μg/ml ampicillin and incubated at 30°C. The number of doublings in subsequent experiments was measured by counting the number of budscars.

The replicative lifespan of gametes from aged and young progenitors was determined as previously described ^42^. Briefly, aged and young cells were isolated as described above, with an additional biotin sorting step to increase purity of aged cells. Young and old cell collections were budscar labeled as above, along with an additional anti-streptavidin 564 nm dye at 1 μg/mL (product info). Cultures were incubated for 24 hours in SPO media with shaking at 30°C in the dark to sporulate. 0.1 OD_600_ units of cells were fixed (see below) before and after the 24h SPO culture to assess pre- and post-sporulation age by counting biotinylation and bud scars. Post-sporulation age-sorted cultures were pelleted and resuspended in 1 mL digestion buffer (0.1 mg/mL zymolyase and 2% β-mercaptoethanol in H_2_O), then incubated for 3 hours on a nutator at 30°C. Cells were washed once and resuspended in H_2_O, the separated by repeated sonication (6x 20 seconds, 50% pulses). Upon separation, spores were filtered (30 µm Pre-separation filter, militenyibiotec) and diluted in YPD 2% + ampicillin (100 μg/mL) to an OD_600_ of 0.8. After 3 hours outgrowth at 30°C to increase gamete cell size, a 1 mL of cell suspension was aliquoted, sonicated, filtered, and loaded into the microfluidics system.

Microfluidics devices (adapted from ^113^; 18++ chips from iBiochips) were setup using the manufacturer’s recommendations on an ECHO Revolution fluorescence microscope in the inverted mode with the environmental chamber set to 30°C. Briefly, 20 mL syringes (Becton, Dickinson and Company) filled with ∼15 mL of YPD 2% + ampicillin (100 μg/mL) + 1% Pluronic F-127 surfactant (P2443, Sigma) were connected to a Clay Adams Intramedic Luer-Stub Adapter (Becton, Dickinson and Company). Non-DEHP Medical Grade Tubing (iBiochips, MTB-100, ID = 0.202”, OD = 0.060”, wall = 0.20”) equipped with a hollow stainless-steel pin (iBiochips, HSP-200) was then used to connect the media syringe to the imaging units using the “media loading” ports. “Cell loading” ports were plugged with solid stainless-steel pins (iBiochips, SSP-200) while imaging units were primed with media for at least 2 hours with a flow rate of 2.4 μL/min using micropumps (iBiochips,78-7200BIO). After all units were filled with media, pumps were stopped an hour prior to cell loading. For cell loading, 5 mL syringes (Becton, Dickinson and Company) with Clay Adams Intramedic Luer-Stub Adapter (Becton, Dickinson and Company) were filled with cells diluted to an OD_600_ of 0.8 in YPD 2% + ampicillin (100 μg/mL). Solid stainless-steel pins were then removed from the “cell loading” ports and tubing and open stainless-steel pins were used to connect the syringe containing cells with the microfluidic device. Cells were loaded manually and, once sufficient cells were loaded, the “cell loading” ports were plugged with solid stainless-steel pins and media flow was resumed at 2.4 μL/min. Timelapse imaging was performed using an Olympus 40x/1.4 NA oil-immersion objective, at 15-20 minute for 72 hours. Image acquisition for chamber was achieved with 2x27 tiled fields of view, with three 3-5 µm z-slices.

After a movie was completed and processed, single Pma2-GFP-positive cells (gametes) were identified and cropped. Next, the replicative lifespan of individual gametes was assessed in the brightfield channel by counting the number of daughter cells produced. To determine the replicative lifespan of vegetatively-grown haploid cells (non-gametes), saturated YPD 2% cultures were re-diluted to 0.2 OD_600_ in YPD 2% + ampicillin (100 μg/mL) right before loading into the microfluidics system, before proceeding as described above.

### Fluorescence Microscopy

All *S. cerevisiae* wide-field fluorescence microscopy was performed using a DeltaVision Elite microscope (GE Healthcare) running the accompanying softWoRx software (version 6.5.2), using a 60x/1.42 NA or 100x/1.40 NA oil-immersion objectives and a PCO Edge sCMOS camera. The specific acquisition settings used in each experiment are listed in Table S4. Live-cell imaging was conducted on either a CellASIC ONIX2 microfluidics platform (EMD Millipore) or in Concanavalin A-treated glass-bottom 96-well plates (Corning; 4580), as previously described^21^. For microfluidics experiments, cells were loaded into Y04E-01 CellASIC plates (EMD Millipore) at 8 psi for 5 seconds, and media flow was maintained at 2 psi for the duration of the experiment. 96-well plates were incubated for 10 minutes with 2 mg/mL Concanavalin to coat bottoms for cell adherence. Acquisition was performed in a temperature-controlled chamber at 30°C.

For meiotic live-cell imaging experiments, cells at an OD_600_ = 1.85 in SPO (see above for sporulation conditions) were sonicated and loaded into a microfluidics plate after approximately 3.5 h in SPO with shaking. Imaging began 5 h after introduction to SPO, typically continuing a further 18 hours with acquisition every 15 minutes. Conditioned SPO (prepared by filter-sterilizing media from a sporulating culture of a wild-type diploid yeast strain after 5 h at 30°C) was used as media for timelapse meiotic experiments, to match initial culture conditions in-flask and promote meiotic progression.

To prepare fixed cells from imaging (e.g. see Fig. 7 & Extended Data 7b), 900 µL of meiotic culture were fixed by addition of formaldehyde to a final concentration of 3.7%, for 15 minutes at room temperature. Cells were washed in 0.1 M potassium phosphate pH 6.4, resuspended in KP*_i_* sorbitol buffer (0.1 M potassium phosphate, 1.2 M sorbitol, pH 7.5), and stored at 4°C prior to imaging. DAPI staining was performed by adding DAPI-containing mounting media (product) to cells (1:1 volume) that had been permeabilized by either: ∼10 second 70% EtOH treatment (after which EtOH was aspirated away), or 5 minute treatment with 1% Triton X-100 in KP*_i_* sorbitol (after which, cells were immediately washed once and resuspended in KP*_i_* sorbitol).

### Transmission Electron Microscopy and Contact Quantification

Samples were prepared for electron microscopy as previously described^21^, with minor modifications. Following 7 – 8 h incubation in SPO media, yeast cells were concentrated by vacuum filtration onto a nitrocellulose membrane and then scrape-loaded into 50 µm-or 100 µm-deep high-pressure freezing planchettes^114^. Freezing was done in a Bal-Tec HPM-010 high-pressure freezer (Bal-Tec AG).

High pressure frozen cells stored in liquid nitrogen were transferred to cryovials containing 1.5 ml of fixative consisting of 1% osmium tetroxide, 0.1% uranyl acetate, and 5% water in acetone at liquid nitrogen temperature (−195°C) and processed for freeze substitution according to the method of McDonald and Webb^115,116^. Briefly, the cryovials containing fixative and cells were transferred to a cooled metal block at −195°C; the cold block was put into an insulated container such that the vials were horizontally oriented and shaken on an orbital shaker operating at 125 rpm. After 3 hr, the block and cryovials had warmed to 20°C and were transitioned to resin infiltration.

Resin infiltration was accomplished by a modification of the method of McDonald^115^. Briefly, cells were rinsed 5 times in pure acetone, removed from the planchettes, and infiltrated with Epon-Araldite resin in increasing increments. Resin (23.5g Eponate 12, 12.5g DDSA, 14g NMA, 375uL BDMA) was made fresh and degassed. Cells were kept rotating during the resin infiltration series and were spun down at 6000 x g for one minute between resin exchanges, as follows. Cell flakes were first resuspended in 33.3% resin (in acetone, v/v) and broken apart, then samples were rotated in 50% resin for 2 h. Samples were then rotated overnight in 66.7% resin. The following day, samples were exchanged to 90% resin once in the morning and once in the evening. Two 90% resin exchanges were once again performed the following day. The next day, three exchanges for 100% resin were performed, with 1 h rotation after each. Finally, cells in pure resin were placed in a silicone mould and polymerized over 2 days in an oven set to 100°C.

Blockfaces were trimmed and serial sections of 70 nm or 100 nm were cut on a Reichert-Jung Ultracut E microtome and picked up on 1 × 2 mm slot grids covered with a 0.6% Formvar film. Sections were post-stained with 1% aqueous uranyl acetate for 10 min and lead citrate for 10 min (Reynolds, 1963). Images of cells on serial sections were taken on an FEI Tecnai 12 electron microscope operating at 120 kV equipped with a Gatan Ultrascan 1000 CCD camera.

Models were constructed from serial sections with the IMOD package^117^, using 3DMOD version 5.1.1. Initial alignment was performed through manual translation and rotation using the Midas tool in the ETomo interface of the IMOD package. Subsequently, fine adjustments and minor warping was performed in Midas to improve alignment. Only cells in mid-late meiosis II, with four PSMs, and for which the entirety of the GUNC region was captured in a serial stack were considered for further segmentation. The nuclear envelope (discerned by its double membrane and a characteristic nucleoplasm staining) and mitochondrial (discerned by shape, staining, and the presence of cristae within a continuous organelle) outer membrane were manually segmented in IMOD by manual tracing using the Drawing Tools plugin created by Andrew Noske. The GUNC region was defined as all nuclear envelope outside the opening of the PSMs. If a serial section was missing or unusable, the Interpolator plugin created by Andrew Noske was used to approximate any contours in the missing slice. Model movies were generated within FIJI using TIFF series exported from 3DMOD.

The degree of membrane contact was quantified in MATLAB from IMOD-generated models. Briefly, models were exported for each “organelle” (either all NE-bound compartments, specifically the GUNC, or mitochondria) as geometric wavefront objects. Then, the surface of the entire nuclear envelope (or specifically the GUNC) was thresholded by proximity to mitochondrial surface, using two distance thresholds of 30nm and 80nm (script provided as Supplemental File 1). The percent nuclear surface within each threshold was calculated by dividing the area within each threshold by the total nuclear surface area. As mitochondrial network size was in-excess relative to exposed nuclear envelope and mitochondrial network morphology varied considerably from cell-to-cell, we did not consider the reciprocal analysis very informative. We found no discernable relationship between the degree of mito-GUNC contact and either the mitochondrial surface area, total mitochondrial volume, or GUNC-to-total NE ratios. Therefore, differences in these characteristics do not account for differences in the degree of mitochondrial-nuclear contact between genotypes.

### Image Preparation, Quantification, and Statistics

All image analysis was performed in FIJI (RRID:SCR_002285 ^118^), described further below. Relevant scripts used for quantification are included as Supplementary Files 3 - 5. Images were deconvolved using DeltaVision SoftWoRx (GE Healthcare). XY drift in timelapse microscopy was corrected using the FIJI Registration plugin, as needed. Maximum z-projections of deconvolved images are shown for each image and were modified for presentation using linear brightness and contrast adjustments in FIJI, unless otherwise indicated. In Figure 1b, a decolvolution edge artifact was present underneath the cell-of-interest. To maintain visual clarity, this region was replaced with sharpened pre-deconvolution data for the same region; however, a slight discrepancy with the background signal remains. Only cells forming mature tetrads or four closed PSMs were quantified, with the exception of genotypes that blocked spore formation.

Nucleoporin inheritance (“Inheritance Index”) was quantified as follows, from timelapse microscopy of single cells. First, Htb1-mcherry signal was used to generate binary masks of nucleoplasmic area within time-lapse microcopy z-stacks (see Supplementary File 3). Next, individual masks of nucleoplasm midsections were chosen for pre-division (mononucleate, just prior to MI) and post-division (immediately following PSM closure, for all four spore nuclei) timepoints for each cell. Then, the user-selected masks were used to generate nuclear periphery ROIs by automated dilation and conversion to a band (see Supplementary File 4). Background was subtracted using the rolling ball method with a 50 pixel radius, then the signal intensity and area was measured for each ROI. An average was calculated from the signal intensity measurements of the four spore nuclei, then this was divided by the measured value of the pre-division nucleus. We did not find difference in ROI area, duration between pre-division and post-division timepoints, or division timing (which could impact measured inheritance due to increased photobleaching) to contribute to differences between genotypes.

To assess particle size or GEMs or age-associated nuclear protein aggregates (marked by Hsp104), a similar strategy was employed. ROIs were selected in maximum intensity z-projections timelapse microscopy for single cells mid meiosis II (assessed by Htb1-mCherry or 3xmCherry-NLS_SV40_). Background was subtracted (as above), contrast settings were matched, and binary masks were then generated for particles in each image. The area and intensity of the largest continuous particle was measured for each cell, in order to prevent contribution from signal artifacts or other cells (see Supplementary File 5). Essentially all cells had a single GEM or aggregate at the time of interest.

The perimeters of gamete PSMs were quantified using the ellipse tool from maximum intensity z-projections immediately upon closure, as assessed by their rounding and a redistribution of gamete nuclei (Htb1-mCherry). The diameter of PSM openings were measured using the line tool to trace the longest aspect of leading edge complex rings (Don1-GFP) in mid-late anaphase II, two frames (30 minutes) prior to PSM closure.

### Immunoblotting

To prepare protein samples for immunoblotting, trichloroacetic acid (TCA) extraction was performed as previously described. For each sample, 3.33 OD_600_ of cells were fixed with 5% TCA for ≥1 hour in the dark at 4°C. Cells were washed first with 750 µL chilled 50 mM Tris pH 7.5 1 mM EDTA, then 750 µL mL chilled acetone, and the resulting pellets were allowed to completely. Glass beads (∼100 µL) and 100 µL chilled protein breakage buffer (50 mM Tris pH 7.5, 1 mM EDTA, 3 mM DTT, 1X cOmplete EDTA-free inhibitor cocktail Roche) were added to the samples, which were then lysed for 5 minutes using the Mini-Beadbeater-96 (BioSpec). 50 µL of 3x SDS sample buffer (250 mM Tris pH 6.8, 8% β-mercaptoethanol, 40% glycerol, 12% SDS, 0.00067% bromophenol blue) was added to the samples, which were then flash-frozen and stored at −20°C. Samples were denatured at 70°C for ten minutes and cleared of insoluble material by centrifugation prior to electrophoresis.

To visualize the proteins, the samples were then subjected to polyacrylamide gel electrophoresis. Samples were run on 4–12% Bis-Tris Bolt gels (Thermo Fisher) at 120V for approximately 90 minutes, along with the Precision Plus Protein Dual Color Standards (Bio-Rad) ladder to determine protein size. Transfer to a nitrocellulose membrane (0.45 μm, Bio-Rad) was performed in a Trans-Blot Turbo System (Bio-Rad), using the 30 min mixed molecular weight protocol. For proteins above a molecular weight of around 100 kDa, a wet transfer was instead performed at 4°C in transfer buffer (25 mM Tris, 195 mM glycine, and 15% methanol), using a Mini-PROTEAN Tetra tank (Bio-Rad) run at 180 mA for 3 h. Membranes were blocked in PBS Intercept Blocking Buffer (LI-COR Biosciences). Blots were incubated overnight in primary antibody diluted in PBS Intercept Blocking Buffer at 4°C. The following primary antibodies were used: 1:2,000 mouse anti-GFP (RRID:AB_2313808, 632381, Clontech); and 1:2,000 mouse anti-3V5 (RRID:AB_2556564, Invitrogen); 1:10,000 rabbit anti-hexokinase (RRID:AB_219918, 100-4159, Rockland). Blots were then washed in PBST (PBS with 0.1% Tween-20) and incubated in secondary antibody diluted in PBS Intercept Blocking Buffer + 0.1% Tween-20 for 3-6 h at RT. The following secondary antibodies were used: 1:10,000 goat anti-mouse conjugated to IRDye 800CW (RRID:AB_621847, 926–32212, LI-COR Biosciences) and 1:10,000 goat anti-rabbit conjugated to IRDye 680RD (RRID:AB_10956166, 926–68071, LI-COR Biosciences). Blots were then washed in PBS-T, rinsed with PBS, and imaged using the Odyssey CLx system (LI-COR Biosciences). Image analysis and quantification of blots was performed in FIJI (RRID:SCR_002285 ^118^).

## Data availability

All the reagents used in this study are available upon request.

## Supporting information

Supplementary Table 1 (Strains)

Supplementary Table 2 (Plasmids)

Supplementary Table 3 (Primers)

Supplementary Table 4 (Imaging Conditions)

Supplementary Files S1-S4 (Scripts)

## Acknowledgements

We thank Jeremy Thorner, Michael Rape, Samantha Lewis, James Olzmann, Tianyao Xiao, Jessica Leslie, Tina Sing, Costa Bartolutti, Alena Bishop, Him Visitvoranat, and the entire Brar-Ünal Lab for comments on this manuscript and for valuable discussions. This work was supported by funds from the National Institutes of Health (R01AG071801) and Astera Institute to EÜ; funds from NIH (R01AG071869 and R35GM134886) and Astera Institute to G.A.B.; an NIH Traineeship (T32 GM007232) to C.T.R., B.S.S., G.A.K., and M.E.W.; an NIH Traineeship (T32 GM132022) to C.T.R. and M.E.W.; an NIH Traineeship (T32 GM007127) and National Science Foundation Graduate Research Fellowship (DGE 1752814; DGE 1106400) to E.M.S and (DGE 1752814; DGE 2146752) to G.A.K, and M.E.W; and a Swiss National Science Foundation mobility fellowship (P500PB_214421) to S.S. The content is solely the responsibility of the authors and does not necessarily represent the official views of the National Institute of Health. We thank Tianyao Xiao, who conducted the genome-wide nucGEM screen alongside Ben Styler and will be an author on the future resulting paper. We thank staff at the UC Berkeley Electron Microscopy Lab for assistance in electron microscopy data collection. We acknowledge technical support from Alena Bishop, Kirsten Aebgayle Tomas, Hannah Chen, Christiane Brune, and Jack Wang.

## Author contributions

C.T.R, B.S.S, E.M.S, and E.Ü. designed the research. C.T.R, B.S.S, E.M.S, G.A.K, S.S, and performed the experiments. C.T.R and E.M.S performed the image analysis. C.T.R and D.M.J performed the electron microscopy sample preparation. C.T.R and E.Ü wrote the original draft of the manuscript. All authors edited and revised the final manuscript.

## Extended Data Figure Legends

**Figure S2 Extended Data.**
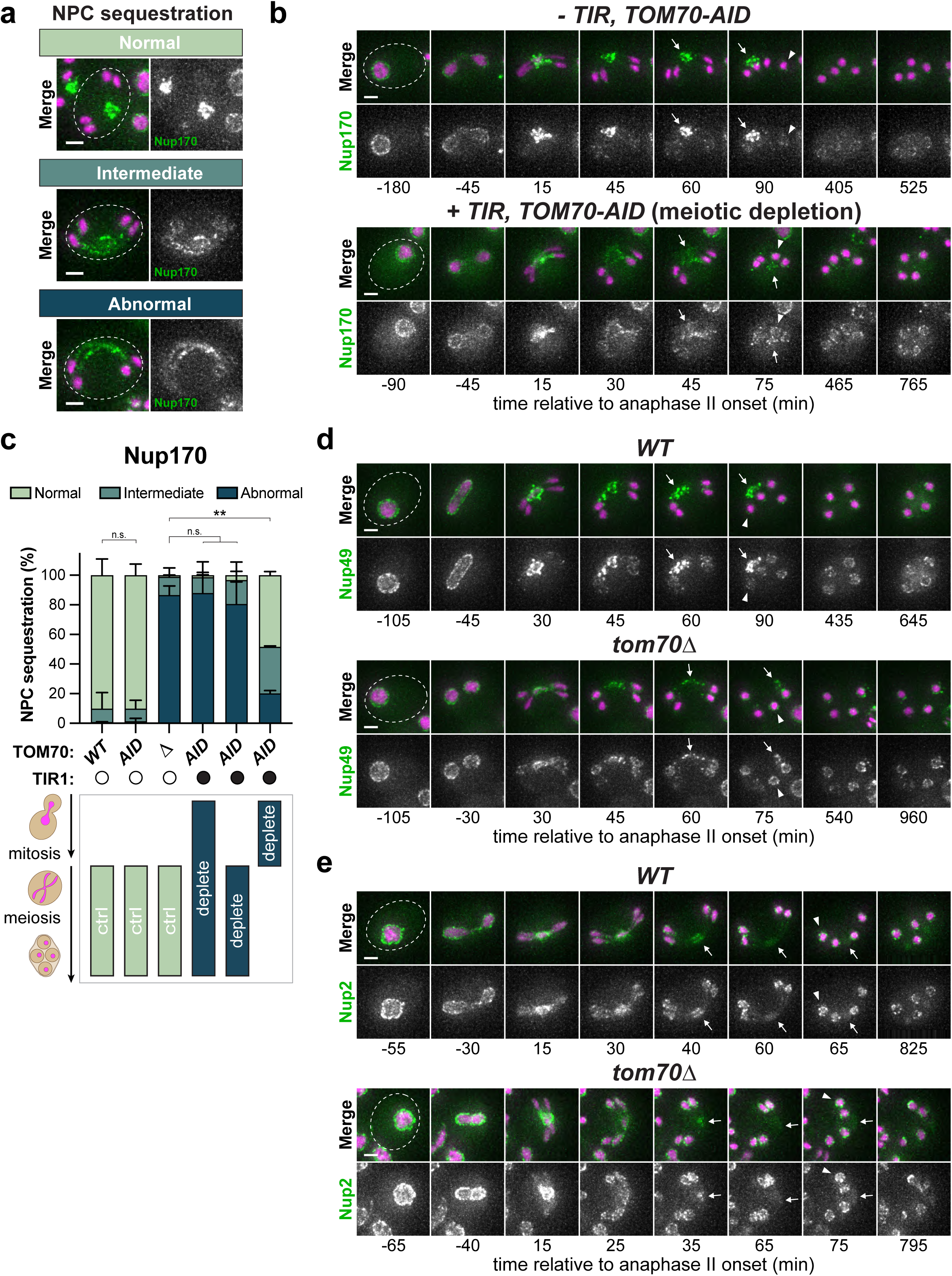
**a,** Representative cells for normal, intermediate, and abnormal nucleoporin sequestration categories (e.g. pertaining to Fig. 2c-d and C, below). Shown are single timepoint images of sporulating cells with Nup170-GFP and Htb1-mCherry (H2B), 45 min after the onset of anaphase II. **b,** Montage of sporulating Tom70-AID cells expressing Nup170-GFP and Htb1-mCherry (H2B), either with or without the *TIR1^F74G^* transgene. 100 µM 5-Ph-IAA auxin and 50 µM CuSO_4_ were added at 0h in SPO, resulting in meiosis-specific depletion of Tom70 in *+TIR* cells, but not in *-TIR* cells (see Extended Data Figure 4e for assessment of protein depletion). Arrows indicate nucleoporin signal accumulation at the GUNC, arrowheads highlight the nucleoporin signal at spore nuclei upon PSM closure. **c,** Quantification of abnormal nucleoporin sequestration (Nup170-GFP) upon depletion of Tom70-AID continually, only during sporulation, or only prior to meiosis. Wild type and *tom70Δ* mutants serve as controls for either complete presence or absence of Tom70 function. Shown is the average ± standard deviation from two experiments, n = 60-154 cells each. ** (p<0.01), two-tailed unpaired t-test. Constitutive and meiosis-specific depletion of Tom70 phenocopied *tom70Δ*, while depletion during mitotic precultures resulted in only a modest defect (p = 0.004, vs *WT*). Notably, Tom70-AID levels under this later condition only recovered to 20% of normal levels (observed in *-TIR, Tom70-AID* cells) after 6h in SPO, after which meiotic divisions begin (data not shown). **d,** Montages of sporulating wild-type and *tom70Δ* cells expressing Nup49-GFP, an FG-repeat containing channel nucleoporin, and Htb1-mCherry (H2B). Arrows indicate nucleoporin signal accumulation at the GUNC, arrowheads highlight the nucleoporin signal at spore nuclei upon PSM closure. **e,** Montages of sporulating wild-type and *tom70Δ* cells expressing Nup2-GFP, a basket nucleoporin, and Htb1-mCherry (H2B). Note that basket nucleoporins detach from the core NPC and are not sequestered to the GUNC in wildtype cells^21,38^. Arrows indicate GUNC region, largely devoid of basket nucleoporins, while arrowheads highlight the nucleoporin signal at spore nuclei upon PSM closure.

**Figure S4 Extended Data.**
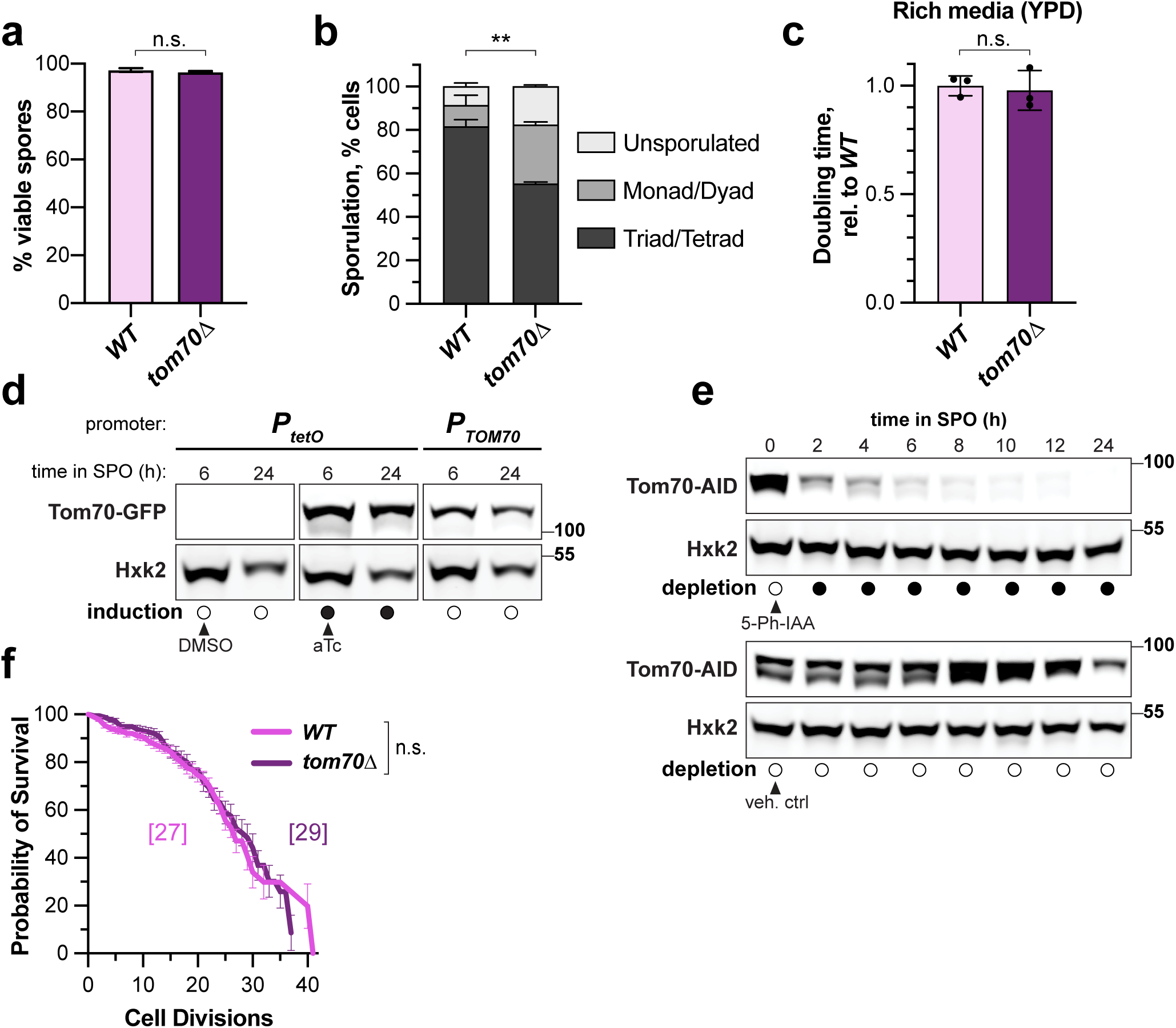
**a,** Gamete viability quantification for wild type and *tom70Δ* mutants, showing percent gametes forming visible colonies. Shown is average ± standard deviation for 2 experiments, n = 96 cells each. n.s. (p = 0.31), two-tailed unpaired t-test. **b,** Sporulation efficiency, assessed by percentage of wild-type and *tom70Δ* mutant cells forming 3-4 spores (triad/tetrad), 1-2 spores (monad/dyad), or no spores (unsporulated) after 24h in SPO at 30°C. Shown is average ± SD for 2 experiments, n = 300 cells each. ** (p<0.01), two-tailed unpaired t-test of percent triad/tetrads. **c,** Relative doubling time of *tom70Δ* mutant cells relative to wild-type cells during log-phase growth in liquid rich media (YPD) at 30°C. n.s. (p = 0.70); two-tailed Mann-Whitney U test. **d,** Western blot of Tom70-GFP levels, either under control of the endogenous *P_TOM70_* promoter or an inducible *P_tetO_* promoter at the endogenous locus, during meiosis. Addition of 5 µg/mL aTc at 0h in SPO results in induction of approximately twice the endogenous Tom70-GFP levels at 6h, just prior to onset of meiotic divisions. In contrast, Tom70-GFP is not expressed when aTc is withheld (DMSO, vehicle control). Hxk2 serves as a loading control. **e,** Western blot of Tom70-AID depletion during meiosis. 100 µM 5-Ph-IAA auxin and 50 µM CuSO_4_ (to induce expression of *TIR1^F74G^*) were added at 0h in SPO, resulting in meiosis-specific depletion of Tom70 in *+TIR* cells (top), but not in *-TIR* cells. The Tom70-AID allele was detected via a V5 epitope tag, Hxk2 served as a loading control. **f,** Replicative lifespan was determined for unaged vegetatively grown wild type and *tom70Δ* haploid cells, in a manner similar as for gametes (see Methods; ^42^). Shown is the Kaplan-Meier plot of a single biological experiment ± standard error of the mean, n = 67-100 cells total per genotype. n.s. (p = 0.69), logrank Mantel-Cox test. We did not detect a reduction in replicative lifespan in haploid *tom70Δ* mutants, in contrast to a previous report^103^. This discrepancy may be due to differences in strain background, as the SK1 strain used in this study exhibits enhanced mitochondrial function relative to BY4741-derived strains through determined genetic variations ^104^.

**Figure S5 Extended Data.**
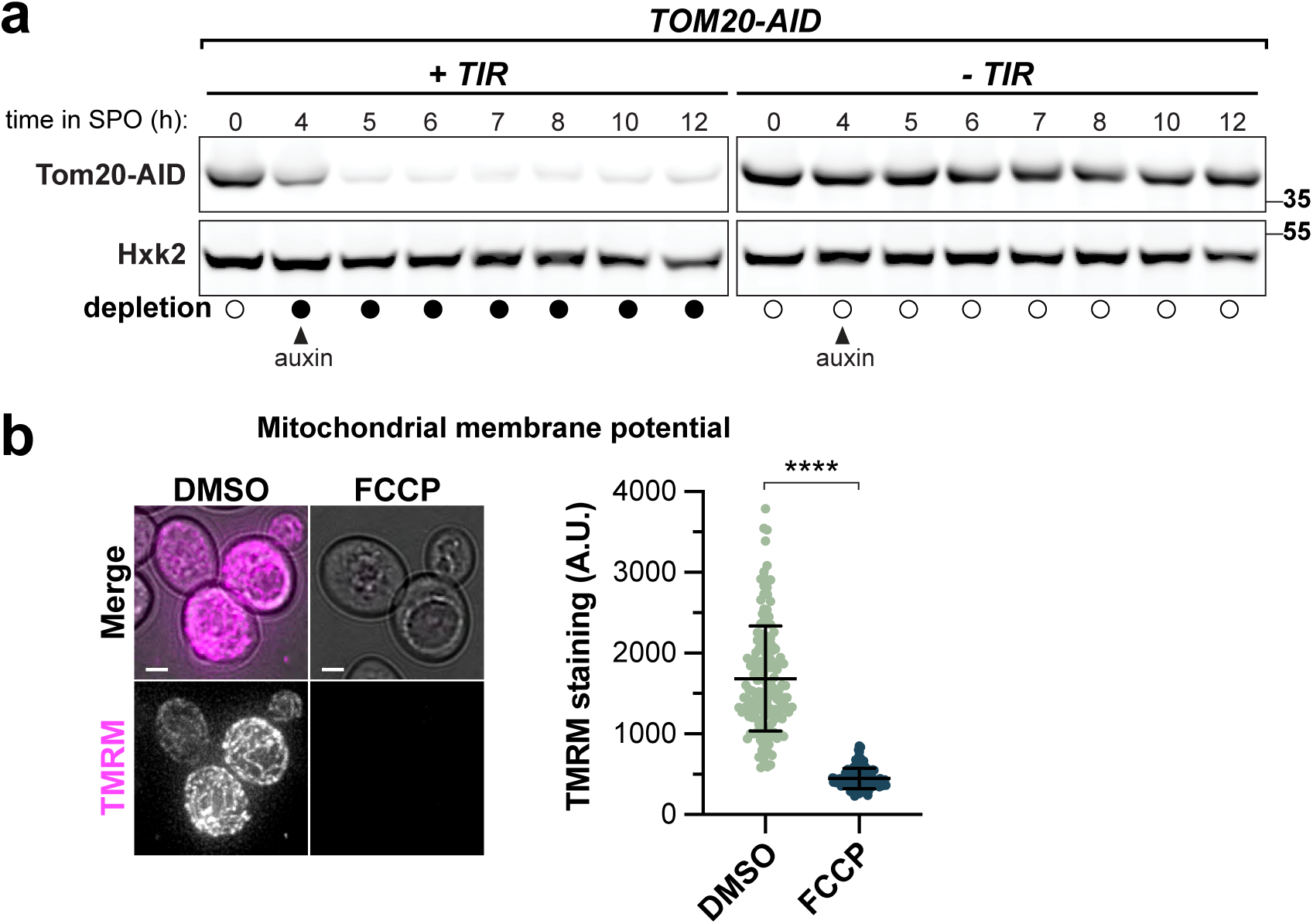
**a,** Western blot of Tom20-AID depletion during meiosis. 50 µM CuSO_4_ was added at 2.5h in SPO to induce expression of TIR1^F74G^ (left), then 1mM NAA auxin was added at 4h in SPO to deplete Tom20-AID prior to onset of meiotic divisions (6h). *-TIR* cells, which do not respond to auxin addition, serve as negative controls for depletion (right). The Tom20-AID allele was detected via a V5 epitope tag. Hxk2 served as a loading control. **b,** Sporulating wild-type cells treated with either mitochondrial uncoupling agent FCCP (50 µM) or vehicle control (DMSO) at 4h in SPO were subject to TMRM (tetramethylrhodamine methyl ester) staining after 30 minutes. Accumulation of the TMRM dye is dependent on mitochondrial membrane potential. Representative images (left) of TMRM stained meiotic cells ± FCCP treatment, along with quantification (right) of mean whole-cell TMRM intensity in single cells. Bars indicate average ± standard deviation from a single experiment, n = 113-154 cells. Fifteen highly TMRM-stained cells in the DMSO group were considered outliers (via ROUT method) and were excluded from the displayed plot and analysis. **** (p<0.0001), Kolmogorov-Smirnov test. Scale bars, 2 µm.

**Figure S6 Extended Data.**
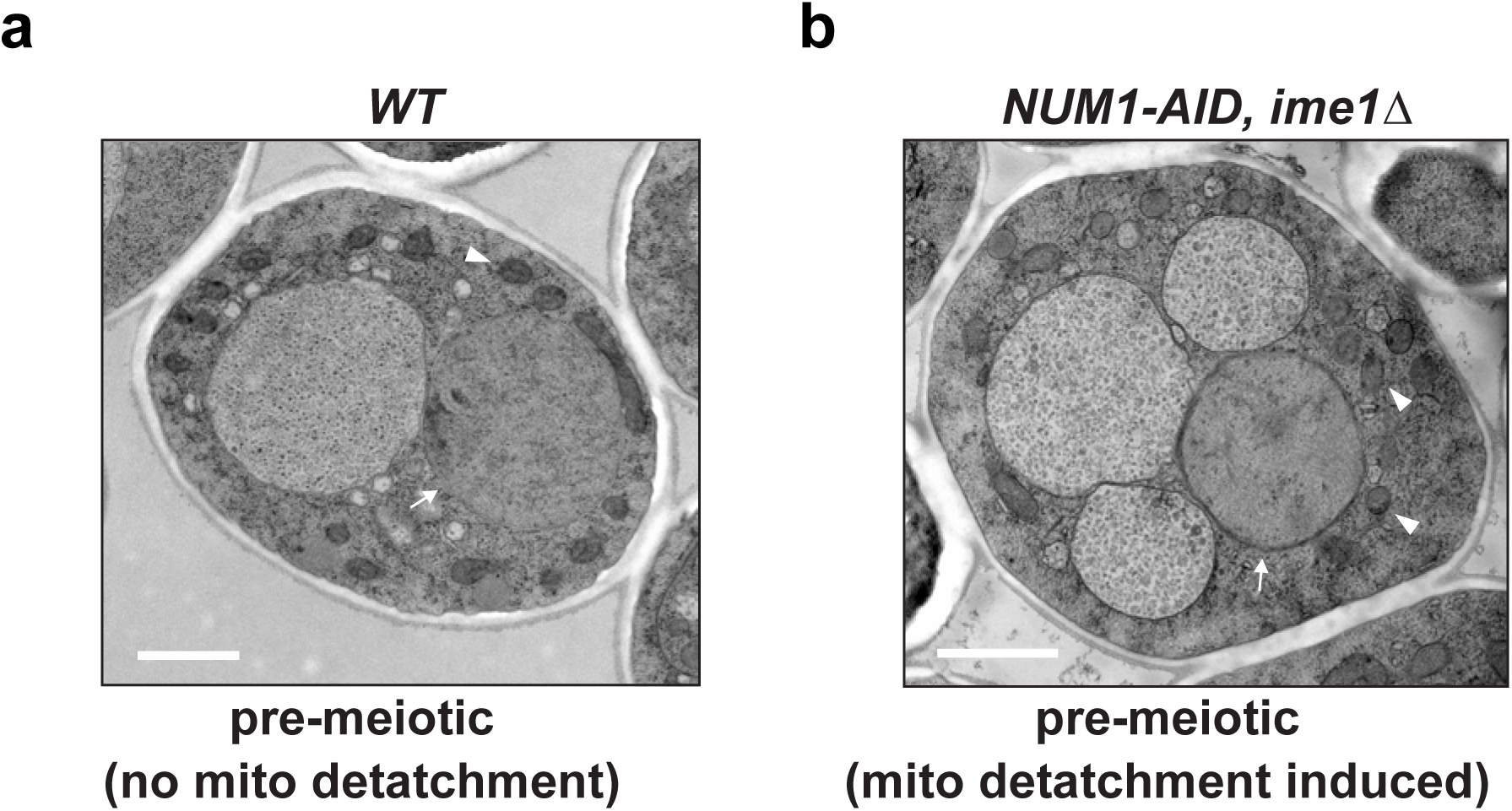
**a,** Transmission electron microscopy (TEM) of pre-division wild-type cells at 7.5h in SPO. **b,** TEM of *ime1Δ* mutant cells, which fail to undergo meiotic entry, at 6.5h SPO upon depletion of cortical mitochondrial tethering protein Num1. Num1-AID was depleted by addition of 50 µM CuSO_4_ at 1.5h in SPO to induce expression of *TIR1*, followed by addition of 500 µM I3AA auxin at 3h. Arrows emphasize the nuclear envelope within each mononucleate cell, while arrowheads indicate mitochondria. Scale bars, 1 µm.

**Figure S7 Extended Data.**
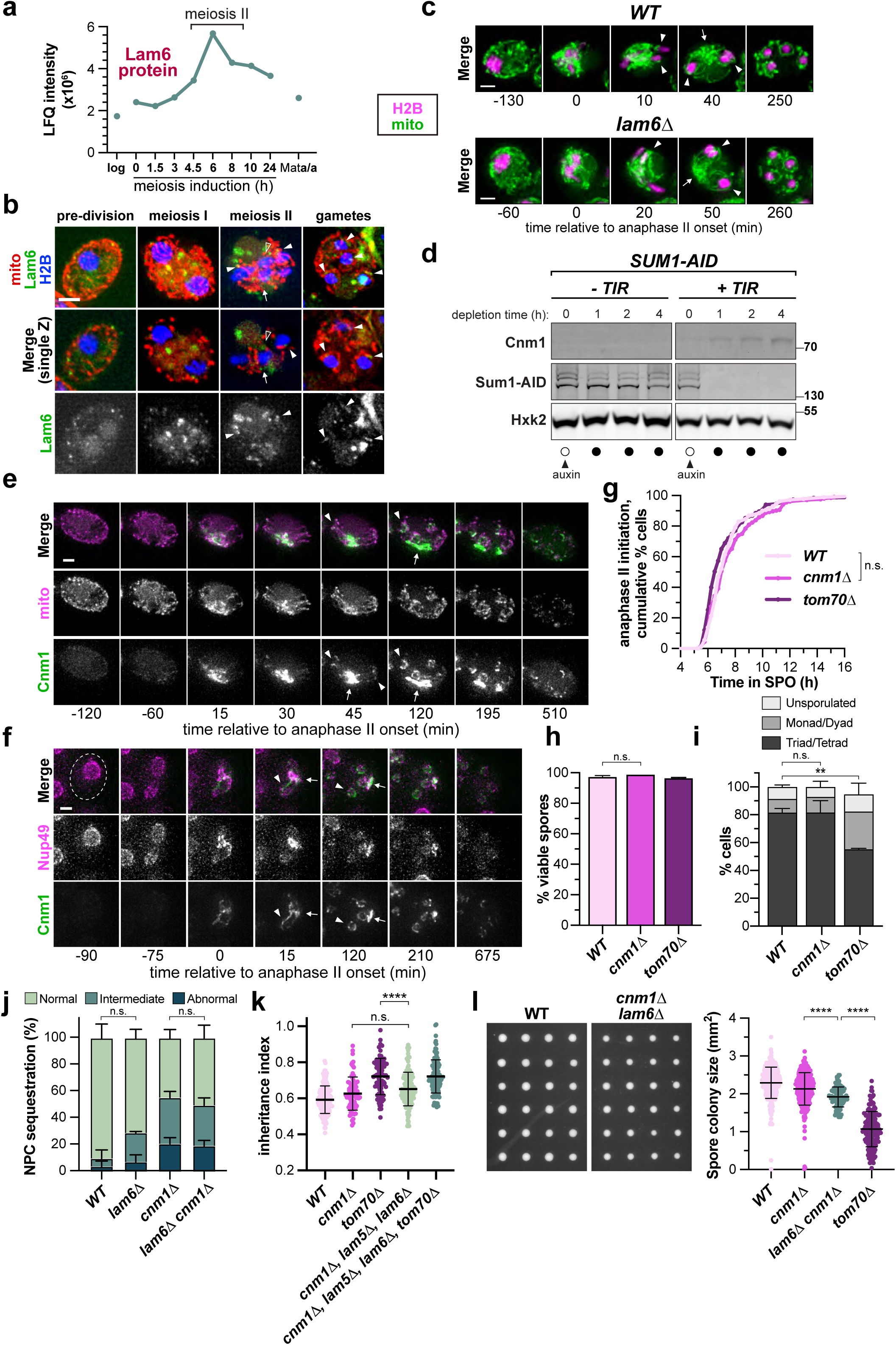
**a,** Lam6 proteins levels during log-phase mitotic growth in YPD (log), the indicated time in sporulation media (h SPO), or in control non-sporulating diploids (Mat**a**/**a**) plotted from previously generated meiotic mass spec dataset^66^. Time window corresponding to execution of meiosis II by cells within the semi-synchronized population is indicated by brackets. **b,** Images of Lam6-GFP in fixed cells of the indicated stage, shown alongside mitoBFP-marked mitochondria (mito) and Htb1-mCherry-marked chromatin (H2B). Arrowheads indicate juxtanuclear Lam6-GFP puncta by mitochondria, arrows indicate signal at GUNC region, and open arrowheads indicate signal at nuclear-vacuole junctions. **c,** Montages of sporulating wild-type or *lam6Δ* mutants tracking Cit1-GFP-marked mitochondria (mito) alongside Htb1-mCherry-marked chromatin (H2B). Arrows mark mitochondrial enrichment at the GUNC, while arrowheads emphasize mitochondrial proximity at gamete nuclei. **d,** Western blotting of endogenously-tagged Cnm1-GFP upon depletion of Sum1, a transcriptional repressor of mid-meiotic genes, by auxin induced degradation. Sum1-AID was depleted upon addition of 50 µM CuSO_4_ (to induce expression of *TIR1)* and 500 µM I3AA auxin to log-phase cultures. Sum1-AID levels were detected via a V5 epitope. Hxk2 serves as a loading control. **e, f,** Montages of sporulating cells with Cnm1-GFP alongside either (e) Cit1-mCardinal-marked mitochondria (mito), or (f) Nup49-mCherry. Arrows emphasize accumulation at the GUNC region, while arrowheads mark Cnm1 enrichment along segregating mitochondrial tubules and nuclear envelope lobes, while arrowheads emphasize Cnm1 localization at gamete nuclei. **g,** Quantification of cumulative percent cells initiating anaphase II over time, assessed by timelapse microscopy of chromatin (Htb1-mCherry). Shown is a representative of two experiments, n = 204 *cnm1Δ* cells. Wild type and *tom70Δ* data are reproduced from Fig 4a. n.s. (p = 0.23), logrank Mantel-Cox test. **h,** Gamete viability quantification for *cnm1Δ* mutants, showing percent gametes forming visible colonies. Shown is average ± standard deviation for 2 experiments, n = 96 cells each. Wild type and *tom70Δ* data are reproduced from Extended Data Fig 4a. n.s. (p = 0.10), two-tailed unpaired t-test. **I,** Sporulation efficiency, assessed by percentage of *cnm1Δ* mutant cells forming 3-4 spores (triad/tetrad), 1-2 spores (monad/dyad), or no spores (unsporulated) after 24h in SPO at 30°C. Shown is average ± SD for 2 experiments, n = 300 cells each. Wild type and *tom70Δ* data are reproduced from Fig 4a. ** (p<0.01), n.s. (p = 0.98); two-tailed unpaired t-test of percent triad/tetrads. **j,** Quantification of abnormal nucleoporin (NPC) sequestration in *lam6Δ* single mutants and *cnm1Δ lam6Δ* double mutants, with wild type and *cnm1Δ* data replotted from Fig. 7i. Shown is the average ± standard deviation from two experiments, n = 131-270 cells each. *WT* vs *lam6Δ* (p = 0.55), *cnm1Δ* vs *lam6Δ cnm1Δ* (p = 0.73); two-tailed unpaired t-test of abnormal sequestration category. **k**, Inheritance index quantification of Nup170-GFP for *lam6Δ lam5Δ cnm1Δ* triple mutants is plotted against wildtype, *cnm1Δ,* and *tom70Δ* (data from Fig. 7j). *LAM5* is a paralog of *LAM6*. The average ± standard deviation from a single experiment is shown, n = 119. Only cells forming four closed spores that remained intact through the entire 18h movie were quantified. *cnm1Δ* vs *lam6Δ lam5Δ cnm1Δ* (p = 0.08), *tom70Δ* vs *tom70Δ lam6Δ lam5Δ cnm1Δ* (p = 0.50),; Kolmogorov-Smirnov test. **l,** Representative image (left) of colonies grown for 48h on YPD at 30°C from individual gametes of the indicated genotype, with quantification (right). Individual colony sizes from a single experiment (*lam6Δ cnm1Δ*) or at least 2 experiments (remaining genotypes) are plotted, n = 96-286 cells each. **** (p<0.0001), Kolmogorov-Smirnov test. The strains used in this experiment lacked fluorescent markers, while previously shown spore colony growth experiments were performed in an *HTB1-mCherry* and *NUP170-GFP* background. Scale bars, 2 µm.

**Figure S8 Extended Data.**
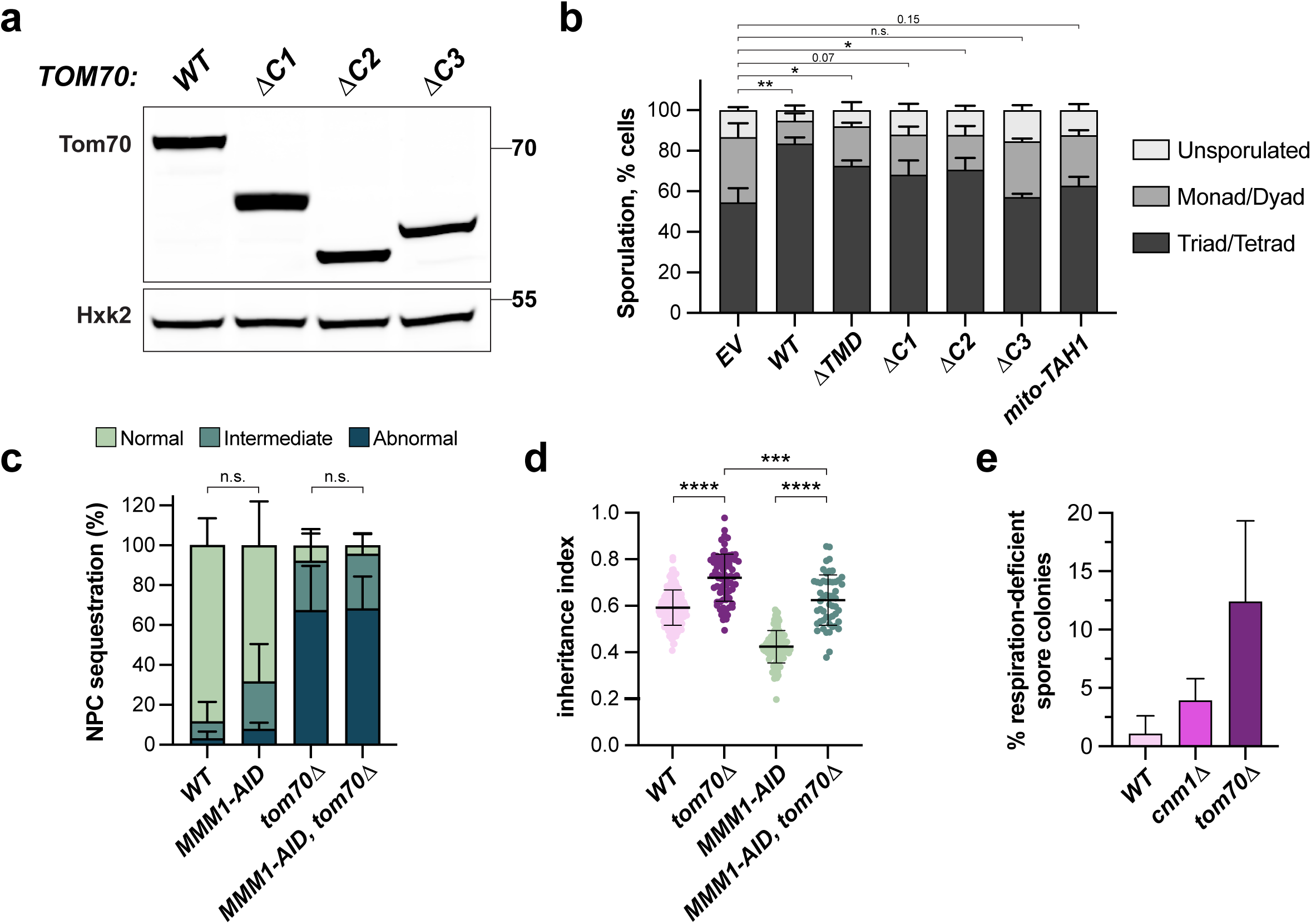
**a,** Western blot of Tom70 wild-type (*WT*) and truncation allele (*ΔC1*, *ΔC2*, or *ΔC3*) protein levels during log-phase vegetative growth in YPD. Transgenes were integrated at an ectopic locus, leaving endogenous full-length Tom70 intact. Protein from Tom70 transgenes were specifically detected via a V5 epitope tag. depletion during meiosis. **b,** Sporulation efficiency quantification for Tom70 truncation mutants, assessed by percentage of wild-type and *tom70Δ* mutant cells forming 3-4 spores (triad/tetrad), 1-2 spores (monad/dyad), or no spores (unsporulated) after 24h in SPO at 30°C. Shown is average ± standard deviation for 2 experiments, n = 300 cells each. * (p<0.01), *** (p<0.001), two-tailed unpaired t-test of percent triad/tetrads. **c,** Penetrance of abnormal Nup170-GFP nucleoporin (NPC) sequestration quantified in meiotic cells depleted of ERMES (via Mmm1-AID) by addition of 50 µM CuSO_4_ at 1.5h in SPO to induce expression of *TIR1*, followed by addition of 500 µM I3AA auxin at 3h. Shown is the average ± standard deviation of two experiments, n = 54-124. *WT* vs *MMM1-AID* (p = 0.29), *tom70Δ* vs *tom70Δ MMM1-AID* (p = 0.97); two-tailed unpaired t-test of abnormal sequestration category. **d,** Quantified inheritance index for Nup170-GFP in individual Mmm1-AID depleted cells. Bars indicate average ± standard deviation from a single experiment, n = 142-52. **** (p<0.0001), *** (p<0.001); Kolmogorov-Smirnov test. **e,** Quantification of percent gamete colonies that are respiration-deficient (assessed by failure to grow on a non-fermentable carbon source, YPG), following 48h growth at 30°C on YPD after separation. Shown is the average ± standard deviation for 2 experiments, n = 82-92 pore colonies per genotype each. *WT* vs *cnm1Δ* (p = 0.23), *WT* vs *tom70Δ* (p = 0.17); two-tailed unpaired t-test.

## Supplementary Video Legends

**Supplementary Video 1.**

Wild-type cell from Fig. 1b expressing a fluorescent nucGEM particle (green; NLS-PfV-mTSapphire) and nuclear envelope marker (magenta; 3xmScarlet-Heh2^93–378^) progressing through meiosis and gamete maturation. Time stamps indicate time elapsed, hr:min. Scale bar, 2 µm.

**Supplementary Videos 2.**

Wild-type cell from Fig. 1c expressing a fluorescent nucGEM particle (green; NLS-PfV-mTSapphire) and chromatin marker (magenta; Htb1-mCherry) progressing through meiosis and gamete maturation. Time stamps indicate time elapsed, hr:min. Scale bar, 2 µm.

**Supplementary Videos 3.**

*tom70Δ* cell from Fig. 1f expressing a fluorescent nucGEM particle (green; NLS-PfV-mTSapphire) and chromatin marker (magenta; Htb1-mCherry) progressing through meiosis and gamete maturation. Time stamps indicate time elapsed, hr:min. Scale bar, 2 µm.

**Supplementary Video 4.**

Wild-type cell from Fig. 2a expressing inner ring nucleoporin Nup170-GFP (green) and chromatin marker (magenta; Htb1-mCherry) progressing through meiosis and gamete maturation. Time stamps indicate time elapsed, hr:min. Scale bar, 2 µm.

**Supplementary Video 5.**

Wild-type cell from Fig. 2b expressing transmembrane nucleoporin Pom34-GFP (green) and chromatin marker (magenta; Htb1-mCherry) progressing through meiosis and gamete maturation. Time stamps indicate time elapsed, hr:min. Scale bar, 2 µm.

**Supplementary Video 6.**

*tom70Δ* cell from Fig. 2a expressing inner ring nucleoporin Nup170-GFP (green) and chromatin marker (magenta; Htb1-mCherry) progressing through meiosis and gamete maturation. Time stamps indicate time elapsed, hr:min. Scale bar, 2 µm.

**Supplementary Video 7.**

*tom70Δ* cell from Fig. 2b expressing transmembrane nucleoporin Pom34-GFP (green) and chromatin marker (magenta; Htb1-mCherry) progressing through meiosis and gamete maturation. Time stamps indicate time elapsed, hr:min. Scale bar, 2 µm.

**Supplementary Video 8.**

Abnormal sequestration of inner ring nucleoporin Nup170-GFP (green) in *tom70Δ* cell, shown relative to chromatin marker (magenta; Htb1-mCherry) through meiosis and gamete maturation. Time stamps indicate time elapsed, hr:min. Scale bar, 2 µm.

**Supplementary Video 9.**

Wild-type cell from Fig. 6c expressing nuclear envelope marker (green; 3xGFP-Heh2^93–378^) alongside channel nucleoporin Nup49-2xmScarlet (magenta) progressing through meiosis. Time stamps indicate time elapsed, hr:min. Scale bar, 2 µm.

**Supplementary Video 10.**

*tom70Δ* cell from Fig. 6c expressing nuclear envelope marker (green; 3xGFP-Heh2^93–378^) alongside channel nucleoporin Nup49-2xmScarlet (magenta) progressing through meiosis. Time stamps indicate time elapsed, hr:min. Scale bar, 2 µm.

**Supplementary Video 11.**

Whole-cell reconstruction from transmission electron microscopy of a wild-type cell mid-meiosis II, as shown in Fig. 6f. The nuclear envelope is shown in magenta and mitochondria are shaded gray. Portions of the nuclear envelope within 30 nm of mitochondria are colored green, and those within 80 nm are colored yellow.

**Supplementary Video 12.**

Whole-cell reconstruction from transmission electron microscopy of a wild-type cell in mid-meiosis II, as shown in Fig. 6f. Nuclear compartments (magenta), gamete plasma membranes (PSMs; transparent red), mitochondrial network (green), and vacuole (transparent cyan) are segmented. Scale bar, 4 µm.

**Supplementary Video 13.**

Transmission electron microscopy image stack of reconstructed wild-type meiotic cell from Fig. 6f and Supplementary Videos 11 & 12. of a wild-type cell in mid-meiosis II. Movie progresses through whole-cell, then backtracks through cell with nuclear compartments and mitochondria overlayed in magenta and green, respectively. Gamete plasma membranes (PSMs) and the vacuole are outlined in thin red and cyan. Scale bar, 2 µm.

**Supplementary Video 14.**

Whole-cell reconstruction from transmission electron microscopy of a wild-type cell in late meiosis II. The nuclear envelope is shown in magenta and mitochondria are shaded gray. Portions of the nuclear envelope within 30 nm of mitochondria are colored green, and those within 80 nm are colored yellow.

**Supplementary Video 15.**

Whole-cell reconstruction from transmission electron microscopy of a wild-type cell mid-meiosis II. Nuclear compartments (magenta), gamete plasma membranes (PSMs; transparent red), mitochondrial network (green), and vacuole (transparent cyan) are segmented. Scale bar, 2 µm.

**Supplementary Video 16.**

Wild-type cell from 7d expressing Cnm1-GFP (green) alongside chromatin marker (magenta; Htb1-mCherry) progressing through meiosis. Time stamps indicate time elapsed, hr:min. Scale bar, 2 µm.

**Supplementary Video 17.**

Wild-type cell from 7e expressing Cnm1-GFP (green) alongside gamete plasma membrane marker (magenta; mKate-Spo20^51–91^) progressing through meiosis. Time stamps indicate time elapsed, hr:min. Scale bar, 2 µm.

**Supplementary Video 18.**

*cnm1Δ* cell from 7h expressing inner ring nucleoporin Nup170-GFP (green) and chromatin marker (magenta; Htb1-mCherry) progressing through meiosis and gamete maturation. Time stamps indicate time elapsed, hr:min. Scale bar, 2 µm.

**Supplementary Video 19.**

Abnormal sequestration of inner ring nucleoporin Nup170-GFP (green) in *cnm1Δ* cell, shown relative to chromatin marker (magenta; Htb1-mCherry) through meiosis and gamete maturation. Time stamps indicate time elapsed, hr:min. Scale bar, 2 µm.

**Supplementary Video 20.**

Wild-type cell from Fig. 8d expressing nuclear envelope marker (green; 3xGFP-Heh2^93–378^) alongside outer mitochondrial membrane marker (magenta; Tom20^1–42^-2xmScarlet) progressing through meiosis. Time stamps indicate time elapsed, hr:min. Scale bar, 2 µm.

**Supplementary Video 21.**

*cnm1Δ* mutant cell from Fig. 8d expressing nuclear envelope marker (green; 3xGFP-Heh2^93–378^) alongside outer mitochondrial membrane marker (magenta; Tom20^1–42^-2xmScarlet) progressing through meiosis. Time stamps indicate time elapsed, hr:min. Scale bar, 2 µm.

**Supplementary Video 22.**

*tom70Δ* mutant cell from Fig. 8d expressing nuclear envelope marker (green; 3xGFP-Heh2^93–378^) alongside outer mitochondrial membrane marker (magenta; Tom20^1–42^-2xmScarlet) progressing through meiosis. Time stamps indicate time elapsed, hr:min. Scale bar, 2 µm.

## Supplementary Information

**Table S1.** Yeast Strains.

| Strain number (yÜB) <sup>1</sup> | MAT | Genotype <sup>2,3</sup> | Source / Reference <sup>4</sup> |
| --- | --- | --- | --- |
| SK1 | Strain background | *ho::LYS2, lys2, leu2::hisG, ura3, his3::hisG, trp1::hisG* |  |
| 13 | MATa (A) | *WT* |  |
| 14 | MATalpha (α) | *WT* |  |
| 15 | MATa/alpha (diploid) | *WT* |  |
| 46381 | diploid | *leu2::pRdl1-3XmScarlet-h2NLS-L-TM::LEU2, trp1::pATG8-SV40NLS-40nmGEM-sapphire::TRP1* | this study |
| 37338 | diploid | *htb1::HTB1-mCherry::HISMx6, trp1::pATG8-SV40NLS-40nmGEM-sapphire::TRP1* | this study |
| 38272 | diploid | *spo21Δ::hphNT1, htb1::HTB1-mCherry::HISMx6, trp1::pATG8-SV40NLS-40nmGEM-sapphire::TRP1* | this study |
| 38145 | diploid | *tom70Δ::KanMX, htb1::HTB1-mCherry::HISMx6, trp1::pATG8-SV40NLS-40nmGEM-sapphire::TRP1* | this study |
| 11513 | diploid | *htb1::HTB1-mCherry::HISMx6, nup170::NUP170-GFP::KanMX* | <sup>21</sup> |
| 35069 | diploid | *tom70Δ::KanMX, htb1::HTB1-mCherry::HISMx6, nup170::NUP170-GFP::KanMX* | this study |
| 13503 | diploid | *htb1::HTB1-mCherry::HISMx6, pom34::POM34-GFP::KanMX* | <sup>21</sup> |
| 36820 | diploid | *tom70Δ::NatMX, htb1::HTB1-mCherry::HISMx6, pom34::POM34-GFP::KanMX* | this study |
| 11820 | diploid | *hsp104::HSP104-mCherry::NatMX, leu2::pATG8-link-yEGFP-SPO20(51-91)::LEU2, flo8::KanMX6* | <sup>21</sup> |
| 45577 | diploid | *tom70Δ::KanMX, hsp104::HSP104-mCherry::NatMX, leu2::pATG8-link-yEGFP-SPO20(51-91)::LEU2, flo8::KanMX6* | this study |
| 16712 | diploid | *htb1::HTB1-mCherry::HISMx6, nsr1::NSR1-GFP::KanMX, flo8* | <sup>21</sup> |
| 35507 | diploid | *tom70Δ::KanMX, htb1::HTB1-mCherry::HISMX6, nsr1::NSR1-GFP::KanMX, flo8* | this study |
| 17338 | diploid | *hta1::HTA1-mApple::HIS5, rDNA::5xtetO (heterozygous), leu2::pREC8-TetR-GFP::LEU2, flo8* | 21 |
| 35509 | diploid | *tom70Δ::KanMX, hta1::HTA1-mApple::HIS5, rDNA::5xtetO (heterozygous), leu2::pREC8-TetR-GFP::LEU2, flo8* | this study |
| 37369 | diploid | *htb1::HTB1-mCherry::HISMX6, trp1::pATG8-SV40NLS-40nmGEM-sapphire::TRP1 (heterozygous)* | this study |
| 37550 | diploid | *tom70Δ::KanMX, htb1::HTB1-mCherry::HISMX6, trp1::pATG8-SV40NLS-40nmGEM-sapphire::TRP1 (heterozygous)* | this study |
| 9724 | diploid | *htb1::HTB1-mCherry::HISMX6, hsp104::HSP104-eGFP::KanMX, flo8::KanMX6* | 21 |
| 42274 | diploid | *ste5::NatMX::pTetO7.1-STE5, ura3::pRNR2-TetR-Tup1, pTetO7.1-TetR::URA3, trp1::pPma2-PMA2-eGFP::TRP1, flo8, amn1(BY4741 allele)* | this study |
| 44258 | diploid | *tom70Δ::KanMX, ste5::NatMX::pTetO7.1-STE5, ura3::pRNR2-TetR-Tup1, pTetO7.1-TetR::URA3, trp1::pPma2-PMA2-eGFP::TRP1, flo8, amn1(BY4741 allele)* | this study |
| 41118 | diploid | *tom70::NatMX6::pTetO7.1-Tom70, ura3::pRNR2-TetR-TUP1_pTetO7.1-TetR::URA3, htb1::HTB1-mCherry::HISMX6, nup170::NUP170-GFP::KanMX* | this study |
| 12434 | diploid | *htb1::HTB1-mCherry::HISMX6, leu2::pATG8-link-yEGFP-SPO20(51-91)::LEU2* | 21 |
| 35258 | diploid | *tom70Δ::KanMX, htb1::HTB1-mCherry::HISMX6, leu2::pATG8-link-yEGFP-SPO20(51-91)::LEU2* | this study |
| 12438 | diploid | *htb1::HTB1-mCherry::HISMX6, don1::DON1-GFP::KanMX* | 21 |
| 35260 | diploid | *tom70Δ::KanMX, htb1::HTB1-mCherry::HISMX6, don1::DON1-GFP::KanMX* | this study |
| 38030 | diploid | *sam37Δ::hphNT1, htb1::HTB1-mCherry::HISMX6, nup170::NUP170-GFP::KanMX* | this study |
| 37273 | diploid | *tom20::TOM20-3V5-IAA17(71-114)::NatMX4, htb1::HTB1-mCherry::HISMX6, nup170::NUP170-GFP::KanMX* | this study |
| 37275 | diploid | *his3::pCup1-OsTIR1::HIS3, tom20::TOM20-3V5-IAA17(71-114)::NatMX4, htb1::HTB1-mCherry::HISMX6, nup170::NUP170-GFP::KanMX* | this study |
| 21933 | diploid | *htb1::HTB1-mCherry::HISMX6, tom70::TOM70-GFP::KanMX* | this study |
| 43587 | diploid | *his3::pRdl1-TOM20TMD(1-42)-3XmScarlet::HIS3, leu2::pRdl1-3xGFP-h2NLS-L-TM::LEU2* | this study |
| 41512 | diploid | *nup49::NUP49-2x mScarlet-l::cgLEU2, leu2::pRdl1-3xGFP-h2NLS-L-TM::LEU2* | this study |
| 41520 | diploid | *tom70Δ::NatMX4, nup49::NUP49-2x mScarlet-l::cgLEU2, leu2::pRdl1-3xGFP-h2NLS-L-TM::LEU2* | this study |
| 7055 | diploid | *ndt80::TRP1::pGAL-NDT80, ura3::pGPD1-GAL4(1-848)-ER::URA3, htb1::HTB1-mCherry::HISMX6 (heterozygous), his3::pGPD1-Su9MTS-eGFP::HIS3 (heterozygous)* | this study |
| 32619 | diploid | *ndt80::TRP1::pGAL-NDT80, ura3::pGPD1-GAL4(1-848)-ER::URA3, cnm1::CNM1-GFP::KanMX, cit1::CIT1-mCardinal::His3MX6* | this study |
| 32897 | diploid | *htb1::HTB1-mCherry::HISMX6, cnm1::CNM1-GFP::KanMX* | this study |
| 32895 | diploid | *leu2::pATG8-link-mKate-SPO20(51-91)::LEU2, cnm1::CNM1-GFP::KanMX* | this study |
| 33779 | diploid | *spo21Δ::hphNT1, htb1::HTB1-mCherry::HISMX6, cnm1::CNM1-GFP::KanMX* | this study |
| 33497 | diploid | *tom70Δ::KanMX, htb1::HTB1-mCherry::HISMX6, cnm1::CNM1-GFP::KanMX* | this study |
| 35262 | diploid | *cnm1Δ::hphNT1, htb1::HTB1-mCherry::HISMX6, nup170::NUP170-GFP::KanMX* | this study |
| 37255 | diploid | *trp1::EV::TRP1, tom70Δ::NatMX4, htb1::HTB1-mCherry::HISMX6, nup170::NUP170-GFP::KanMX* | this study |
| 37257 | diploid | *trp1::pTom70-TOM70::TRP1, tom70Δ::NatMX4, htb1::HTB1-* | this study |
|  |  | *mCherry::HISMx6, nup170::NUP170-GFP::KanMX* |  |
| 37259 | diploid | *trp1::pTom70-TOM70(ΔC1_99-214)::TRP1, tom70Δ::NatMX4, htb1::HTB1-mCherry::HISMx6, nup170::NUP170-GFP::KanMX* | this study |
| 37261 | diploid | *trp1::pTom70-TOM70(ΔC2_248-460)::TRP1, tom70Δ::NatMX4, htb1::HTB1-mCherry::HISMx6, nup170::NUP170-GFP::KanMX* | this study |
| 37263 | diploid | *trp1::pTom70-TOM70(ΔC3_461-617)::TRP1, tom70Δ::NatMX4, htb1::HTB1-mCherry::HISMx6, nup170::NUP170-GFP::KanMX* | this study |
| 37265 | diploid | *trp1::pTom70-TOM70TMD(1-41)-TAH1::TRP1, tom70Δ::NatMX4, htb1::HTB1-mCherry::HISMx6, nup170::NUP170-GFP::KanMX* | this study |
| 37267 | diploid | *trp1::pTom70-TOM70(ΔTMD_2-41)::TRP1, tom70Δ::NatMX4, htb1::HTB1-mCherry::HISMx6, nup170::NUP170-GFP::KanMX* | this study |
| 41865 | diploid | *trp1::EV::TRP1 (heterozygous), leu2::pGPD-CNM1-GFP::LEU2 (heterozygous), tom70Δ::NatMX4, tom71Δ::hphNT1* | this study |
| 41867 | diploid | *trp1::pTom70-TOM70::TRP1 (heterozygous), leu2::pGPD-CNM1-GFP::LEU2 (heterozygous), tom70Δ::NatMX4, tom71Δ::hphNT1* | this study |
| 41869 | diploid | *trp1::pTom70-Tom70(ΔC1_99-214)::TRP1 (heterozygous), leu2::pGPD-CNM1-GFP::LEU2 (heterozygous), tom70Δ::NatMX4, tom71Δ::hphNT1* | this study |
| 41871 | diploid | *trp1::pTom70-Tom70(ΔC2_248-460)::TRP1 (heterozygous), leu2::pGPD-CNM1-GFP::LEU2 (heterozygous), tom70Δ::NatMX4, tom71Δ::hphNT1* | this study |
| 41873 | diploid | *trp1::pTom70-Tom70(ΔC3_461-617)::TRP1 (heterozygous), leu2::pGPD-CNM1-GFP::LEU2 (heterozygous), tom70Δ::NatMX4, tom71Δ::hphNT1* | this study |
| 47202 | diploid | *trp1::pTom70-TOM70TMD(1-41)-TAH1::TRP1 (heterozygous), leu2::pGPD-CNM1-GFP::LEU2 (heterozygous), tom70Δ::NatMX4, tom71Δ::hphNT1* | this study |
| 45008 | diploid | *cnm1Δ::hphNT1, his3::pRdl1-TOM20TMD(1-42)-3XmScarlet::HIS3, leu2::pRdl1-3xGFP-h2NLS-L-TM::LEU2* | this study |
| 45010 | diploid | *tom70Δ::KanMX, his3::pRdl1-TOM20TMD(1-42)-3XmScarlet::HIS3, leu2::pRdl1-3xGFP-h2NLS-L-TM::LEU2* | this study |
| 44417 | diploid | *trp1::pTom70-Tom70(86)^linkerGS-V5-AID\*::TRP1 (heterozygous), tom70Δ::KanMX, htb1::HTB1-mCherry::NatMX4, nup170::NUP170-GFP::LEU2* | this study |
| 44225 | diploid | *his3::pCup1-OsTIR1(F74G)::HIS3, trp1::pTom70-Tom70(86)^linkerGS-V5-AID\*::TRP1 (heterozygous), tom70Δ::KanMX, htb1::HTB1-mCherry::NatMX4, nup170::NUP170-GFP::LEU2* | this study |
| 36092 | diploid | *htb1::HTB1-mCherry::HISMX6; nup49::NUP49-Knop-eGFP::KanMX* | this study |
| 36582 | diploid | *tom70Δ::NatMX4, htb1::HTB1-mCherry::HISMX6; nup49::NUP49-Knop-eGFP::KanMX* | this study |
| 15305 | diploid | *nup2::NUP2-GFP-KanMX, htb1::HTB1-mCherry::HISMX6* | 21 |
| 35071 | diploid | *tom70Δ::KanMX, nup2::NUP2-GFP-KanMX, htb1::HTB1-mCherry::HISMX6* | this study |
| 40755 | diploid | *tom70::NatMX6::pTetO7.1-TOM70-GFP::KanMX, ura3::pRNR2-TetR-TUP1_pTetO7.1-TetR::URA3* | this study |
| 39969 | diploid | *tom70::TOM70-GFP::KanMX, ura3::pRNR2-TetR-TUP1_pTetO7.1-TetR::URA3* | this study |
| 10254 | diploid | *htb1::HTB1-mCherry::HISMX6, cit1::CIT1-GFP::His3MX6* | 19 |
| 30698 | diploid | *num1::NUM1-IAA7-3V5::NatMX, ime1Δ::hphNT1, his3::pCup1-OsTIR1::HIS3, om45::OM45-mKate::His3MX6, leu2::pARO10-NVJ1(I295R, A307R, F319E)-FLAG-GFP::LEU2* | this study |
| 20119 | diploid | *htb1::HTB1-mCherry::HISMX6, lam6::LAM6-GFP::TRP1, 2μ(NatMX)_pGPD1-mito-TagBFP* | this study |
| 20463 | diploid | *lam6Δ::hphNT1, htb1::HTB1-mCherry::HISMX6, cit1::CIT1-GFP::His3MX6* | this study |
| 32393 | MATa | *sum1::SUM1-3V5-IAA7::KanMX, cit1::CIT1-mCardinal::His3MX6, cnm1::CNM1-GFP::KanMX, ura3::pGPD1-GAL4(1-848)-ER::URA3* | this study |
| 32389 | MATa | *his3::pCup1-OsTIR1::HIS3, sum1::SUM1-3V5-IAA7::KanMX, cit1::CIT1-mCardinal::His3MX6, cnm1::CNM1-GFP::KanMX, ura3::pGPD1-GAL4(1-848)-ER::URA3* | this study |
| 32893 | diploid | *cit1::CIT1-mCardinal::His3MX6, cnm1::CNM1-GFP::KanMX* | this study |
| 32901 | diploid | *nup49::NUP49-mCherry::His3MX6, cnm1::CNM1-GFP::KanMX* | this study |
| 36051 | diploid | *lam6Δ::hphNT1, htb1::HTB1-mCherry::HISMX6, nup170::NUP170-GFP::KanMX* | this study |
| 36053 | diploid | *lam6Δ::hphNT1, cnm1Δ::NatMX, htb1::HTB1-mCherry::HISMX6, nup170::NUP170-GFP::KanMX* | this study |
| 45020 | diploid | *lam5Δ::KanMX, lam6Δ::hphNT1, cnm1Δ::NatMX, htb1::HTB1-mCherry::HISMX6, nup170::Nup170-GFP::LEU2* | this study |
| 44158 | diploid | *lam5Δ::KanMX, lam6Δ::hphNT1, cnm1Δ::NatMX, tom70Δ::hphMX6, htb1::HTB1-mCherry::HISMX6, nup170::Nup170-GFP::LEU2* | this study |
| 32109 | diploid | *cnm1Δ::hphNT1* | this study |
| 33080 | diploid | *lam6Δ::hphNT1, cnm1Δ::hphNT1* | this study |
| 37171 | diploid | *tom70Δ::KanMX* | this study |
| 37472 | diploid | *trp1::pTom70-TOM70::TRP1* | this study |
| 42526 | diploid | *trp1::pTom70-TOM70(ΔC1_99-214)<sup>(51)</sup>V5::TRP1* | this study |
| 42528 | diploid | *trp1::pTom70-TOM70(ΔC2_248-460)<sup>(51)</sup>V5::TRP1* | this study |
| 42530 | diploid | *trp1::pTom70-TOM70(ΔC3_461-617)<sup>(51)</sup>V5::TRP1* | this study |
| 35517 | diploid | *his3::pCup1-OsTIR1::HIS3, mmm1::MMM1-3V5-IAA7::hphNT1, htb1::HTB1-mCherry::HISMX6, Nup170-GFP::KanMX* | this study |
| 36279 | diploid | *tom70Δ::NatMX4, his3::pCup1-OsTIR1::HIS3, mmm1::MMM1-3V5-IAA7::hphNT1, htb1::HTB1-mCherry::HISMX6, Nup170-GFP::KanMX* | this study |
| 40504 | diploid | *tom70Δ::NatMX4, htb1::HTB1-mCherry::HISMX6, nup170::NUP170-GFP::KanMX* | this study |

<sup>1</sup> Strains are listed according to Figure order.
<sup>2</sup> Alleles are separated by commas.
<sup>3</sup> All indicated alleles in diploids are homozygous, unless otherwise specified.
<sup>4</sup> Previously generated alleles are further described elsewhere (see Methods).

**Table S2.**
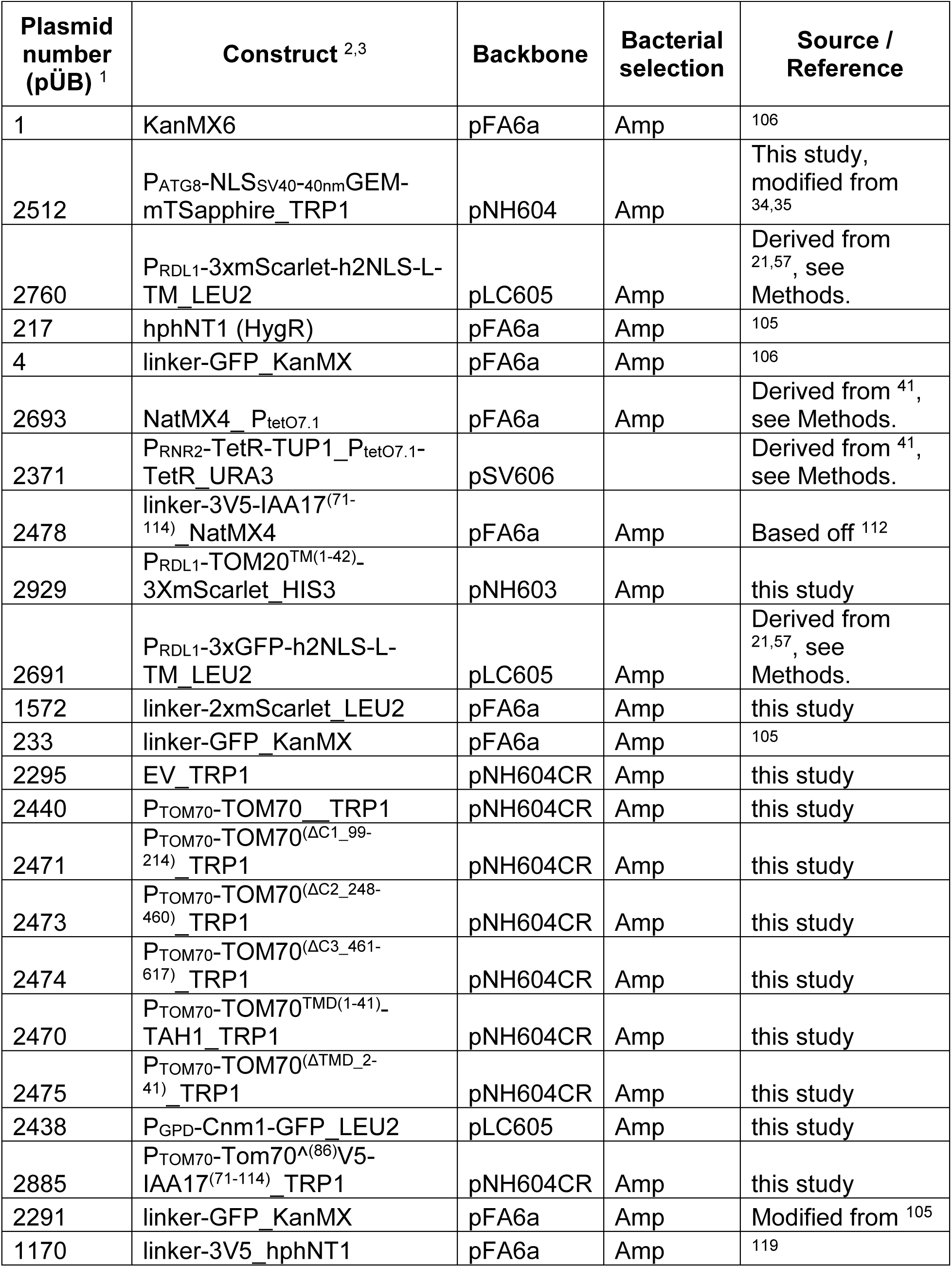
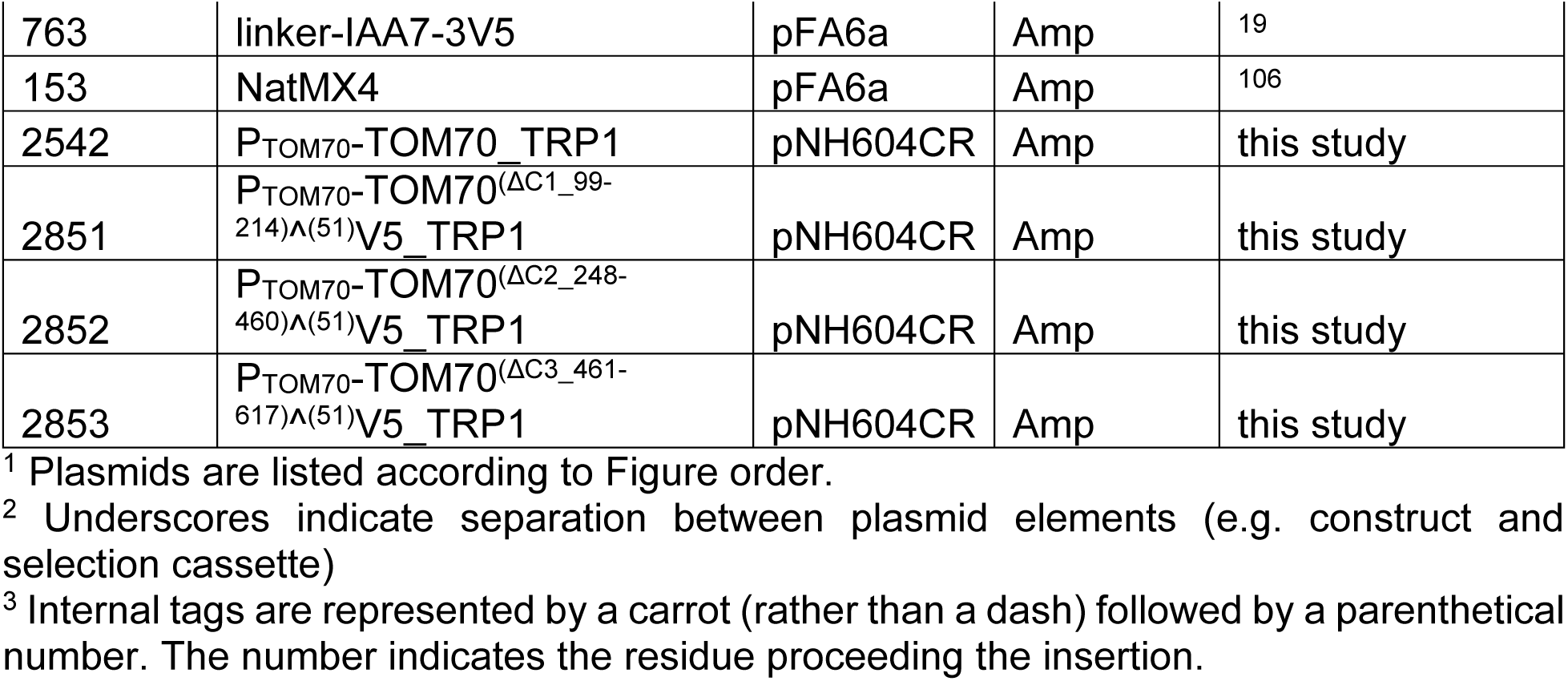
Plasmids.

**Table S3.**
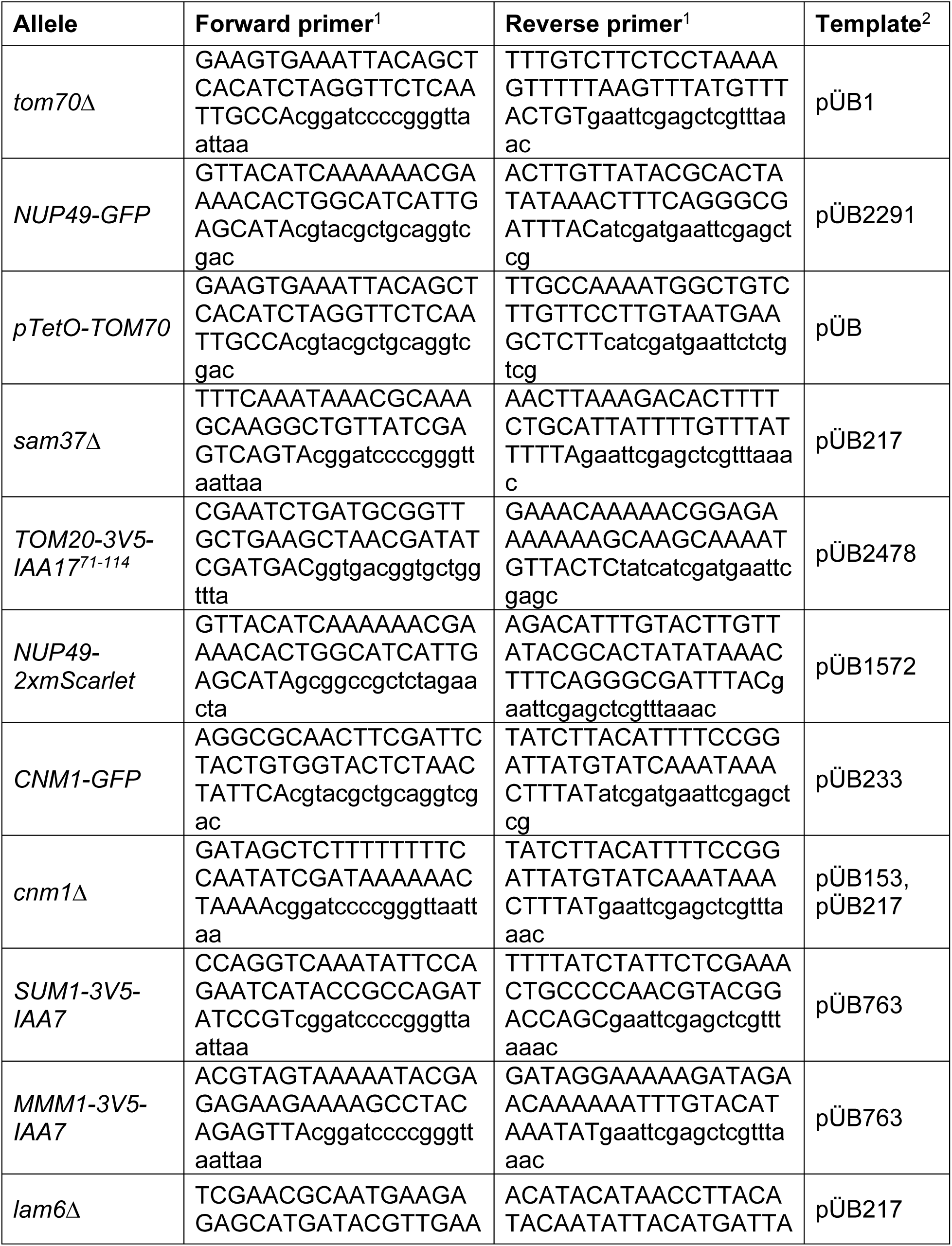
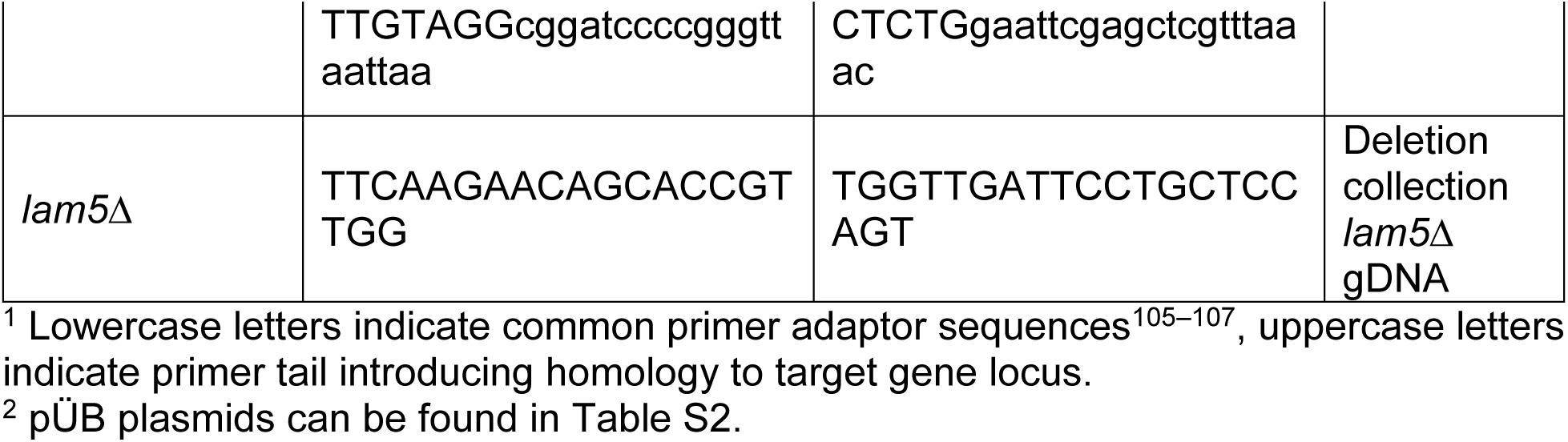
Primers.

**Table S4.** Imaging Conditions.

**Tabel S4. Imaging Conditions**
| Sub Figure | Experiment type | Exposures | Section spacing; objective | Acquisition frequency, duration |
| --- | --- | --- | --- | --- |
| 1b; | Movie, CellASIC | 25 ms, 10%T TRITC;<br>25 ms, 10%T GFP | 1 μm/slice,<br>8 total;<br>60x obj. | 15 min/frame,<br>18h total |
| 1c, 1d, 1e, 1f,<br>1g; 3f, 3g, 3h | Movie, 96-well | 25 ms, 10%T mCherry;<br>25 ms, 10%T GFP | 1 μm/slice,<br>8 total;<br>60x obj. | 15 min/frame,<br>18h total |
| 2a, 2b, 2c, 2d,<br>2f, 2g; 3d, 3e;<br>4a; 5a, 5b, 5c,<br>5d, 5e, 5f, 5g;<br>6a; 7f, 7g, 7h,<br>7i, 7j; 8b; S2a,<br>S2b S2c, S2d;<br>S7g, S7j, S7k;<br>S8c, S8d, S8e | Movie, CellASIC | 25 ms, 10%T mCherry;<br>25 ms, 10%T GFP | 1 μm/slice,<br>8 total;<br>60x obj. | 15 min/frame,<br>18h total |
| S2e | Movie, CellASIC | 25 ms, 10%T mCherry;<br>25 ms, 10%T GFP | 1 μm/slice,<br>8 total;<br>60x obj. | 5 min/frame,<br>8h total |
| 3b, 3c; 7d, 7e;<br>S7e, S7f | Movie, CellASIC | 25 ms, 32%T mCherry;<br>25 ms, 10%T GFP | 1 μm/slice,<br>8 total;<br>60x obj. | 15 min/frame,<br>18h total |
| 4c | Movie, 96-well | NA | 1 μm/slice,<br>8 total;<br>60x obj. | 5 min/frame,<br>10h total |
| 4e, S4f | Movie, RLS microfluidics | NA | 3-5 μm/slice,<br>3 total;<br>40x obj. | 15-20 min/frame,<br>72h total |
| 6b, 6c; 8d, 8e | Movie, CellASIC | 25 ms, 10%T TRITC;<br>25 ms, 10%T GFP | 1 μm/slice,<br>8 total;<br>60x obj. | 5 min/frame,<br>8h total |
| 7c | still images, fixed cells | 50 ms, 100%T mCherry;<br>25 ms, 100%T GFP;<br>10 ms, 100%T DAPI | 0.3 μm/slice,<br>30 total;<br>100x obj. | NA |
| 7d, 7e | Movie, CellASIC | 25 ms, 32%T mCherry;<br>25 ms, 32%T GFP | 1 μm/slice,<br>8 total;<br>60x obj. | 15 min/frame,<br>18h total |
| 8c | still images, live cells | 25 ms, 100%T TRITC;<br>25 ms, 100%T GFP | 0.3 μm/slice,<br>30 total; 100x obj. | NA |
| S5b | still images, live cells | 25 ms, 100%T TRITC | 1 μm/slice,<br>8 total; | NA |
|  |  |  | 60x obj. |  |
| S7b | still images,<br>fixed cells | 15 ms, 100%T mCherry;<br>50 ms, 100%T GFP;<br>10 ms, 100%T DAPI | 1 μm/slice,<br>8 total;<br>60x obj. | 10 min/frame,<br>18h total |

