## Supplementary Files S1-S4 (Scripts) for "Tom70-mediated mitochondria–nuclear envelope contacts regulate nuclear pore complex inheritance during gametogenesis": SupFile-1__MembraneProximityAnalysis_v6.m

```
% Triangle-to-Mesh Proximity Analysis
% --------------------------------------------------
% This script computes the shortest distance from each triangle in Object 1
% to the surface of Object 2, and exports area + centroid + distance data.
% It also reports what fraction of Object 1's surface lies within two
% user-defined proximity thresholds from Object 2.

% --- User-defined paths ---
objFile1 = '/Users/User1_name/Desktop/Temp_IN/cnm1/GUNC-ONLY_WT-Cell-1_Model-v1.obj';  % Object 1 (reference; e.g. nuclear envelope)
objFile2 = '/Users/User1_name/Desktop/Temp_IN/cnm1/Mito-ONLY_WT-Cell-1_Model-v1.obj';  % Object 2 (target; e.g. mitochondrion)
outputFile = '/Users/User1_name/Desktop/Temp_OUT/cnm1/GUNC-to-mito_WT-Cell-1_MembraneProximity_v6.txt';
figOutputFile = strrep(outputFile, '.txt', '_Preview.fig');  % Save .fig in same location


% --- User-defined Scaling Factor ---

% Optional user-defined per-instance scaling:
% scaleFactor = input('Enter isotropic scaling factor (e.g., 1.0 if no scaling): ');

scaleFactor = 2.2222;
if isempty(scaleFactor) || ~isnumeric(scaleFactor) || scaleFactor <= 0
    error('Invalid scaling factor.');
else
    fprintf('Corrected with an isotropic scaling factor of: %.4f\n', scaleFactor);
end


% --- checks for package containing point2mesh function ---
if exist('point2trimesh', 'file') ~= 2
    error('point2trimesh not found. Download it from File Exchange and add it to your MATLAB path.');
end

% --- Load meshes using robust reader ---
obj1 = readObj(objFile1);
obj2 = readObj(objFile2);

% --- Assign vertices ---
V1 = obj1.v;
V2 = obj2.v;
% --- Apply scaling to vertices ---
V1 = V1 * scaleFactor;
V2 = V2 * scaleFactor;


% --- Prepare Object 2 faces ---
if isstruct(obj2.f)
    face_cells = struct2cell(obj2.f);
    F2 = vertcat(face_cells{:});
else
    F2 = obj2.f;
end

% --- Sanity check for Object 2 ---
% Validate face indices do not exceed vertex count

if max(F2(:)) > size(V2, 1)
    error('Invalid mesh: max face index (%d) exceeds number of vertices (%d)', max(F2(:)), size(V2,1));
end

% --- Prepare Object 1 faces ---
if isstruct(obj1.f)
    face_cells = struct2cell(obj1.f);
    F1 = vertcat(face_cells{:});
else
    F1 = obj1.f;
end

% --- Sanity check for Object 1 ---
% Validate face indices do not exceed vertex count
if max(F1(:)) > size(V1, 1)
    error('Invalid mesh: max face index (%d) exceeds number of vertices (%d)', max(F1(:)), size(V1,1));
end

% --- Debug: Check mesh integrity ---
fprintf('Object 1 - Max face index: %d | Num vertices: %d\n', max(F1(:)), size(V1,1));
fprintf('Object 2 - Max face index: %d | Num vertices: %d\n', max(F2(:)), size(V2,1));

% --- Parameters ---
threshold1 = 30;  % First proximity threshold (nm)
threshold2 = 80;  % Second proximity threshold (nm)

% --- Visualization colors (customizable) ---
threshold1_color = [0.0 0.8 0.2];       % Color for distances < threshold1 (e.g. green)
threshold2_color = [0.5 0.7 1.0];       % Color for distances < threshold2 but >= threshold1 (e.g. light blue)
above_thresholds_color = [1.0 0.3 0.3]; % Color for distances >= threshold2 (e.g. red)

% --- Compute triangle distances and areas (mostly deleted after changing the measurement strategy from for loop to something else)---
num_tri = size(F1, 1);

% --- Compute triangle distances and areas ---

% Edge vectors for all triangles
vec1 = V1(F1(:,2), :) - V1(F1(:,1), :);
vec2 = V1(F1(:,3), :) - V1(F1(:,1), :);

% Areas using cross products
areas = 0.5 * vecnorm(cross(vec1, vec2, 2), 2, 2);

% Centroids
centroids = (V1(F1(:,1), :) + V1(F1(:,2), :) + V1(F1(:,3), :)) / 3;

% Comupte distance distance from all centroids (in chunks) to Object 2, with waitbar
% (chunking is to prevent RAM from being overwhelmed)
% 
chunk_size = 500;
num_centroids = size(centroids, 1);
distances = zeros(num_centroids, 1);

h = waitbar(0, 'Calculating distances...');

for idx = 1:chunk_size:num_centroids
    batch_idx = idx:min(idx+chunk_size-1, num_centroids);
    batch_centroids = centroids(batch_idx, :);

    % Query distances for this batch
    batch_distances = point2trimesh('Faces', F2, 'Vertices', V2, ...
                                    'QueryPoints', batch_centroids);

    distances(batch_idx) = batch_distances;

    % Update waitbar
    waitbar(batch_idx(end) / num_centroids, h);
end

close(h);  % Close waitbar when done

% --- Fix: use absolute values for distances ---
distances = abs(distances);

% --- Area-based summaries for both thresholds ---
total_area_1 = sum(areas);

within_mask_1 = distances < threshold1;
within_mask_2 = distances < threshold2;

area_within_1 = sum(areas(within_mask_1));
area_within_2 = sum(areas(within_mask_2));

percent_within_1 = 100 * area_within_1 / total_area_1;
percent_within_2 = 100 * area_within_2 / total_area_1;

% --- Print summary ---
fprintf('Total surface area in Object 1: %.2f\n', total_area_1);
fprintf('Area within %.1f units: %.2f (%.2f%%)\n', ...
    threshold1, area_within_1, percent_within_1);
fprintf('Area within %.1f units: %.2f (%.2f%%)\n', ...
    threshold2, area_within_2, percent_within_2);

% --- Export triangle-wise data to file (including vertices) ---
V1_indices = F1(:,1);
V2_indices = F1(:,2);
V3_indices = F1(:,3);

V1_coords = V1(V1_indices, :);
V2_coords = V1(V2_indices, :);
V3_coords = V1(V3_indices, :);

resultsTable = table((1:num_tri)', areas, distances, ...
    centroids(:,1), centroids(:,2), centroids(:,3), ...
    V1_coords(:,1), V1_coords(:,2), V1_coords(:,3), ...
    V2_coords(:,1), V2_coords(:,2), V2_coords(:,3), ...
    V3_coords(:,1), V3_coords(:,2), V3_coords(:,3), ...
    'VariableNames', {'TriangleIndex', 'Area', 'ShortestDistance', ...
                      'CentroidX', 'CentroidY', 'CentroidZ', ...
                      'V1x', 'V1y', 'V1z', ...
                      'V2x', 'V2y', 'V2z', ...
                      'V3x', 'V3y', 'V3z'});

writetable(resultsTable, outputFile, 'Delimiter', '\t');
fprintf('Saved triangle-wise data (with vertices) to: %s\n', outputFile);

% --- Visualization of proximity zones ---
figure; hold on;

% Zones:
% < threshold1      → threshold1_color
% < threshold2 only → threshold2_color
% ≥ threshold2      → above_thresholds_color

% Region 1: Within threshold1
patch('Faces', F1(within_mask_1,:), 'Vertices', V1, ...
      'FaceColor', threshold1_color, 'EdgeColor', 'none', 'FaceAlpha', 0.8);

% Region 2: Within threshold2 but not threshold1
in_thresh2_only = within_mask_2 & ~within_mask_1;
patch('Faces', F1(in_thresh2_only,:), 'Vertices', V1, ...
      'FaceColor', threshold2_color, 'EdgeColor', 'none', 'FaceAlpha', 0.5);

% Region 3: Beyond threshold2
beyond_thresh2 = ~within_mask_2;
patch('Faces', F1(beyond_thresh2,:), 'Vertices', V1, ...
      'FaceColor', above_thresholds_color, 'EdgeColor', 'none', 'FaceAlpha', 0.3);

% Overlay Object 2 mesh (semi-transparent)
patch('Faces', F2, 'Vertices', V2, ...
      'FaceColor', [0.6 0.6 1], 'EdgeColor', 'none', 'FaceAlpha', 0.1);

axis equal; view(3); camlight;
title(sprintf('< %.0f nm (%.1f%%), < %.0f nm (%.1f%%)', ...
              threshold1, percent_within_1, threshold2, percent_within_2));

% --------------------------------------------------
% --- FUNCTION: Robust OBJ reader ---
function obj = readObj(filename)
    fid = fopen(filename);
    if fid == -1
        error('Cannot open OBJ file: %s', filename);
    end

    v = {};
    f = {};

    while ~feof(fid)
        tline = strtrim(fgetl(fid));
        if isempty(tline) || startsWith(tline, '#')
            continue;
        end

        tokens = strsplit(tline);
        switch tokens{1}
            case 'v'
                v{end+1, 1} = str2double(tokens(2:4));

            case 'f'
                face = zeros(1, 3);
                for k = 1:3
                    parts = strsplit(tokens{1 + k}, '/');
                    face(k) = str2double(parts{1});  % just take the vertex index
                end
                f{end+1, 1} = face;
        end
    end

    fclose(fid);

    % Convert to matrices
    obj.v = cell2mat(v);
    obj.f = cell2mat(f);
end

% --- Compute total surface area of Object 2 ---
vecA2 = V2(F2(:,2), :) - V2(F2(:,1), :);
vecB2 = V2(F2(:,3), :) - V2(F2(:,1), :);
triangleAreas_obj2 = 0.5 * vecnorm(cross(vecA2, vecB2, 2), 2, 2);
total_area_2 = sum(triangleAreas_obj2);


% --- Compute Volume for Each Object ---
v1 = V1(F1(:,1), :); v2 = V1(F1(:,2), :); v3 = V1(F1(:,3), :);
signedVolumes1 = dot(v1, cross(v2, v3, 2), 2) / 6;
volume1 = abs(sum(signedVolumes1));

v1_2 = V2(F2(:,1), :); v2_2 = V2(F2(:,2), :); v3_2 = V2(F2(:,3), :);
signedVolumes2 = dot(v1_2, cross(v2_2, v3_2, 2), 2) / 6;
volume2 = abs(sum(signedVolumes2));

% --- Append summary data at end of output file ---
fid = fopen(outputFile, 'a');
fprintf(fid, '\n%% Summary Metrics (no per-triangle values)\n');
fprintf(fid, '%% TotalArea_Obj1\tTotalArea_Obj2\tArea_Within_T1\tArea_Within_T2\tPctArea_T1\tPctArea_T2\tVol_Obj1\tVol_Obj2\n');
fprintf(fid, '%.6f\t%.6f\t%.6f\t%.6f\t%.3f\t%.3f\t%.6f\t%.6f\n', ...
    total_area_1, total_area_2, area_within_1, area_within_2, ...
    percent_within_1, percent_within_2, volume1, volume2);
fclose(fid);

% --- Save interactive preview figure ---
savefig(figOutputFile);
```
