## Supplementary Files S1-S4 (Scripts) for "Tom70-mediated mitochondria–nuclear envelope contacts regulate nuclear pore complex inheritance during gametogenesis": SupFile-2__Apply-C1-Threshold_Htb1-mCherry-Masks_v2.ijm

```
//	Htb1-mCherry Masking (v2)
//	-------------------------
//
//	Generates a binary mask from Htb1-mCherry signal (C1), using a user-defined thresholding method, then adds this mask as a third channel the starting 16-bit 2-channel hyperstack.

input = "/Users/User1_name/Desktop/Temp_IN"
output ="/Users/User1_name/Desktop/Temp_OUT"

threshold_method = getString("Desired thresholding method:", "Otsu");
contrast_min = getNumber("Min, contrast value:", 100);
contrast_max = getNumber("Max, contrast value:", 4500);

file_list = getFileList(input);
for (i=0; i<file_list.length; i++) {
		current_file = file_list[i];
		run("Bio-Formats (Windowless)", "open=" + input + "/" + current_file + " color_mode=Composite view=Hyperstack stack_order=XYCZT use_virtual_stack");
		selectImage(current_file);
		run("Split Channels");
		
//	If using a 3-channel hyperstack (e.g. with merged brightfield),consider removing using the the following commented block to remove it;
//	this additional channel can drastically slow down image handling:
//
//		selectImage("C3-" + current_file);
//		close();
		
		selectImage("C1-" + current_file);
		setMinAndMax(contrast_min, contrast_max);
		
		setAutoThreshold("Otsu dark no-reset");
		setOption("BlackBackground", true);
		run("Convert to Mask", "method=Otsu background=Dark calculate black create");

		selectImage("MASK_C1-" + current_file);
		run("16-bit");

		run("Merge Channels...", "c1=C1-" + current_file + " c2=C2-" + current_file + " c3=MASK_C1-" + current_file + " create");
		selectImage(current_file);
		Stack.setDisplayMode("composite");
		Stack.setChannel(3);
		run("Grays");
		
		selectImage(current_file);
		saveAs("Tiff", output + "/Masked_" + current_file);

run("Close All");

}

run("Close All");
```
