## Supplementary Files S1-S4 (Scripts) for "Tom70-mediated mitochondria–nuclear envelope contacts regulate nuclear pore complex inheritance during gametogenesis": SupFile-3__NUP-Inheritance-Index_Quantification_v4.ijm

```
//	NUP Inheritance Index measurement (v4)
//	--------------------------------------
//
//	Generates and quantified C2 signal in a nuclear envelope (NE) banded ring ROI around chromatin signal of user-defined nuclei (marked by masks of Htb1-mCherry signal).
//	User must define and save appropriate nucleoplasmic ROIs (time/frame and z-slice of midsection, for cells forming four intact spores/PSMs), beforehand.
//	ImageJ/FIJI measurement preferences should be set to include at least the following measurements: mean intensity, median intensity, area, frame (time), slice, and channel.
//
//	Set ROIs and add to ROI Manager with [t]:
//	-	For each cell properly forming four spores (or four persisting intact PSMs), select the time frame and Z-slice for a predivision "Progenitor" nuclei and the four daughter "Spore" nuclei.
//		NOTE: Choose (1) a "Progenitor" nuclear ROI, then choose (2-5) "Spore" nuclear ROIs for each gamete produced withing the progenitor.
//			  The ROI order is extrememely important, as proper measurement and downstream analysis assume five ROIs per cell in this order.
//		-	Use the Wand Tool on the Htb1-mCherry masks (chanel 3)
//		-	Select the latest pre-division frame lacking a nuclear flare and having relatively normal morphology (i.e. somewhat round) and minimal/tolerable chromatin signal exlcusion at the periphery from the nucleolus.
//		-	Select the earliest post-closure frame, defining closure as the redistribution/repositioning of nuclei following MII and the rounding of daughter nuclei (vs teardrop-shaped) as PSMs round.
//			-	Use the midsection z-slice for nuclei (i.e. with the greatest circumference and best apparent focus for chromatin structures/texture).
//			-	If desired nuclear crossection has imporper mask (e.g. merged with mask of other nucleus), use ellipse or polygon tool to manually outline chromatin signal, or skip cell.
//
//	Analysis Notes:
//	-	For subsequent analysis, compare measurements of "Spore"s to their respective "Progenitor", e.g. by dividing the average of the four "Spore"'s mean intensity by that of the "Progenitor".
//	- 	For quality control, assess whether there are differences between compared groups (e.g. mutants, treatments, etc) in area of both "Progenitor" and "Spore" NE, frame (time) of "Spore" measurements,
//	  	and the delta of "Spore"-to-"Progenitor" frames (time) and slices. These assessments can reveal potentially confounding differences in NE size/shape, meiotic entry (which can result in inter-genotype differences in photobleaching and singal quality),
//		duration of meiotic divisions, and z-drift (which could lead to inter-genotype differences in signal quality).


input = "/Users/User1_name/Desktop/Temp_IN";
ROI_input = "/Users/User1_name/Desktop/Temp_ROI";
output ="/Users/User1_name/Desktop/Temp_OUT";
sub_background = getString("Enter desired subtract background rolling pxl radius:", "50");
measure_channel = getNumber("Select desired channel for measurement:", 2);

//loop through directory
file_list = getFileList(input);
for (i=0; i<file_list.length; i++) {
	current_file = file_list[i];
	file_length = lengthOf(current_file);
	file_stem = substring(current_file, 0, file_length-4);	// Removes ".tif" file ending from stored image name.
	print("Now processing: "+ current_file);
	
	run("Bio-Formats (Windowless)", "open=" + input + "/" + current_file + " color_mode=Composite view=Hyperstack stack_order=XYCZT use_virtual_stack");
	selectWindow(current_file);
	
	run("Split Channels");
	close("C3-" + current_file);	// Removes now-unneccesary Htb1-mCherry mask channel. 
	run("Merge Channels...", "c1=C1-" + current_file + " c2=C2-" + current_file + " create");
	selectWindow(current_file);

	Stack.setChannel(measure_channel);
	print("Subtracting Background: " + current_file);
	run("Subtract Background...", "rolling=" + sub_background + " stack");
	
	roiManager("Open", ROI_input + "/RoiSet_" + file_stem + ".zip");
	n = roiManager('count');
		for (j = 0; j < n; j++) {
 		roiManager('select', j);
			if (j % 5 != 0) {
			run("Enlarge...", "enlarge=-1 pixel");
			print("ROI #" + j+1 + " is classified as a spore nuclei, per ROI order.");
			run("Make Band...", "band=0.3");
			}
			else {
			run("Enlarge...", "enlarge=-1 pixel");
			print("ROI #" + j+1 + " is classified as a progenitor nuclei, per ROI order.");
 			run("Make Band...", "band=0.4");	// Note: predivision NUP signal occupies a thicker band than of spores (presumably due to uncreased abundance). 
			}
		Stack.setChannel(measure_channel);
		run("Measure");
	
		}

	selectWindow("Results");
	saveAs("txt", output + "/" + "v4_NUP-Inheritance-Quantification_" + current_file);

	run("Close All");
	run("Fresh Start");

	}
	
run("Close All");
```
