## Supplementary Files S1-S4 (Scripts) for "Tom70-mediated mitochondria–nuclear envelope contacts regulate nuclear pore complex inheritance during gametogenesis": SupFile-4__Measure_Largest-Object_Approximate-Area_v4.ijm

```
//	Largest Particle Size-Approximation (v4)
//	----------------------------------------
//
//	This is script is for determining GEM or aggregate particle size (or approximating, given limitations inherent to partice size and the diffraction limit).
//	A binary threshold is used to detect particles, then the ROI of the largest thresholded particle is measured. Designed for use on MAX- or SUM z-projected 16-bit 2-channel hyperstacks.
//	ImageJ/FIJI measurement preferences should be set to include at least the following measurements: area, mean intensity, median intensity, and channel. Frame (time) information is automatically saved by the script for each measurement.

//	The MorphoLibJ plugin (see IJPB-plugins) is required for choosing the largest thresholded object.	
//	Requires user-defined ROIs (e.g. drawn with the rectangle tool) to specify the desired region and frame (time) for each cell measured. 
//
//	For quality control, confirm there are no appreciable differences between compared groups (e.g. genotypes) for the frame (time) measured, as differential photobleaching may confound results.


input = "/Users/User1_name/Temp_IN";
ROI_input = "/Users/User1_name/Temp_ROI";
output = "/Users/User1_name/Temp_OUT";

waitForUser("Currently set to measure particles only in channel 2");
//measure_channel = getNumber("Select desired channel for measurement:", 2);
sub_background = getString("Enter desired subtract background rolling pxl radius:", "50");
contrast_min = getNumber("Min, contrast value:", 90);
contrast_max = getNumber("Max, contrast value:", 10000);

Min_string = toString(contrast_min);
Max_string = toString(contrast_max);

//loop through directory
file_list = getFileList(input);
for (i=0; i<file_list.length; i++) {
	current_file = file_list[i];
	file_length = lengthOf(current_file);
	file_stem = substring(current_file, 0, file_length-4);	//removes ending ".tif" from filename
	print("Now processing: "+ current_file);
	
	run("Bio-Formats (Windowless)", "open=" + input + "/" + current_file + " color_mode=Grayscale view=Hyperstack stack_order=XYCZT use_virtual_stack");
	selectWindow(current_file);

	print("Subtracting Background: " + current_file);
	run("Subtract Background...", "rolling=" + sub_background + " stack");
	
	Stack.setChannel(2);
	setMinAndMax(contrast_min, contrast_max);
		
	roiManager("Open", ROI_input + "/RoiSet_" + file_stem + ".zip");
	n = roiManager('count');
// 	The following array is for properly tracking the frames (time) used, which is lost from the image metadata during intermediate processing.
	ROI_timepoint_tracking = newArray(n);
		for (j = 0; j < n; j++) {
 		selectWindow(current_file);
 		roiManager('select', j);
 		Stack.getPosition(channel, slice, frame);
 		ROI_timepoint_tracking[j] = frame;
 		Stack.setChannel(2);
	    run("Duplicate...", "title=CurrentROI duplicate frames=" + frame);

 		selectImage("CurrentROI");
 		run("Split Channels");
 		selectImage("C1-CurrentROI");
		close;
		
 		selectImage("C2-CurrentROI");
 		run("Duplicate...", "title=C2-CurrentROI_copy duplicate");
 		selectImage("C2-CurrentROI_copy");
		setAutoThreshold("Otsu dark no-reset");
		setOption("BlackBackground", true);
		run("Convert to Mask", "method=Otsu background=Dark calculate black create");
 		
 		selectImage("C2-CurrentROI_copy");
 		
//		MorphoLibJ plugin is required (IJPB-plugins) for following command:		
		run("Keep Largest Region");
		selectImage("C2-CurrentROI_copy-largest");
		run("Create Selection");
		
		selectImage("C2-CurrentROI");
		run("Restore Selection");
		run("Measure");
		
		close;
		selectImage("C2-CurrentROI_copy-largest");
		close;
		selectImage("C2-CurrentROI_copy");
		close;
	}
	
	for (k = 0; k < ROI_timepoint_tracking.length; k++) {
    setResult("ROI_timepoint_tracking", k, ROI_timepoint_tracking[k]);
}	
	
	selectWindow("Results");
	saveAs("txt", output + "/" + "v4_GEM-size-quant_Min" + Min_string + "Max" + Max_string + "_" + current_file);
	
	ROI_timepoint_tracking = newArray(ROI_timepoint_tracking.length); 
	run("Close All");
	run("Fresh Start");

}

	run("Close All");
	run("Fresh Start");
```
